# Efficient Representations of Tumor Diversity with Paired DNA-RNA Aberrations

**DOI:** 10.1101/2020.04.24.060129

**Authors:** Qian Ke, Wikum Dinalankara, Laurent Younes, Donald Geman, Luigi Marchionni

**Affiliations:** Department of Applied Mathematics and Statistics, Johns Hopkins University, Baltimore, MD, USA; Department of Oncology, Johns Hopkins University School of Medicine, Baltimore, MD, USA; Department of Pathology and Laboratory Medicine, Weill Cornell Medicine, New York, NY, USA

## Abstract

Cancer cells display massive dysregulation of key regulatory pathways due to now well-catalogued mutations and other DNA-related aberrations. Moreover, enormous heterogeneity has been commonly observed in the identity, frequency and location of these aberrations across individuals with the same cancer type or subtype, and this variation naturally propagates to the transcriptome, resulting in myriad types of dysregulated gene expression programs. Many have argued that a more integrative and quantitative analysis of heterogeneity of DNA and RNA molecular profiles may be necessary for designing more systematic explorations of alternative therapies and improving predictive accuracy.

We introduce a representation of multi-*omics* profiles which is sufficiently rich to account for observed heterogeneity and support the construction of quantitative, integrated, metrics of variation. Starting from the network of interactions existing in Reactome, we build a library of “paired DNA-RNA aberrations” that represent prototypical and recurrent patterns of dysregulation in cancer; each two-gene “Source-Target Pair” (STP) consists of a “source” regulatory gene and a “target” gene whose expression is plausibly “controlled” by the source gene. The STP is then “aberrant” in a joint DNA-RNA profile if the source gene is DNA-aberrant (*e.g.*, mutated, deleted, or duplicated), and the downstream target gene is “RNA-aberrant”, meaning its expression level is outside the normal, baseline range. With *M* STPs, each sample profile has exactly one of the 2*^M^* possible configurations.

We concentrate on subsets of STPs, and the corresponding reduced configurations, by selecting tissue-dependent minimal coverings, defined as the smallest family of STPs with the property that every sample in the considered population displays at least one aberrant STP within that family. These minimal coverings can be computed with integer programming. Given such a covering, a natural measure of cross-sample diversity is the extent to which the particular aberrant STPs composing a covering vary from sample to sample; this variability is captured by the entropy of the distribution over configurations.

We apply this program to data from TCGA for six distinct tumor types (breast, prostate, lung, colon, liver, and kidney cancer). This enables an efficient simplification of the complex landscape observed in cancer populations, resulting in the identification of novel signatures of molecular alterations which are not detected with frequency-based criteria. Estimates of cancer heterogeneity across tumor phenotypes reveals a stable pattern: entropy increases with disease severity. This framework is then well-suited to accommodate the expanding complexity of cancer genomes and epigenomes emerging from large consortia projects.

**Author Summary:** A large variety of genomic and transcriptomic aberrations are observed in cancer cells, and their identity, location, and frequency can be highly indicative of the particular subtype or molecular phenotype, and thereby inform treatment options. However, elucidating this association between sets of aberrations and subtypes of cancer is severely impeded by considerable diversity in the set of aberrations across samples from the same population. Most attempts at analyzing tumor heterogeneity have dealt with either the genome or transcriptome in isolation. Here we present a novel, multi-omics approach for quantifying heterogeneity by determining a small set of paired DNA-RNA aberrations that incorporates potential downstream effects on gene expression. We apply integer programming to identify a small set of paired aberrations such that at least one among them is present in every sample of a given cancer population. The resulting “coverings” are analyzed for six cancer cohorts from the Cancer Genome Atlas, and facilitate introducing an information-theoretic measure of heterogeneity. Our results identify many known facets of tumorigenesis as well as suggest potential novel genes and interactions of interest.

**Data Availability Statement:** RNA-Seq data, somatic mutation data and copy number data for The Cancer Genome Atlas were obtained through the Xena Cancer Genome Browser database (https://xenabrowser.net) from individual cancer type cohorts. Computational functionality for the optimization procedure is provided at https://github.com/wikum/lpcover and the code for the analysis in the manuscript is provided at https://github.com/wikum/CoveringAnalysis. Processed data in the form of TAB delimited files, and selected tissue-level coverings (in excel format) are provided as additional supplementary material and are also available from the Marchionni laboratory website (http://marchionnilab.org/signatures.html)

## 1 Introduction

Cancer cells evade the normal mechanisms controlling cellular growth and tissue homeostasis through the disruption of key regulatory pathways controlling these processes. Such dysregulation results from genetic and epigenetic aberrations, encompassing mutations, copy number alterations, and changes in chromatin states, which affect the genes participating in such regulatory networks.

Over the past several decades, the list of known genetic and genomic aberrations in cancer has greatly expanded, thanks to large-scale projects such as the The Cancer Genome Atlas (TCGA, (Cancer Genome Atlas Research Network et al., 2013), the Catalogue Of Somatic Mutations In Cancer (COSMIC, Tate et al. (2019)), the MSK/IMPACT study (Zehir et al., 2017), and recents efforts from the ICGC/TCGA Pan-Cancer Analysis of Whole Genomes Consortium (ICGC/TCGA Pan-Cancer Analysis of Whole Genomes Consortium, 2020).

Whereas the number of aberrations which suffice for progression to an advanced cancer is thought to be rather small, at least for solid tumors (Vogelstein and Kinzler, 2015; Tomasetti et al., 2015) and at the pathway level (Sever and Brugge, 2015), the number of ways (combinations of aberrations) for which this can be actualized is very large. In particular, the landscape collectively emerging from these studies exhibits a high degree of variation in the identity, frequency, and location of these aberrations, as well as tissue- and expression-dependency (Haigis et al., 2019; Lawrence et al., 2013). These differences—collectively referred to as *tumor heterogeneity*—are “context-specific”, differing among tissue types and epigenetic conditions (Haigis et al., 2019), across different cells within a lesion (*intra-tumor heterogeneity*), between tumor lesions within the same individual (*inter-tumor heterogeneity*), and across distinct individuals with the same cancer type or sub-type (*across-sample or population-level heterogeneity*).

In addition, such DNA defects, in order to be “functional” (*i.e.*, manifest themselves) and ultimately alter the cellular phenotype, must propagate through the signaling and regulatory network and alter the downstream gene expression programs (Osmanbeyoglu et al., 2017; PCAWG Transcriptome Core Group et al., 2020). These downstream transcriptional changes are in fact also context-specific, varying within and among cancers, local environments, and individuals. Most importantly, it has been speculated that transcriptionally heterogeneous tumors may be more adaptable to changes in the tumor microenvironment and therefore more likely to acquire new properties such as metastatic potential and resistance to treatments, leading to dismal patient outcomes; in addition, for predicting the response to targeted therapies, gene expression profiles may be more discriminating than mutational status (Costello et al., 2014). The analysis of heterogeneity of molecular profiles, both DNA and RNA, is therefore of paramount importance. Consequently, a deeper, integrative and quantitative analysis of tumor heterogeneity is necessary for achieving a better understanding of the underlying biology, for designing more systematic explorations of candidate therapies, and for improving the accuracy of prognosis and treatment response predictions.

Unsurprisingly, even *representing* such high-dimensional variability poses great challenges, especially if a major goal is to find suitable metrics to quantify the level of tumor heterogeneity. We assume that large-scale projects (see above) and studies (*e.g.*, Bailey et al. (2018)) have already provided reasonably comprehensive lists of the most important recurrent molecular alterations driving cancer initiation and progression. But merely counting or cataloging aberrations will not suffice to precisely measure heterogeneity in a tumor population, and to quantify how this differs across diverse contexts (*e.g.*, between cancer arising in distinct organs, or between tumor sub-types). In order to identify functional aberrations potentially exploitable as biomarkers and therapeutic targets, it is necessary to go well beyond frequency estimates to more powerful representations rooted in biological mechanism and accounting for statistical dependency among aberrations.

We introduce a representation of *omics* profiles which is sufficiently rich to account for observed heterogeneity and to support the construction of quantitative, integrated metrics. Our framework is centered on the joint analysis of “paired DNA-RNA aberrations” that represent prototypical and recurrent patterns of dysregulation in cancer. Specifically, we represent the space of gene alterations that result in network perturbations and downstream changes of gene expression levels as a catalogue of mechanistic, two-gene “ Source-Target Pairs” (STPs), each consisting of a “source” gene (important driver) and a “target” gene for which the mRNA expression is controlled either directly by the source gene or indirectly by a close descendant of the source.

We extend STPs from a network property to a sample property (like the existence of individual aberrations) by declaring an STP to be “aberrant” in a joint DNA-RNA profile if the source gene is DNA-aberrant (*e.g.*, mutated, deleted, or duplicated), and the target gene is RNA-aberrant, meaning its expression level is “divergent” (*i.e.*, outside the normal, baseline range (Dinalankara et al., 2018)). This defines one binary random variable per STP, of which there are typically hundreds of thousands, most of which have a very small probability to be realized in a sample.

Samples are then characterized by their entire set of paired DNA-RNA aberrations, or aberrant STPs. Therefore, given there are *M* STPs, exactly one of the 2^*M*^ possible configurations is assigned to each sample. The extent to which these subsets vary from sample to sample is then a natural measure of heterogeneity in the population from which the samples are drawn.

Due to the difficulty of estimating rare events with the modest sample sizes available in cancer genomics today, any multivariate property of the probability distribution over the 2^*M*^ STP configurations (for example, its entropy) cannot be accurately approximated without a substantial further reduction of complexity. Such a reduction is provided by the concept of *minimal coverings* of a population (previously employed for modeling networks (Hristov and Singh, 2017)). Here, we focus on smallest collections *C* of paired aberrations with the property that (nearly) every tumor sample has at least one aberrant STP in *C*. Indeed, since nearly all tumor samples exhibit multiple aberrant STPs, a *minimal covering* necessarily exists (perhaps not unique), which can be found using well-known algorithms for formulating “optimal set covering” as the solution of an integer-programming problem (see Methods).

Our main contribution is then a method for integrating DNA and RNA data which yields novel insights about regulatory mechanisms in cancer, and consists of three parts:

1. A representation of network dysregulation based on matched pairs of genes, one gene with aberrant DNA and the other gene downstream, with aberrant RNA expression.
2. An algorithm for finding the minimal covering of a cancer (sub)population by aberrant genes or gene pairs.
3. An information-theoretic characterization of inter-sample heterogeneity as the entropy of the distribution of covering states.

Our methods are described in more detail in the next sections, followed by a presentation of our results. We conclude this paper with a discussion and provide additional results in supplementary material.

## 2 Methods

### 2.1 Overall Strategy

Identifying and quantifying the cross-sample heterogeneity of *omics* datasets with large numbers of random variables requires making simplifying assumptions and approximations on the joint distribution of the considered variables to make it feasible. We performed our analyses using matched DNA mutations, copy number alterations, and RNA expression data, pre-processed with the method previously described in (Dinalankara et al., 2018). In the present study we specifically focused on six distinct tumor types (TCGA code in parenthesis): breast invasive carcinoma (BRCA), prostate adenocarcinoma (PRAD), lung adenocarcinoma (LUAD), liver hepatocellular carcinoma (LIHC), kidney renal clear cell carcinoma (KIRC), and colon adenocarcinoma (COAD). For simplicity, hereafter, we will refer to these tumor types according to the tissue of origin (breast, prostate, lung, kidney, liver, and colon cancer).

Our definition of aberrant expression of RNA (Dinalankara et al., 2018) requires expression data from a baseline population, taken here as corresponding normal tissue (see 2.2.1). Consequently, our selection of cancer types was constrained by having enough normal samples in TCGA to estimate the “normal expression range” of the RNA-Seq data. In addition, we also consider a variety of clinical scenarios across different patient populations. Our approach is depicted in the schematic of Figure 1.

**Figure 1:**
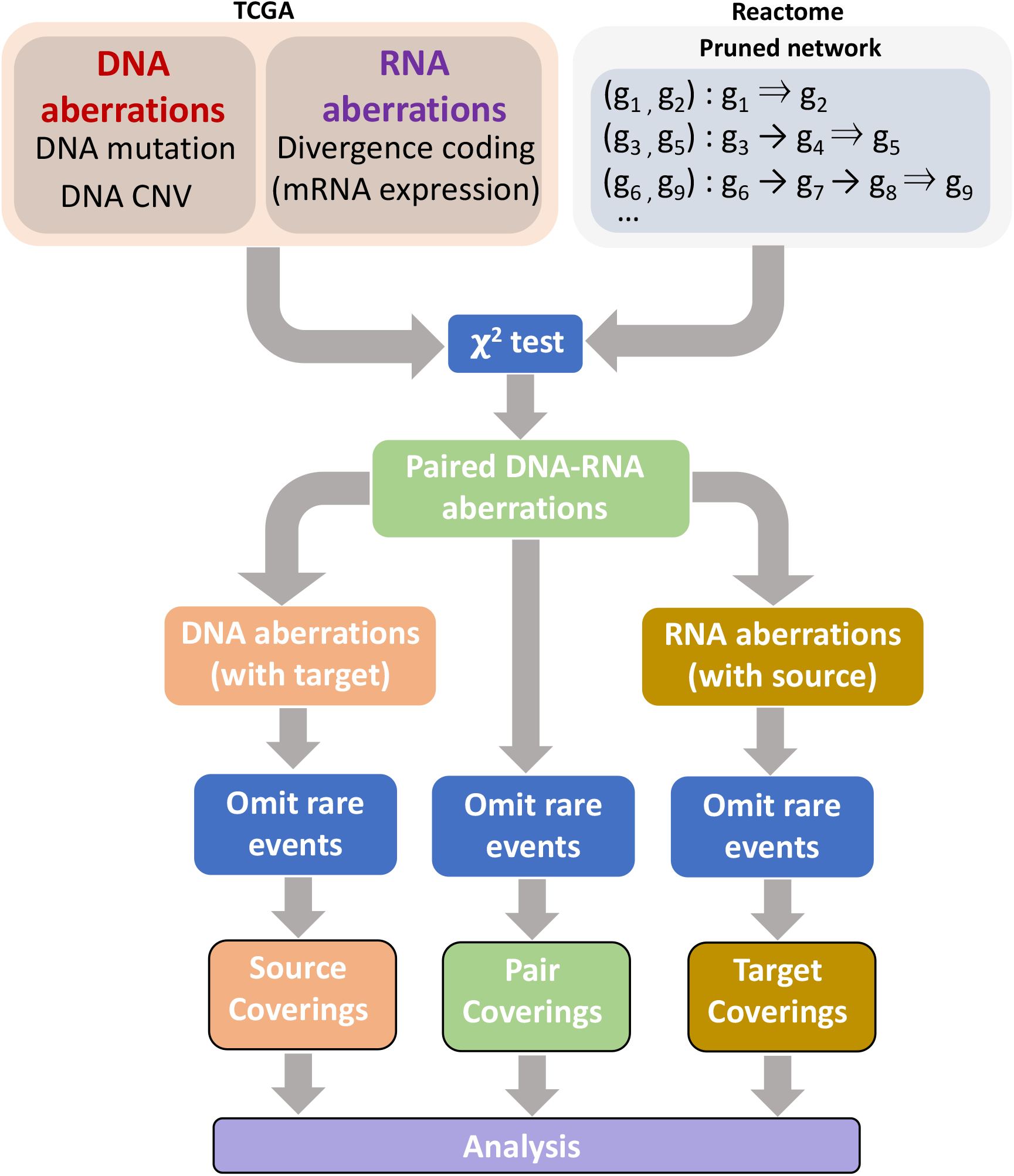
Overall analytical workflow. Source-target pairs (STPs) are constructed using the links available in Reactome (Jassal et al., 2020). In the TCGA cancer cohorts, the mutation and copy number variation data are used to construct binary DNA aberration profiles; the presence of either a mutation or high/low copy number variation at a given gene is treated as an aberration for the given gene for that sample *omics* profile. The gene expression data are used to construct binary RNA aberration profiles based on falling outside the “normal” expression range (in quantiles) for each gene based on TCGA normal tissue expression data, as previously described (Dinalankara et al., 2018). The binary profiles are combined to produce paired DNA-RNA aberrations, following which filtering is performed by selecting pairs that are determined to be significant (two-sided *χ*^2^ test). The selected STPs then give rise to individual source (DNA) and target (RNA) aberrations, providing binary *omics* profiles at the level of source, target, and pairs. STPs that are present in less than 2% of samples for a given tissue are omitted. Then coverings are computed at the pair, source and target levels and subtype analysis and heterogeneity analysis carried out.

#### Cancer phenotypes

We also focused on specific patient subgroups defined based on standard clinical and pathological variables (which are ordinal) routinely used for patient risk stratification. Tumor stage (from I to IV) indicates extension of a cancer and whether it has spread beyond the site of origin. The lymph node status (positive versus negative) indicates the presence of lymph node metastases. Tumor T status (from T1 to T4) indicates the size of the primary tumor. Tumor histologic grade (from G1 to G3 or G4, depending on the tumor type) captures the progressive departure from the the normal tissue and cellular architecture observed under a microscope. The Gleason grading system (Humphrey, 2004) is specific to prostate cancer and it accounts for 5 grades. The Gleason sum results from the two predominant grade patterns observed (*i.e.*, “primary” and “secondary”), with a sum of 6 (3+3) corresponding to indolent tumors, and sums from 7 to 10 associated with increasingly aggressive phenotypes. Finally, the PAM50 breast cancer subtypes (Parker et al., 2009) and the colorectal cancer CRIS classes (Isella et al., 2017) are patient subgroups with distinct prognosis defined based on specific gene expression signatures.

#### Aberration detection

We reduce the data to binary variables indicating deviations from normal behavior, and the resulting indicators are furthermore filtered using an STP-based analysis requiring plausible mechanisms leading to the aberrations.

#### Covering estimation

In order to reduce the number of variables under consideration, we estimate subsets of “important variables,” called coverings, defined as minimal sets of variables from which, with high probability, cancer samples have at least one aberrant observation (see Section 2.3 on their computation). Because such optimal coverings are generally not unique, we include the consideration of variables that are present in at least one covering (union), or the restriction to variables that appear in all of them (intersection), that we refer to as “core” variables, or the use of a single covering, for example the one maximizing the sum of frequencies of aberrations among its variables.

#### Entropy estimation

We assess the heterogeneity of a population of samples by computing the entropy over a limited family of configurations determined by a covering of this population. This computation is not straightforward; even though reduced profiles involve a relatively small number *m* of binary variables (typically a few dozen) indicating aberration of STPs, the observed sample size remains insufficient to allow for the estimation of the probabilities of the 2^*m*^ joint configurations of these variables. Some approximations are necessary and are described in Section 2.4.

#### Code-based reduction

Using a tree-based decomposition, we decompose tumor samples resulting from a given cancer type into cells, or bins, associated with a small number of conjunctions and disjunctions of aberrations. It is then possible to visualize and compare the resulting histograms in sub-populations defined by specific subtypes or phenotypes. This is described in Section 2.5.

### 2.2 Aberrations

#### 2.2.1 Univariate deviation from normality in *omics* data

We transform the original data into sparse binary vectors indicating whether each variable deviates from a reference state or normal range when observed on a given sample.

Our pre-processing of DNA data, which already provides deviations from wild type, is quite simple. We consider that a gene *g* is aberrant at the DNA level if it includes a mutation that differs from the wild type, or if its copy number corresponds to a homozygous deletion (which would entail a complete gene inactivation), or a gain of 2 copies (to increase the plausibility of aberrant over-expression). We exclude heterozygous deletions and single copy gains since their impact is more difficult to interpret biologically. We will write 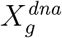 for the corresponding binary random variable, so 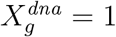 when *g* is DNA-aberrant.

The binarization of RNA data is more involved, and is based on the notion of “divergence” we previously developed (Dinalankara et al., 2018). Briefly, following a rank transformation, the range of RNA expression is estimated for normal samples for genes of interest. Then for each tumor sample and each gene, there is a binary variable with values 1 or 0 depending on whether the expression of the gene is outside or inside the expected normal region. Thus a gene is declared as RNA-aberrant if its ranking among other genes in the same sample falls outside of its normal range estimated from baseline data. Let 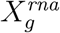 be the corresponding binary random variable. This dichotomization requires a training step, solely based on normal tissue data, in order to estimate these normal intervals of variation. This being done, the decision for a gene to be RNA-divergent in a tumor sample only involves the RNA profile of this sample and is in particular independent of other tumor observations in the dataset.

#### 2.2.2 Building source-target pairs

These binary *omics* variables are filtered by requiring that the deviations they represent have a plausible causal explanation as parts of STPs. Such STPs, denoted (*g_s_* ⇒ *g_t_*), are built using apriori information representing known gene-gene interactions from signaling pathways and biological processes. In our implementation, we used the Reactome database (Jassal et al., 2020) as retrieved from Pathway Commons (version 10) (Cerami et al., 2011), since the network information contained therein is comprehensive and well curated (see supplementary text for further details on network curation and summary statistics). Our approach may be implemented using other databases providing gene-gene interaction information to build STPsas well…

Let 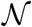 denote the family of directed pairs of genes from this database, annotated as regulator and target, including two kinds of links *g* → *g′* for which “*g* controls state change of *g′*” (notation: 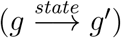 or “*g* controls the expression of *g′*” (notation 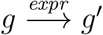). We say that two genes *g_s_*, *g_t_* form a “*source-target pair (STP)*”, with notation *g_s_* ⇒ *g_t_* if there exists a sequence of *l* intermediate genes *g_1_*, …, *g_l_* such that

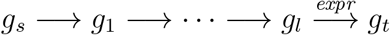

where the intermediate links are either 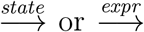 and the last link is 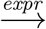. Such a sequence has *k* = *l* + 1 links, and the minimal number of links required to achieve the STP is called the length of *g_s_* ⇒ *g_t_*.

Let 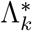 denote the set of STPs of length *k* or less deduced from the pathway database. This set, which is tissue independent, includes a large number of pairs (more than 200,000 for *k* = 3). For computational efficiency, we take *k* = 3 in our experiments. For a more detailed analysis on the selection of *k* = 3, see Supplementary text. Let 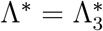 from here on. To select pairs that are most relevant for a tissue, this set is reduced by applying a *χ*^2^ test for independence, only keeping STPs (*g_s_* ⇒ *g_t_*) for which the independence between the events “*g_s_* DNA aberrant” and “*g_t_* RNA aberrant” is rejected at a 5% level by the test (without correction for multiple hypotheses, because we want to be conservative with this selection) using a dataset of tumor samples. Let Λ denote the set of remaining pairs (typically 5,000–10,000), which is therefore tissue dependent (see Figure 1).

We then let 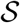 denote the set of sources in Λ, *i.e.*, the set of genes *g* such that there exists *g*′ with (*g* ⇒ *g*′ ∊ Λ) and, similarly, let 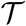 be the set of all possible targets. We let *T* (*g*) denote the set of all the targets of 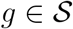, that is, 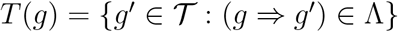 the set of all sources pointing to 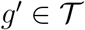.

#### 2.2.3 Paired aberrations

We can now define a family of binary random variables (*Z_λ_*, *λ* ∈ Λ) of “Paired Aberrations” with *Z_λ_* = 1 for STP *λ* = (*g_s_* ⇒ *g_t_*) if and only if *g_s_* is aberrant at the DNA level (either due to mutation or copy-number variation) and *g_t_* is RNA-aberrant. That is 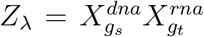 a product of binary variables. For *λ* = (*g_s_* ⇒ *g_t_*), we will also use the notation *s*(*λ*) = *g_s_* and *t*(*λ*) = *g_t_* for the source and target in *λ*.

From this, we also define binary variables 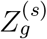 indicating aberrations at the source level letting 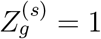 if and only if *g* participates in an aberrant motif as a source gene. Therefore,

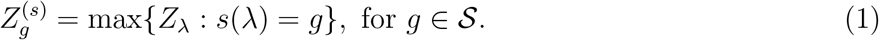

Similarly, we consider aberrations at the target level letting

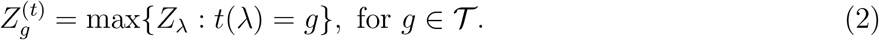

We will refer to the event 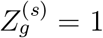 as a “source aberration with target” for gene *g* and the event 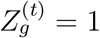 as an “target aberration with source” for gene *g*.

### 2.3 Coverings

#### 2.3.1 Definition

We have defined three types of aberrations involving multiple genes (STP, source, target), indexed by three different sets of pairs, sources or targets: i) STP aberrations which involve one source and one target gene; ii) “source aberration with target” which involve one source gene and all its targets; iii) “target aberrations with source” which involve one target gene and all its sources. We will use the generic notation 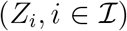 to refer to the variables associated to any one of them, so that 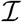 is one of the index sets Λ, 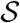 or 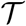 and *Z_i_* = *Z_λ_*, 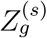 or 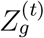 respectively. We identify small subsets of 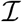 of “important variables” for describing the stochastic behavior of *Z*, such that at least one aberration occurs with high probability.

Denote by (Ω, *P*) the probability space on which 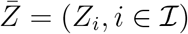 is defined. If *α* ∈ [0, 1] and *J* is a subset of 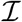, we will say that 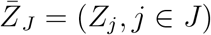 is a covering (or 1-covering) of Ω at level *α* if, with probability larger than 1 − *α*, at least one of its variables is aberrant, *i.e.,*

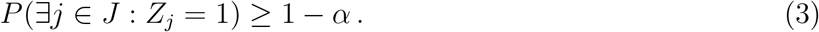

In other terms, Ω is covered (up to a subset of probability less than *α*) by the union of events *U_j_* = {*Z_j_* = 1}, *j* ∈ *J*. For simplicity, we will also refer to the index set, *J*, as a covering rather than the set of variables it indexes. More generally, we can define an *r*-covering at level *α* as a set *J* for which at least *r* of the variables in *J* are aberrant with probability 1 − *α*, *i.e.,*

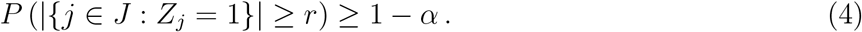

#### 2.3.2 Optimal coverings

We assume that a family of weights 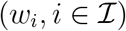 is given and consider the function

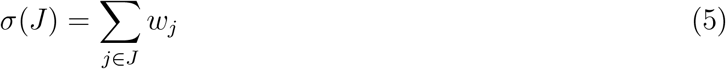

representing the weighted size of *J*. (Although, in our experiments, we use *w_j_* = 1 for all *j*, in which case *σ*(*J*) is just the number of elements in *J*, we present a weighted version of the problem, which can be useful in some situations.) We define a minimal covering as any covering minimizing *σ* among all other coverings.

To rephrase this as an integer programming problem, we note that the subsets *J* of 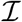 are in one-to-one correspondence with the set of all configurations 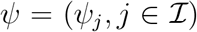, where *Ψ_j_* = 1 if *j* ∈ *J* and 0 otherwise. The minimal covering problem can then be reformulated as minimizing 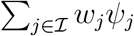 subject to the existence of a random variable *Y*: Ω → {0, 1} such that *P*(*Y* = 1) ≥ 1 − *α* and

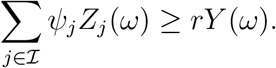

We have a finite sample of the distribution *P*, represented by a finite subset 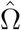 of Ω. We can approximate the covering problem by enforcing the constraints only for 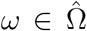 and replacing *P*(*Y* = 1) ≥ 1 − *α* by a sample fraction over 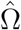. We then determine a minimal *r*-covering at level *α* by minimizing

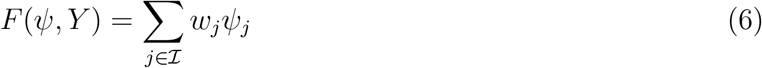

subject to the constraints

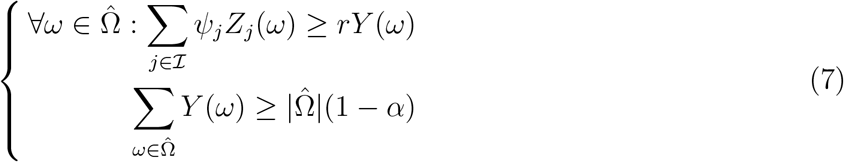

Many optimal solutions are equal *w_j_*. Assume that one obtains several such sets *J* ^(1)^, …, *J*^(*N*)^, all with same cardinality, and all providing coverings at level *α* of the considered population. (While it may be computationally prohibitive to compute all solutions, it is often possible to collect a large number of them.) These sets can be combined in at least two obvious ways, namely via their union

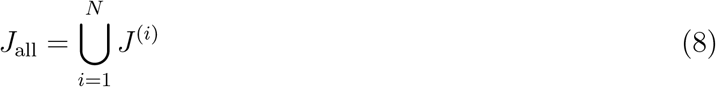

or their intersection

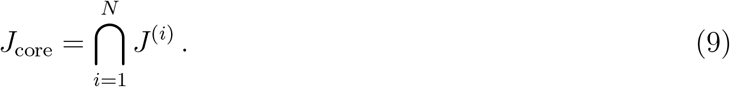

The latter (while not being a covering by itself) we particularly focus on as it captures essential abnormalities observed. Other possibilities include maximizing the sum of probabilities that each element is aberrant.

Once a subset *J* of variables is chosen (a covering, or a core), we obtain a representation of each sample *ω* as a binary vector 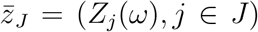, which should retain essential information from the whole *omics* profile associated to the sample. It has, in addition, a mechanistic interpretation, since each variable *Z_j_* is associated to one or a group of STPs (*g* ⇒ *g′*). Because of the relatively small number of variables involved, all these events can be rendered together, using, for example, the visualization provided in Figures 2 or 4.

**Figure 2:**
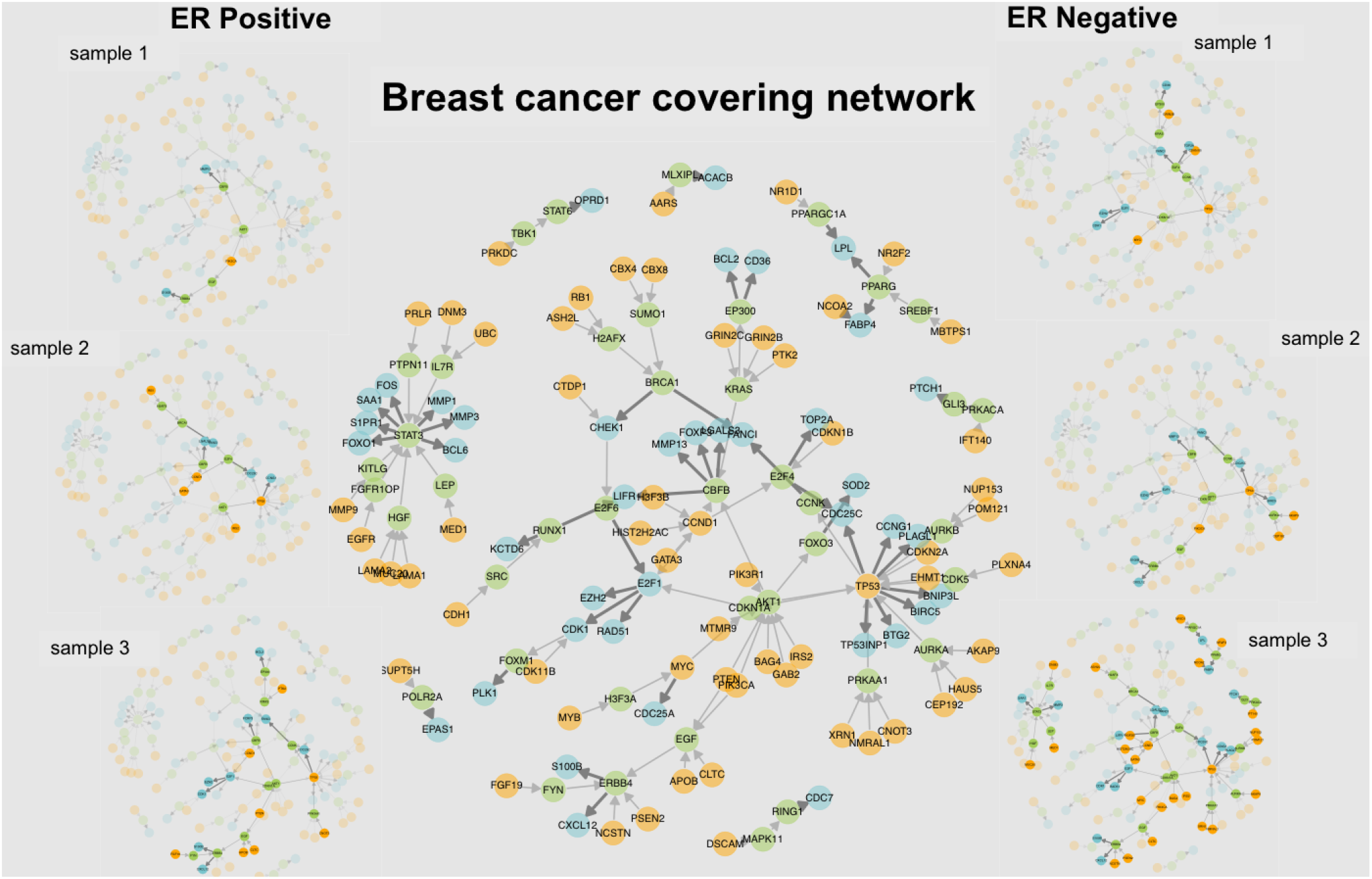
Networks of pair coverings in breast cancer. The network shown in the center depicts one covering of breast cancer samples by STPs, with source genes in orange, target genes in blue, and intermediary link genes in green. The thin and thick edges represent, respectively, the two types of relationships: “controls state change of”and “controls expression of”as designated in Reactome (Jassal et al., 2020). On the left are presented a selection of covering realizations for three ER-positive samples, where aberrant STPs are highlighted, while and on the right, three ER-negative samples samples are shown. The samples have different realizations over the covering network, and are ranked (top to bottom) by the number of events they exhibit. The sample networks demonstrate the inter-sample heterogeneity among the source and target realizations.

### 2.4 Measuring heterogeneity

We want to quantify the heterogeneity of a family of binary random variables 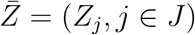, defined on the probability space Ω, where *J* is a subset of 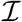 (*e.g.*, a covering). Similarly to the previous section, we assume that only a finite number of observations are available, represented by a finite subset 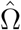 of Ω. A natural measure for heterogeneity is the Shannon entropy (Shannon, 1948; Cover and Thomas, 2012), that we need to estimate based on the finite random sample 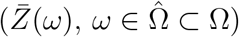. In our results, we focus on the entropy of 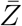 conditional to a specific cancer condition, phenotype or subtype.

Let 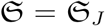 denote the set of all binary configurations 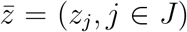, which has 2^|*J*|^ elements. Let 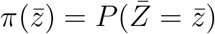, so that the Shannon entropy of 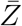 is

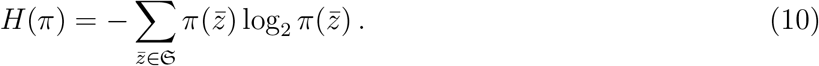

The sample probability mass function of 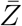 is then given by

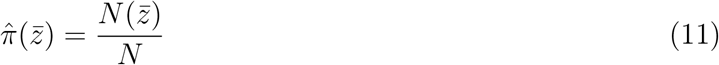

where 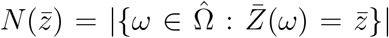 and 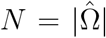. One can plug these relative frequencies in the definition of the entropy to obtain the estimator

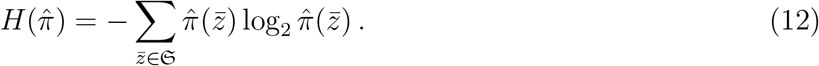

This estimator, however, significantly under-estimates the entropy for small and even moderate sample sizes, and several bias-correction methods have been introduced in the literature (see Schürmann (2004), from which (7) is obtained, and Grassberger (1988, 2003)). This estimator computes the entropy using the expression (in which ***ψ*** denotes the digamma function)

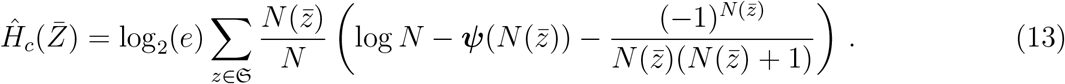

Still, this estimator is only accurate when the number of variables, |*J* |, is small, because the ratio is the first order term in the expansion of the entropy bias (Miller, 1955; Schürmann, 2004) in powers of 1/*N*. In our experiment, we use *L* = 4, that is we estimate the entropy for at most 4 variables together. For a more detailed analysis on the selection of *L* = 4, see Supplementary text. Since sets *J* of interest are typically larger, we estimate an upper-bound to the entropy in the following way.

Given two random variables *X* and *Y*, one always has *H*(*X*, *Y*) ≤ *H*(*X*) + *H*(*Y*). This implies that, if the set *J* is partitioned into subsets *J_1_*, …, *J_l_* (*i.e.*, 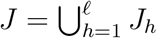 and *J*_*h*_ ∩ *j_h′_* = *∅* if *h* ≠ *h′*), then

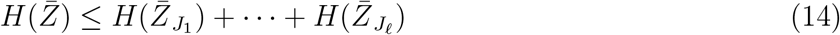

where 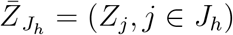, *h* = 1, …, *ℓ*. We use the right-hand side as an upper-bound, determining the partition *J*_1_, …, *J*_*ℓ*_ using the following greedy aggregating procedure:

i. Initialize the partition with singletons, *i.e.*, *J_j_* = {*j*}, *j* ∈ *J*, computing the estimated entropy 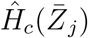 of the binary variable 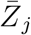. Fix a maximal subset size, *L*.
ii. Given a current decomposition *J*_1_, …, *J_l_*, compute, for all pairs *h*, *h′* such that |*J_h_* ∪ *J_h′_*| ≤ *L*, the difference 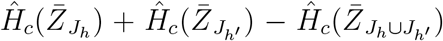 remove the two sets *J_h_, J_h′_* for which this difference is largest and replace then by their union (setting *ℓ* → *ℓ* − 1).
iii. If no pair *h*, *h′* satisfies |*J_h_* ∪ *J_h′_*| ≤ *L* stop the procedure.

The obtained decomposition also provides a statistical model (denoted 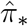) approximating the distribution of 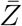 namely the one for which 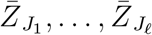 are independent and the distribution of 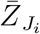 is estimated using relative frequencies. To allow for comparisons between entropies evaluated for different sub-populations, we used this model within a Monte-Carlo simulation to estimate confidence intervals for *H*(*π*). We generated *M* =1,000 new *N*-samples of 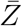 (recall that *N* is the size of the original sample of 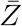 used to estimate the entropy), using the distributions 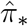 resulting in *M* new empirical distributions 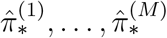 with associated corrected entropies 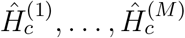. Fixing a probability β > 0, we let 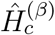 and 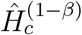 denote the *β* and 1 − *β* quantiles of the sample 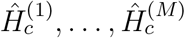 so that 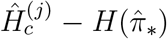 belongs to 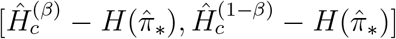 with probability 1 − 2*β*. We use the same interval for the difference 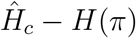, yielding the confidence interval for *H*(*π*):

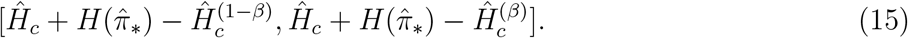

### 2.5 Subtype analysis through partitioning

We assume here again a family of binary random variables 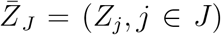, where *J* is a tissue-dependent covering, observed through a finite sample 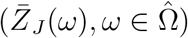 We partition the sample space into disjoint subsets (*S*_1_, …, *S_*ℓ*_*) where each *S_j_* is specified by a small number of events involving conjunctions or disjunctions of aberrations. This partition will be associated with the terminal nodes of a binary “coding tree” of limited depth *d* (*e.g.*, *d* = 5), so that *ℓ* = 2^*d*^. To each node in the tree we associate a subset *S* of 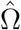, and unless *S* is a terminal node, its children form a partition *S* = *S′ ⋃ S′′* where *ω* ∈ *S′* is based on a certain splitting criterion.

While there are many ways to build such a tree-structured code, we opt for a decomposition for the tissue population that is as balanced as possible and unsupervised to compare the distributions between cancer subtypes (for a given tissue) with respect to a fixed partition. A sample is weighted inversely proportional to the size of its subtype, and at each node the event which balances the weight of the two daughter nodes is selected.

The standard choice for a binary tree are individual binary features, so events of the form {*Z_j_*_(*S*)_ = 1}, for a suitably chosen *j*(*S*) ∈ *J*. One could also use more complex splitting criteria, such as 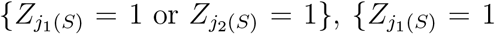 and 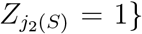 with *j_1_(s)*, *j_2_(s)* ∈ *J*. (We have used both types of events in our experiments: two-gene disjunctions for trees based on source aberrations with targets and two-gene conjunctions for trees based on target aberrations with sources.) The stopping criterion is that either all samples at the node have identical configurations or a maximum depth has been reached.

## 3 Results

As described previously, we have delineated many gene pairs (STP, or “source-target pair”), with a binary random variable corresponding to each pair indicating whether a sample is source and target aberrant.

Given *M* STPs, there are 2^*M*^ possible “states” or “configurations”for each sample. We defined *cross-sample heterogeneity* as the entropy of the probability distribution *P* over configurations. Estimating the entropy of *P* is not feasible for modest sample sizes, since it requires estimating the probabilities of many rare events.

To overcome this computational barrier, the pool of STPs was substantially reduced using the notion of a “minimal set covering” in combinatorial optimization. In our case, the set to be covered is a population of cancer samples for a particular phenotype or subtype, a “covering” is a set of STPs for which, with high probability, cancer samples have at least one aberrant STP from the covering, and “minimal” means the smallest covering. All minimal coverings are necessarily of the same size, on the order of 10 − 100 for each tissue we study (breast, colon, liver, kidney, lung and prostate). In summary, our STPs are derived from Reactome, and then subsets of STPs of interest are identified for each cancer type based on the TCGA omics data.

Minimal set coverings are typically not unique. However, despite the differences between the coverings we can define a “core”, namely, the STPs that appear in *every* (minimal) solution. From a biological perspective, the core is a novel signature of the most salient events associated with tumors of a given type. We apply these concepts (STPs, cores and estimated entropies) to measuring cross-sample heterogeneity in tumor populations for a selection of tissues represented in TGCA data.

### 3.1 Source-target pairs

Based on the genes and interactions found in the Reactome (Jassal et al., 2020), source-target pairs (STPs) (*g_s_* ⇒ *g_t_*) are built, with the only parameter used being the maximum length of the directed chain from the source to the target (*k*; see Supplementary table S1). There are then 272,237 valid STPswith 3,124 distinct source genes and 598 distinct target genes.

Our samples are those in TCGA with available matched mutation, extreme copy number variation (deleting or amplifying both copies), and mRNA expression data; the conversion of expression counts to aberration states was described in Section 2.2.1. Nearly all samples exhibit at least one paired aberration.

### 3.2 Filtering

Given there are too many STPs to meaningfully analyze, we first filter based on rejecting the hypothesis that the existence of source and target aberrations are independent (see Section 2.2.2). The statistics of the STPs remaining after this filtering procedure for different tissue types are shown in Supplementary Table S2. For example, for the 953 TGCA breast cancer samples, there are 17, 261 valid STPs after the test for independence, with 2, 130 source genes and 421 target genes.

Next we omit very rare events. For each tissue, and at each of the three levels (source, target, pair), we require each binary variable to be aberrant in at least 2% of the samples for that tissue. We have a detailed discussion of optimal choice of 2% in Supplement Section. The number of qualifying variables after the 2% filter was applied are given in Supplementary Table S4. For instance, for breast cancer samples, there are 4, 026 STPs, 690 distinct source genes, and 256 distinct target genes after separately applying the 2% filter at each level.

### 3.3 Paired aberrations

Table 1 shows examples of STPs λ = (*g* ⇒ *g*′) and their associated probabilities of aberration in the indicated tissue. For example, in the STPs shown for colon cancer in Table 1, *APC* is the source gene, *AXIN2* is the target gene, and there exists a directed signaling path from *APC* to *AXIN2* of length at most three links (two intermediate genes) in Reactome such that the second-to-last link, namely the direct parent of *AXIN2*, controls the mRNA expression of *AXIN2*. This STP is aberrant in a given sample if *APC* is either mutated, deleted, or amplified *and* the mRNA expression of *AXIN2* is aberrant (with respect to baseline mRNA expression for *AXIN2*). In the case of *APC*, the DNA aberration is nearly always a mutation and *AXIN2* is over-expressed. See supplementary table S3 for more information.

**Table 1:** Examples of STPs. For each of the six tissues, one example of a common STP *λ* = (*g* ⇒ *g*′) is shown. *P* (DNA&RNA) is our sample-based estimate of the probability that *λ* is an aberrant pair, namely, the fraction of samples of the indicated tissue for which the source gene *g* is DNA-aberrant and the target gene *g′* is RNA-aberrant. Similarly, *P* (DNA) (respectively, *P* (RNA)) is the fraction of samples for which *g* is DNA-aberrant (resp., *g′* is RNA-aberrant), and *P* (RNA|DNA) is the (estimated) conditional probability that *g′* is RNA-aberrant given *g* is DNA-aberrant.

| Tissue | Pair | $P(\text{DNA\&RNA})$ | $P(\text{DNA})$ | $P(\text{RNA})$ | $P(\text{RNA}\vert \text{DNA})$ | $P(\text{DNA}\vert \text{RNA})$ |
| --- | --- | --- | --- | --- | --- | --- |
| Breast | $PIK3CA \Rightarrow S100B$ | 0.316 | 0.356 | 0.838 | 0.888 | 0.377 |
| Colon | $APC \Rightarrow AXIN2$ | 0.585 | 0.739 | 0.676 | 0.791 | 0.864 |
| Kidney | $VHL \Rightarrow CA9$ | 0.482 | 0.485 | 0.967 | 0.994 | 0.498 |
| Liver | $TP53 \Rightarrow MYBL2$ | 0.308 | 0.319 | 0.814 | 0.965 | 0.379 |
| Lung | $TP53 \Rightarrow TOP2A$ | 0.529 | 0.535 | 0.923 | 0.988 | 0.573 |
| Prostate | $PTEN \Rightarrow TWIST1$ | 0.161 | 0.216 | 0.654 | 0.745 | 0.246 |

The probability *P* (DNA&RNA) is the sample estimate, namely the fraction of colon samples for which *APC* is mutated *and AXIN2* is RNA-aberrant. Similarly, *P* (DNA) and *P* (RNA) stand for the marginal probabilities that the source is DNA-aberrant and the target RNA-aberrant, respectively. The conditional probabilities are then self-explanatory. For example, *APC* is mutated in 73.9% of our samples and in 79.1% of those samples *AXIN2* is RNA-aberrant. Multiplying these two probabilities gives the frequency of the joint occurrence (58.5%). Other STPs commonly found in colon samples include the four core STPs described in Table 3.

The probabilities for *APC* ⇒ *AXIN2* are atypically large. In particular, most pair probabilities are smaller than.575, generally of order 0.01–0.10 with a few above 0.3, usually involving main tumor drivers such as *PIK3CA* in breast cancer, and *TP53* and *KRAS* in lung cancer. Moreover, DNA aberrations tend to be considerably rarer than RNA aberrations, *i.e.*, the marginal source probabilities are generally far smaller than the marginal target probabilities. It is noteworthy that the conditional probability of a particular RNA aberration given a particular DNA aberration (as those in Table 1) is usually in the range 0.5–1, whereas the reverse is not the case: given a target gene is RNA-aberrant the probability of any particular gene serving as a source rarely exceeds 0.2 (see Supplementary Tables S5–S9).

We have also defined separate source-level and target-level events in the sense of partially aberrant STPs; see Section 2.2.2 of Methods. Recall that 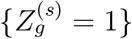 represents the event that a given source gene *g* is DNA-aberrant *and that there exists some* target of *g* which is RNA-aberrant, denoted as “aberration with target”for “source aberration with downstream target aberration”. The probability of this event is denoted by *P* (DNA & *downstreamream*RNA); see Table 4 for some examples in Colon. Similarly, for the other direction, 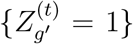 is the event that *some* source gene renders *g ⇒ g′* an aberrant STP. Table 4 and Table 5 provide the probabilities and conditional probabilities for selected core genes at the source and target levels in colon; many other examples appear in Supplementary Tables S5 to S19.

Given a source gene *g* is aberrant, typically there is a strong likelihood that at least one of its targets *g′* is RNA-aberrant. These targets represent plausible downstream consequences of *g* having a DNA-aberration. The converse, however, is not valid; in particular, there are many targets *g′* for which there is no upstream DNA-aberrant source linked to *g′*. This makes sense since a gene can be RNA-aberrant for many reasons other than an upstream genetic aberration. In particular, the event driving the aberration of *g′* might be some perturbation not considered here, for example be epigenetic or fusion-related or as yet unrecognized.

### 3.4 Coverings

Recall that indexing a covering by source genes refers to leaving the particular aberrant target gene unspecified (indexing by targets is the opposite). The corresponding events were denoted in Methods by 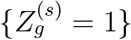 for a source gene and 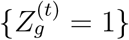 for a target gene.

As described in Section 2.3, minimal coverings composed of pairs, sources, or targets are all found with the same optimization program. For the pair and source levels, we calculate the optimal covering with the smallest possible *α* ≥ 0 and *r* = 1. (Here, the smallest *α* is such 1 − *α* is the fraction of samples that have at least *r* aberrant STPs.) At the target level, however, we select but *r* = 3, still using the smallest possible *α*; that is, we attempt to cover tumor samples with at least three target aberrations (with source). This choice is justified by the higher frequency of RNA-aberrations in tumor samples.

Table 2 shows the optimal covering statistics at all levels for the six tissues of origin. For example, the minimum number of STPs (resp., sources, targets) necessary to cover the 953 breast cancer samples is 67 (resp., 60, 53), with the realized rate being 95% (resp., 96%, 96%) with *α* = 0.05 (resp., 0.04, 0.04). In contrast, all colon samples can be covered with many fewer STPs, namely 11. In addition, the minimal covering size (size of solution) is usually largely determined by the incidence of aberrations in any given population, e.g., mutation rates. In particular, given two phenotypes A and B, if the samples of B are consistently more aberrant than those of A, then the minimal B covering will be smaller. More comprehensive statistics for all tissues are given in Supplementary Table S4; as seen in the column labeled “No. of solutions”, there are in general a great many instances of minimal coverings.

**Table 2:** Statistics of optimal coverings. For each of the six tissues, this table provides basic information about the optimal coverings at all levels: STP, source with target, target with source. For instance, for breast cancer, there are 4,026 candidate STPs after both filters (rejecting source-target independence and 2% tissue sample frequency); the minimal covering size is 67 STPs; at least one of these 67 STPs is aberrant in 95.4% of the breast cancer samples; and there are 21 STPs which appear in *every* minimal covering.

| Tissue | Samples | Covering Type | Quantity | Size of solution | Fraction of samples covered | Size of core set |
| --- | --- | --- | --- | --- | --- | --- |
| Breast | 953 | STP | 4,026 | 67 | 0.954 | 21 |
|  |  | Source | 690 | 60 | 0.964 | 34 |
|  |  | Target | 256 | 53 | 0.955 | 35 |
| Colon | 207 | STP | 1,195 | 11 | 1.000 | 4 |
|  |  | Source | 525 | 10 | 1.000 | 5 |
|  |  | Target | 226 | 15 | 0.995 | 6 |
| Kidney | 336 | STP | 347 | 26 | 0.827 | 12 |
|  |  | Source | 133 | 28 | 0.854 | 21 |
|  |  | Target | 176 | 60 | 0.890 | 45 |
| Liver | 360 | STP | 1,198 | 32 | 0.931 | 11 |
|  |  | Source | 460 | 34 | 0.958 | 20 |
|  |  | Target | 287 | 41 | 0.942 | 26 |
| Lung | 465 | STP | 3,154 | 27 | 0.985 | 10 |
|  |  | Source | 908 | 25 | 0.989 | 19 |
|  |  | Target | 350 | 29 | 0.985 | 26 |
| Prostate | 491 | STP | 430 | 53 | 0.686 | 32 |
|  |  | Source | 211 | 53 | 0.743 | 42 |
|  |  | Target | 160 | 72 | 0.699 | 66 |

**Table 3:** Colon core STPs. There are four “core” STPs which appear in every minimal covering of
the colon samples. P(DNA&RNA) is the fraction of samples for which the source gene *g* is DNA-aberrant
and target gene *g′* is RNA-aberrant; P(DNA) is the fraction of samples satisfying the source gene *g* is
DNA-aberrant; P(RNA) is the fraction of samples with *g′* RNA-aberrant; P(RNAjDNA) is the fraction of
DNA-aberrant samples for which *g′* is RNA-aberrant.

| | $P(\text{DNA \& RNA})$ | $P(\text{DNA})$ | $P(\text{RNA})$ | $P(\text{RNA}\vert \text{DNA})$ | $P(\text{DNA}\vert \text{RNA})$ |
| --- | --- | --- | --- | --- | --- |
| $APC \Rightarrow AXIN2$ | 0.585 | 0.739 | 0.676 | 0.791 | 0.864 |
| $TP53 \Rightarrow PTPN12$ | 0.396 | 0.560 | 0.604 | 0.707 | 0.656 |
| $PIK3CA \Rightarrow TNFRSF10B$ | 0.198 | 0.271 | 0.589 | 0.732 | 0.336 |
| $MAML1 \Rightarrow PBX1$ | 0.034 | 0.039 | 0.401 | 0.875 | 0.084 |

**Table 4:** Colon core source genes. There are five “core”source genes which appear in every minimal source covering of the colon samples. *P* (DNA) is the fraction of samples for which the indicated source gene is DNA-aberrant; *P* (DNA & downstream RNA) is the fraction of samples for which the indicated source gene is DNA-aberrant and there exists an RNA-aberrant gene among its targets. *P* (downstream RNA|DNA) is the fraction of the samples with the indicated source gene DNA-aberrant for which there exists some RNA-aberrant gene among its targets.

| | $P(\text{DNA \& downstream RNA})$ | $P(\text{DNA})$ | $P(\text{downstream RNA}\vert \text{DNA})$ |
| --- | --- | --- | --- |
| *APC* | 0.585 | 0.739 | 0.791 |
| *TP53* | 0.560 | 0.560 | 1.000 |
| *KRAS* | 0.425 | 0.425 | 1.000 |
| *LAMA5* | 0.217 | 0.217 | 1.000 |
| *MAML1* | 0.034 | 0.039 | 0.875 |

**Table 5:** Colon core target genes. There are six “core”target genes which appear in every minimal target covering of the colon samples. *P* (RNA) is the fraction of samples for which the indicated target gene is RNA-aberrant; *P* (RNA & upstream DNA) is the fraction of samples for which the indicated target gene is RNA-aberrant and there exists an DNA-aberrant gene among its sources. *P* (upstream DNA|RNA) is the fraction of the samples with the indicated gene RNA-aberrant for which at least one of its sources is DNA-aberrant.

| Target | $P(\text{RNA \& upstream DNA})$ | $P(\text{RNA})$ | $P(\text{upstream DNA}\vert \text{RNA})$ |
| --- | --- | --- | --- |
| *PERP* | 0.710 | 0.807 | 0.880 |
| *PDX1* | 0.671 | 0.957 | 0.702 |
| *AXIN2* | 0.662 | 0.676 | 0.979 |
| *SALL4* | 0.638 | 0.918 | 0.695 |
| *TNFRSF10B* | 0.565 | 0.589 | 0.959 |
| *MYBL2* | 0.261 | 0.300 | 0.871 |

Figure 2 shows one such tissue-level covering obtained for breast cancer as a graphical network with nodes representing genes forming the STPs. The source and target genes are shown in orange and blue respectively, while genes representing the intermediary links are shown in green. Note that while the union of the coverings may also be visualized in a similar manner, it contains many more STPs that make readability of the resulting graph difficult; therefore we have opted to show only individual coverings here. Supplementary Figures S1 – S5 depict the networks associated with the coverings obtained for the other types of cancer.

These visual representations allow us to go beyond lists of names and numbers and begin to interpret coverings in biological terms and incorporate mechanism (see Discussion). For instance, in the breast network shown in Figure 2, several important breast cancer genes (e.g., *STAT3* (Huynh et al., 2019), *TP53* (Olivier et al., 2010), *BRCA1* (Semmler et al., 2019), and *ERBB4*(Segers et al., 2020)) all form important hubs through which multiple sources and targets in the covering link according to Reactome. Similarly, the network figures for the remaining networks show similar positioning for many important cancer genes: *NOTCH1* (Viatour et al., 2011) in liver, *NOTCH3* (Aster et al., 2017; Nowell and Radtke, 2017) and *EGFR* (Sigismund et al., 2018) in lung are some other examples. Finally, *KRAS*, *TP53* (Olivier et al., 2010) and *STAT3* (Huynh et al., 2019) make an appearance in multiple cancers. See Supplementary Figures S1–S5.

For a given tissue and fixed covering level, about 30%–60% of the genes appearing in any covering in fact appear in all coverings, referred to as the *core set* (see Tables 3, 4 and 5 for colon, and in Supplementary Tables S5–S19 for the other tissues). The STP *TP53* ⇒ *PTPN12* is aberrant in 39.6% of the colon samples (see Table 3), the source gene *KRAS* in 42.5% of samples (see Table 4), and the target gene *PERP* in 80.7% of samples (see Table 5). From Table 4 we see that there is some aberrant target for *every sample* for which *KRAS* is DNA-aberrant in colon; hence the probability that *KRAS* is DNA-aberrant *and* there is a matching target gene is again 42.5%. Finally, from the target covering we see that targets gene *PDX1* is RNA-aberrant in 95.7% of colon samples (see Table 5) but only 70.2% of samples for which *PDX1* is RNA-aberrant have some corresponding corresponding upstream DNA-aberrant source gene.

Figure 3 shows 18 core source genes across multiple tissues. *TP53* is a core source gene shared by all 6 tissues, and is DNA-aberrant in more than 60% of colon cancers, and also a large percentage in other cancers. All core source genes across multiple tissues are shown in Figure S10. In Supplementary Figure S11, we show all core target genes across multiple tissues. For instance, *CDC25C* appears in core set of breast, lung and prostate cancer, and the probability that *CDC25* is RNA aberrant and there exist an upstream aberrant source is nearly 0.8.

**Figure 3:**
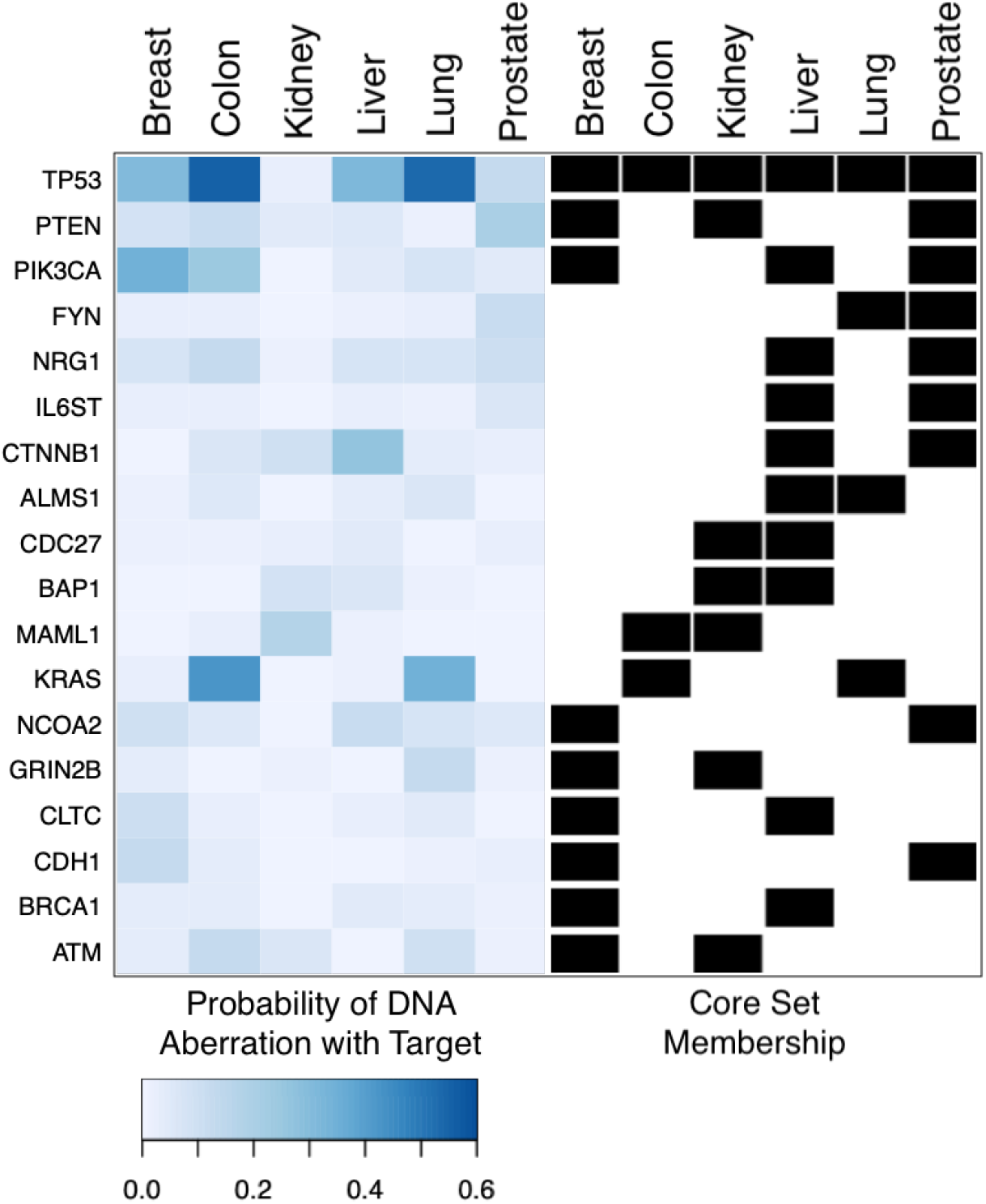
Core set across tissues at source level. There are 18 source genes which appear in the core set of at least two tissues. For instance, gene *TP53* is a core gene for all six tissues, and genes *PTEN* and *PIK3CA* are core genes for three tissues. The color in the heatmap on the left represents the probability that the corresponding source gene is DNA-aberrant and there exists an RNA-aberrant target gene (thereby forming an aberrant *source-target pair*). On the right, black marks indicate the membership of each gene to the corresponding core set for each tumor type.

**Figure 4:**
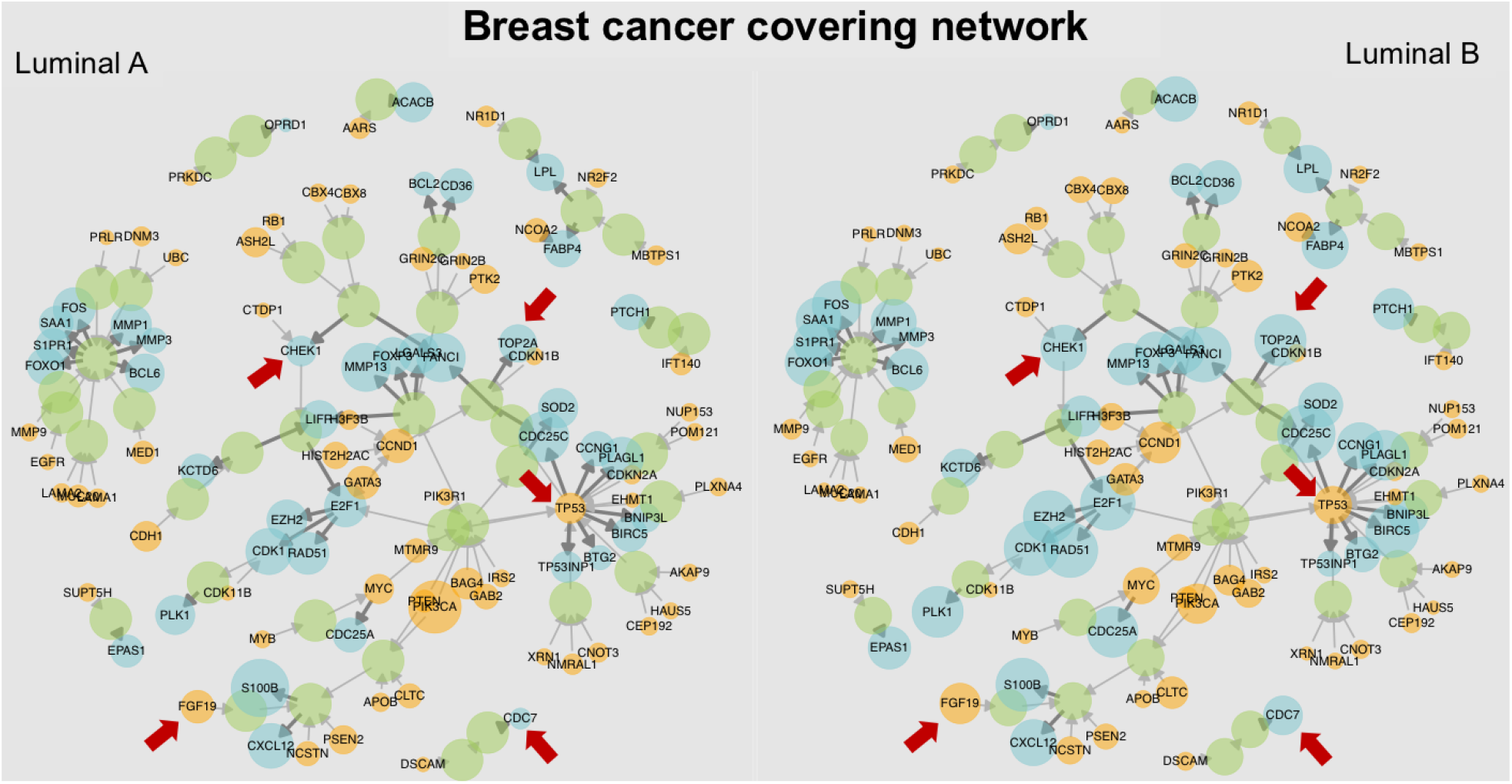
Comparison of one covering network for luminal breast cancer subtypes. The probabilities of DNA aberration (with targets) and RNA aberration (with sources) over the Luminal A and Luminal B populations of breast cancer are depicted by the size of each node in the network, which corresponds to one possible covering. The red arrows indicate some sources and target genes that have noticeable differences in the respective probabilities between the two luminal subtypes (*e.g.*, *TP53*, *CHEK1*, *PIK3CA*, and *TOP2A*, also see Table 6).

### 3.5 Subtype coverings

Having computed the tissue-level coverings, we examine them with respect to certain phenotypes of interest, including the PAM50 subtypes in breast cancer (Parker et al., 2009), smoking history in lung cancer (Pfeifer et al., 2002), Gleason grade in prostate cancer (Humphrey, 2004), and and the CRIS-classes in colon cancer (Isella et al., 2017). We observe a large range of aberration frequencies among subtypes. Table 6A shows the probabilities of DNA-aberration (with targets) for PAM50 subtypes, with genes selected from the core set of source breast cancer coverings; other panels show similar selections of sources.

**Table 6:**
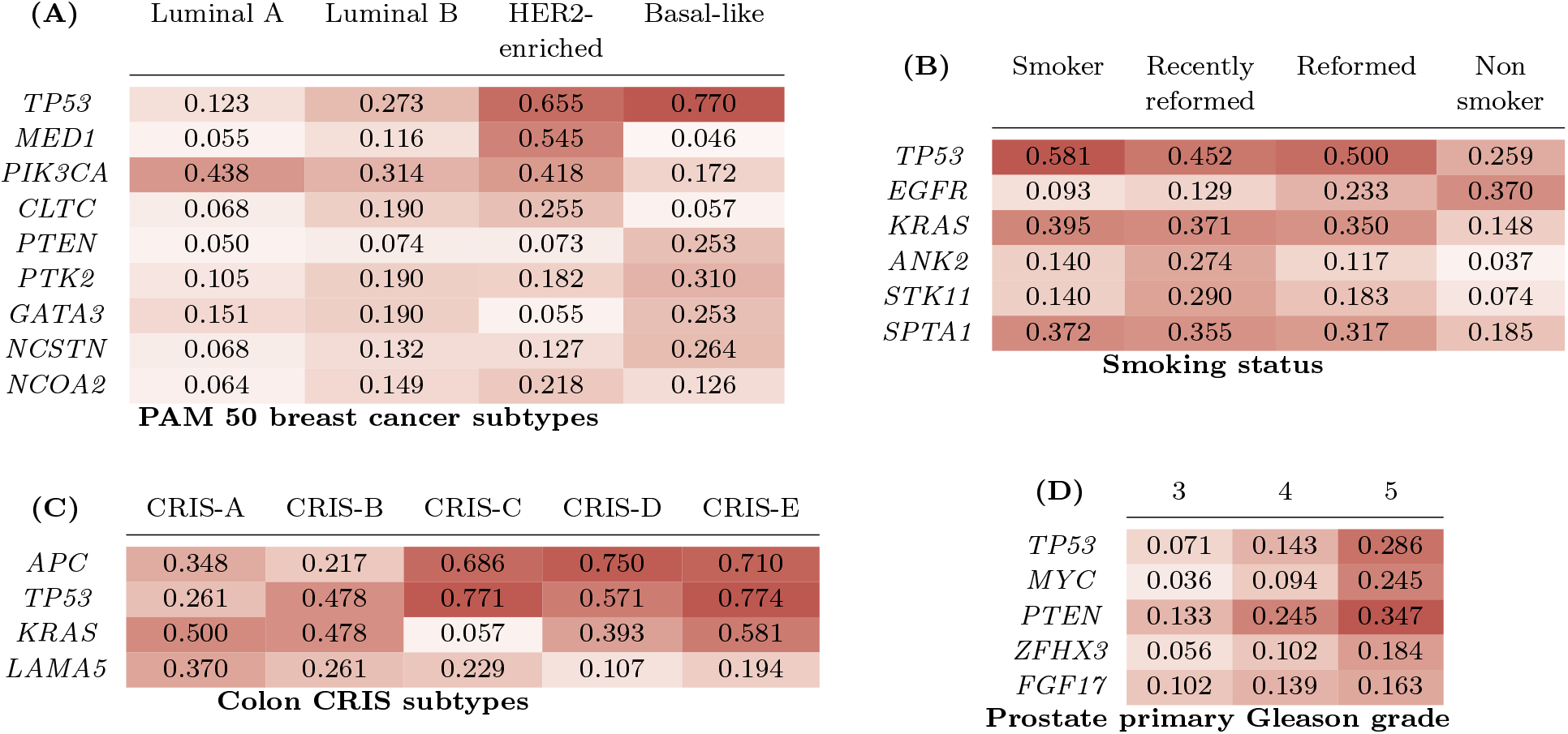
Probabilities of source aberration with downstream target. For various subtypes of breast, lung, colon and prostate cancer, the heatmaps represent the probabilities that the indicated gene is a DNA- aberrant source gene with some downstream RNA-aberrant target. The sources are selected from the set of core genes for coverings of the given tissue; the selection criterion is that the probability of a DNA-aberration is high for at least one of the subtypes for that tissue. Core sources with varying probabilities present interesting candidates for discrimination between subtypes. For example, among the PAM50 subtypes, the DNA-aberration frequency of *TP53* is much higher in the HER2-enriched and Basal-like subtypes than in Luminal A and Luminal B, whereas an aberration in *PIK3CA* is less frequent among basal-like samples than among the other subtypes. In the case of smoking history in lung cancer samples, *TP53* and *KRAS* are both more frequently DNA-aberrant (with some downstream RNA-aberrant target) among smokers than non-smokers whereas *EGFR* is a more aberrant source among non-smokers.

We observe potential discriminating *sources* between subtypes. For example, *TP53* has a much lower likelihood of aberration for the luminal subtypes in comparison to the basal-like and HER2- enriched subtypes. Similar observations can be made among the subtypes of other cancers (see Table 6). Finally, such patterns persist for target-level analyses and are presented in Supplementary Tables S20 to S24.

A comparison between subtypes can also be captured as a graphical network, as shown in Figure 4. Supplementary Figure S6 presents the breast covering with the size of the nodes representing the source (with target) and target (with source) aberration probabilities for the subtypes considered. Similar networks for lung cancer with respect to smoking history are presented in Supplementary Figure S7 and Supplementary Figure S8 presents primary Gleason grade for prostate cancer.

We also compared coverings of subtypes controlling for population sizes. For each of the phenotypes under a given sub-typing, an equal number of samples were selected and coverings for all these samples simultaneously were obtained. Then we examined the proportion from each subtype that was covered, repeating over multiple sampling iterations (Figure 5). A general pattern of more pathological phenotypes having higher coverage proportions can be observed throughout these results (see Supplementary Figure S9 for further results). The more malignant phenotypes tend to have larger aberration probabilities. This corresponds with the observation that the size of the covering obtained for a subtype while sampling equal numbers from each group indicated larger covering solutions obtained for more benign subtypes in comparison to more malignant ones.

**Figure 5:**
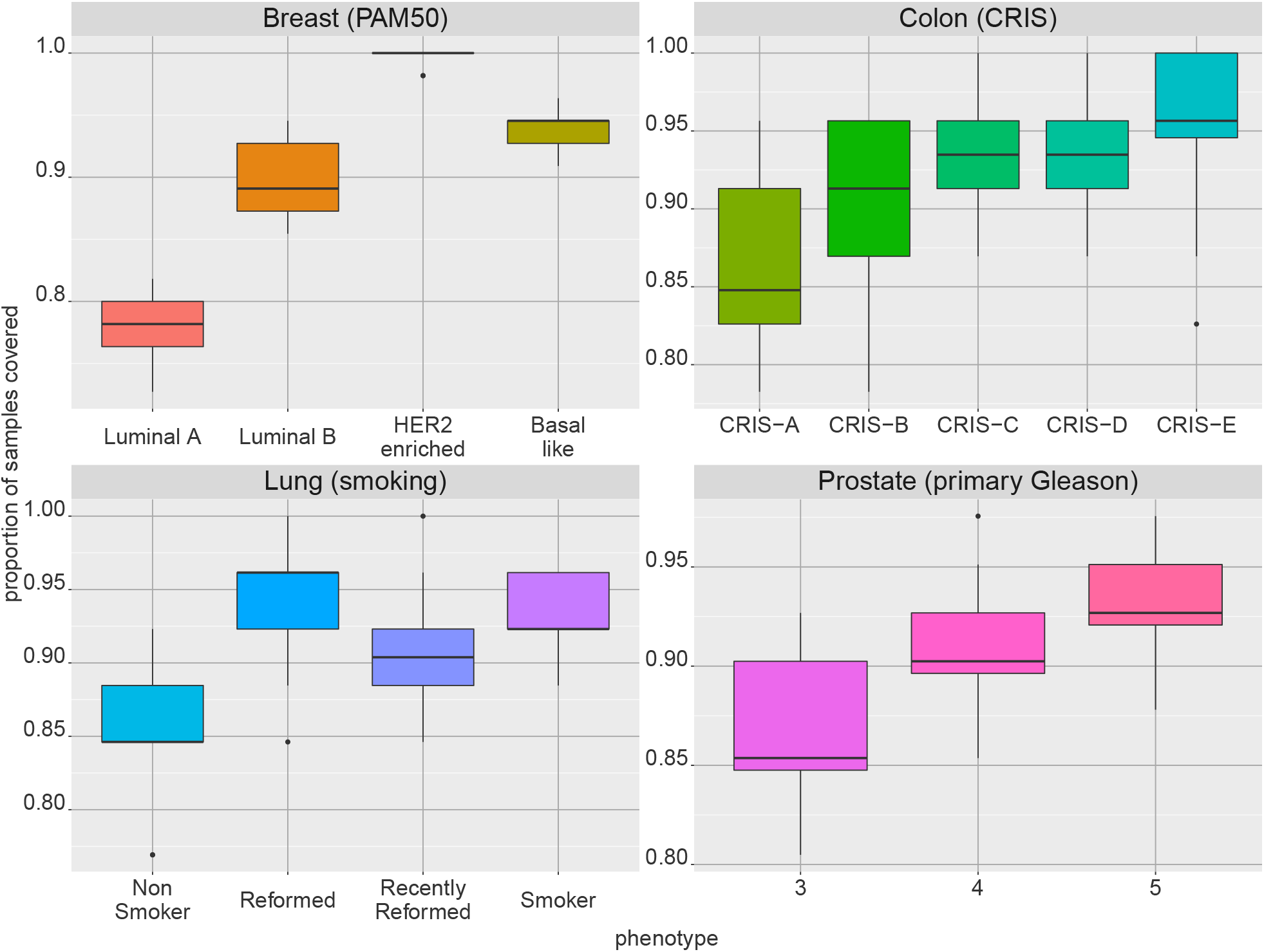
Rates of covering assembly. For each of four tissues (breast, colon, lung and prostate), several phenotypes are compared based on the proportion of samples actually covered when requesting 90% coverage or more for the given tissue by the optimization procedure. The boxplots represent the results of 20 iterations of normalizing for sample size among the phenotypes by random sampling. In general, coverings for more aggressive phenotypes assemble faster.

### 3.6 Measuring heterogeneity

We applied the approach described in Section 2.4 to assess the relative heterogeneity of different cancer phenotypes and subtypes in the analyzed tissues. Note that we are analyzing heterogeneity within phenotypes at a population level, so that our measurements are primarily about the variability across tumors in this population. Without single-cell data, one cannot evaluate variability within a tumor, although it is likely that a higher variability at this level should also trigger larger heterogeneity at the population level. It is worth noting, however, that the analytical framework we propose here can be easily extended to single cell data once paired molecular measurements will become available in the future.

We base our analysis on coverings estimated on source aberration with targets and on target aberration with sources (see Supplemental Tables S26 and S27). In all cases, coverings are obtained for each tissue of origin separately, and entropy estimates are computed after restricting the data to samples exhibiting each considered cancer phenotype (*e.g.*, the breast cancer molecular subtypes, smoking status in lung cancer, and so on…).

At the source level, the general trend is that heterogeneity estimates increase with increasing disease severity. In prostate cancer, for instance, entropy grows with Gleason sum, primary Gleason grade, and with tumor status, while no clear ordering is observable for lymph node status (see Table 7). Similar observations can be also made for tumors originating in other tissues (see Supplementary Table S26). In breast cancer, the entropy for ER positive tumors is less than that for ER negative ones, and it also increases with tumor size, and with more aggressive molecular subtypes (*i.e.*, Luminal A *<* Luminal B *<* HER 2 *<* Basal, with a small overlap between confidence intervals for Luminal B and HER 2). For lung, samples from patients with recent smoking history (reformed for less than 15 years or current smokers) have a higher entropy than those with either ancient or no history.

**Table 7:** Entropy estimation at source level. Entropy estimates for source aberrations with target for prostate Gleason sum, primary Gleason grade, tumor status, and lymph-node status. *N* is the total number of samples available in the given subtype.

| Subtype | Value | N | Entropy | Conf. Interval |
| --- | --- | --- | --- | --- |
| All |  | 491 | 12.16 | [11.67, 12.68] |
| Gleason sum | 6 | 45 | 8.05 | [6.82, 8.98] |
| Gleason sum | 7 | 244 | 10.80 | [10.10, 11.45] |
| Gleason sum | 8 | 63 | 11.33 | [10.07, 12.41] |
| Gleason sum | 9 | 135 | 14.21 | [13.24, 15.06] |
| Primary Gleason grade | 3 | 196 | 9.45 | [8.68, 10.12] |
| Primary Gleason grade | 4 | 245 | 12.33 | [11.65, 13.01] |
| Primary Gleason grade | 5 | 49 | 16.60 | [15.07, 17.88] |
| Tumor Status | T2 | 186 | 10.73 | [9.92, 11.45] |
| Tumor Status | T3-T4 | 298 | 12.88 | [12.26, 13.52] |
| Lymph Node Status | Negative | 342 | 12.14 | [11.55, 12.71] |
| Lymph Node Status | Positive | 77 | 12.82 | [11.66, 13.76] |

At the target level, a similar trend of increasing heterogeneity with increasing disease severity is observed: in prostate cancer for all variables considered (Gleason sum, primary Gleason grade, tumor stage, size, and lymph node status), in kidney cancer for tumor stage and tumor size, and in breast cancer for the molecular subtypes (with Luminal A samples exhibiting the lowest heterogeneity while Luminal B the highest). Finally, in the tumor types originating in the other tissues, we observe large overlaps between confidence intervals, and no obvious and clear trends emerged across cancer subtypes. Complete summaries for this analysis can be found in Supplementary Table 7.

### 3.7 Partitioning

We applied the approach described in Section 2.5 to all six cancer types. Let *T* denote a coding tree with terminal nodes {*t*_1_*, t*_2_*, …, t_l_*}. Recall that for each subtype, the resulting histogram (number of samples per bin) is, by construction, as balanced as possible for the whole population. It is then easy to visualize the histograms conditional on tumor sub-populations (samples for a given subtype), and assess differences across subtypes or phenotypes in this representation (See Figure 6, for ER status in breast cancer based on target aberration with sources).

**Figure 6:**
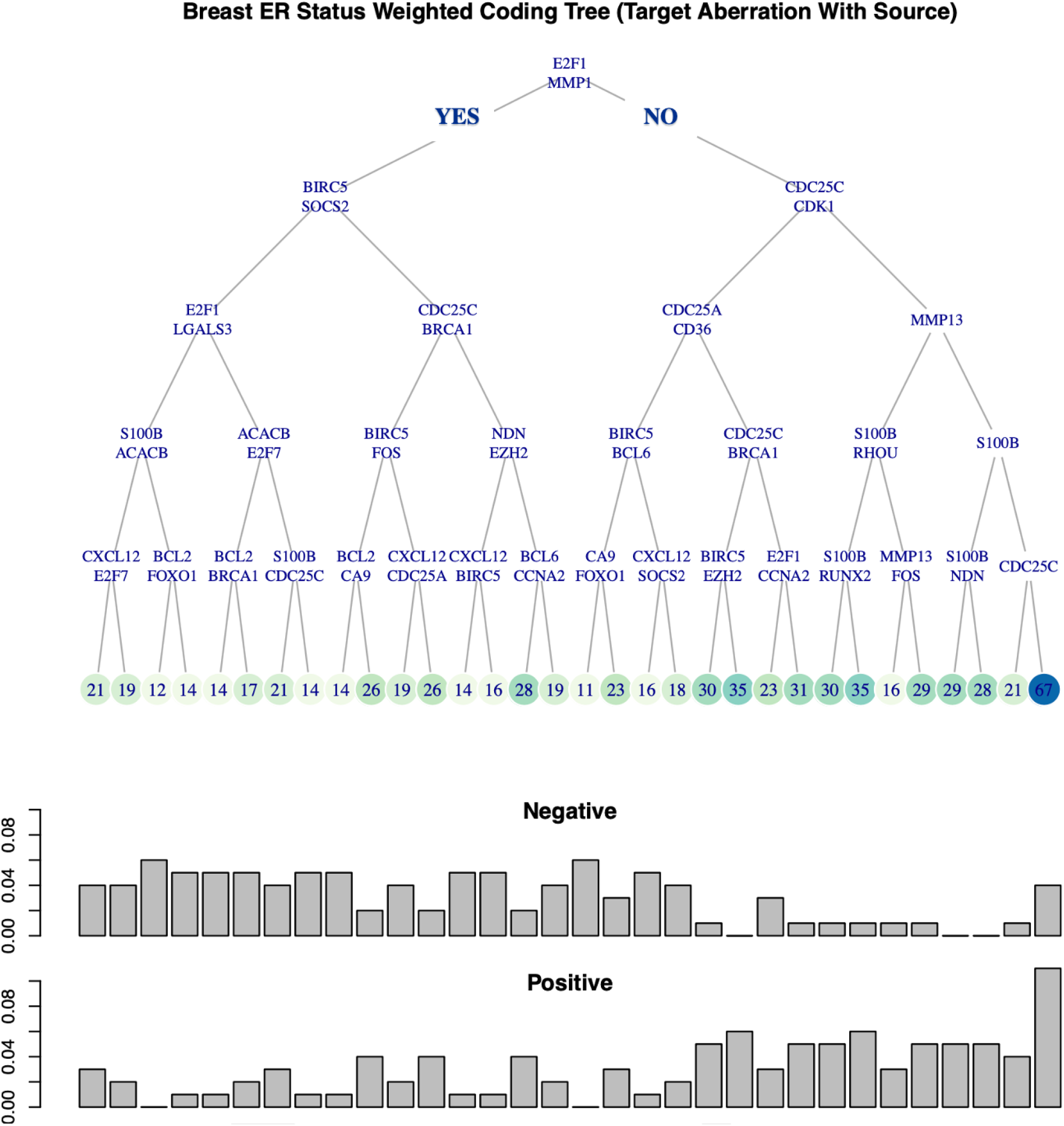
Coding tree for breast ER status at target level. For Breast tumor samples, a weighted coding tree *T* with depth *d* = 5 is constructed using one covering at target level. At each internal node, a sample is sent to the left if the indicated two genes at the node are both RNA aberrant with some aberrant source, whereas it is sent to the right otherwise. Sample counts for each terminal nodes are indicated and highlighted using a green palette. The two histograms at the bottom show the sample distribution at the 32 terminal nodes for the ER negative and ER positive sub-populations.

For instance, the event chosen at the root node is whether both *E2F1* and *MMP1* are RNA aberrant, each with some upstream DNA aberrant source. Such samples take the left branch whereas all others go to the right. The number in the terminal node *t* is the overall number of tumor samples reaching *t* (regardless of subtypes). The two histograms show the numbers of samples collected at terminal nodes for the ER positive and the ER negative sub-populations. In particular, the two distributions are significantly different (permutation test *p* ≤ 0.0001). Other coding trees for other cancer subtypes are showed in Figures S12–S14.

## 4 Discussion

Our approach in introducing this framework is rooted in the biological tenant that cancer is driven by genetic and genomic alterations that alter normal cell behavior through the modification of down-stream gene expression programs governing cell proliferation, cell identity, and cell differentiation. The representation of a cancer profile is binary and integrative, centered on STPs which encode paired DNA and RNA aberrations. The set of possible pairings is fixed, based on signaling pathways and regulatory network topology; the STP is aberrant in an individual profile if the first, or “source”, gene carries the specified DNA aberration and the gene expression of the second, or “target”, gene diverges from the normal baseline. Note that we are not assuming that the DNA alterations are decidedly always “drivers”of cancer or that the source-target links are necessarily causative; rather, given an aberrant STP, we regard the aberrant target as a putative effect of the upstream source aberration, which itself provides a putative explanation for the aberration of the target.

Whereas we do not deal directly with the driver-passenger distinction, *i.e.*, we treat all mutations in the same way, most of the genes that emerge from our analysis are known drivers, particularly the core covering. This is further expected given our requirement that candidate DNA aberrations have at least 2% incidence in the cancer population. That said, given the heterogeneous nature of cancer as a disease, it is likely that the list of currently accepted, known cancer drivers is not exhaustive. In this perspective, our approach—requiring additional constraints for defining aberrant STPs—might also be helpful in prioritizing previously unknown driver events from the large set of potential candidates.

We apply STPs, together with integer programming, to extract parsimonious sets of important aberrations in the tumor populations. In addition to minimal coverings with paired aberrations, the same algorithm can be applied to find the minimal coverings by DNA aberrations alone for which there is some downstream effect (the source gene appears in some aberrant STPs), and vice-versa for coverings by RNA-aberrant target genes (*i.e.*, gene expression alterations plausibly associated with some upstream DNA aberrations).

UNCOVER by (Basso et al., 2019) utilizes a similar computational method, attempting to distinguish driver and passenger somatic event sets with a high degree of mutual exclusivity between alterations. As posed in (Basso et al., 2019), the problem is NP-hard (Aho and Hopcroft, 1974) and only small sets of genes, usually of size three or four, can be found. In contrast, we seek to cover a tumor population with sets of paired aberrations, of size usually ranging from 10-100. Our use of an apriori network of interactions and binarization of aberrations allow this to computationally feasible.

The search for minimal coverings somewhat resembles the body of work on maximal frequent sets (Pasquier et al., 1999; Alves et al., 2010) whose goal is to list *all* “closed”sets of binary variables that are aberrant in a large proportion of the samples, where closedness refers to some notion of maximality among such frequent sets. In contrast, with minimal coverings the sets *intersect* the aberrant set of a large proportion of the samples (the frequent set problem is of polynomial complexity (Uno et al., 2005) while the minimal covering problem is NP hard).

In (Geistlinger et al., 2011), the authors presents Gene Graph Enrichment Analysis (GGEA) to detect consistently and coherently enriched gene sets based on prior knowledge driven from a directed gene regulatory network. And in (Vaske et al., 2010), the authors proposes a probabilistic graphic models (PARADIGM) based on factor graphs to infer patient specific genetic activities incorporating curated pathway interactions among genes. In contrast we do not score gene interactions, instead we use the core set to quantify tumor heterogeneity.

### 4.1 Cores

In general the minimal coverings are not unique; of particular interest are the core STPs which are those belonging to *every* minimal solution. The restriction to coverings, especially to cores, massively reduces the number of considered aberrations in a given tumor population, making it mathematically feasible to quantify and measure tumor heterogeneity at the population level in a natural, information-theoretic way.

Overall, we were able to identify well-known cancer aberrations, as well as to uncover novel potential molecular circuits involved in this disease. The presence of tumor suppressor *TP53* in all cores confirm the well-known notion that this DNA alteration is the most frequent and (possibly) important aberration across multiple tumor types of different lineage (Levine and Oren, 2009). Other aberrations involving key cancer drivers, are discussed in our supplementary results section.

No single gene appears in the core target set of all tissue types. In fact, only one gene, *FABP4*, lies in at least four out of six tissue-specific core signatures. Irrespective of source, target, or pair level, all core genes belong to signaling pathways commonly disrupted in the analyzed cancer types. For instance, paradigmatic examples of affected cancer pathways emerging from our analyses are *Ras* and *Wnt* in colon cancer and the *PI3K* and *mTor* pathways in breast and prostate cancers.

More unexpected and presumably novel genes of interest include for instance, *GRIN2B*—a gene encoding for a subunit of a N-methyl-D-aspartate (NMDA) receptor family member—is a source-level core gene for both breast and kidney cancer. Despite a relatively low incidence of DNA aberration (2.1% and 3.6% respectively in kidney and breast cancer), this gene was always associated with the divergent expression of a downstream RNA in both tumor types. Our findings, along with the previously reported promoter hyper-methylation observed in gastric (Liu et al., 2007), esophageal (Kim et al., 2006), and lung cancer (Tamura et al., 2011), collectively suggest this gene might play a role as tumor suppressor.

Another interesting example is *PTK6* which encodes a cytoplasmic protein kinase also known as breast cancer kinase. This is a core target gene in breast, lung, and prostate cancer, with high probabilities of RNA aberration and upstream DNA aberration in breast and lung (17.2% and 50.1% respectively). Possibly, effects of different upstream DNA alterations can propagate and converge on downstream targets to explain their aberrant expression. Thus *PTK6* could represent a suitably “unifying”target for treatment, despite the heterogeneous set of mutations observed in the patient population. Interestingly, inhibition of *PTK6* has been proposed for treatment in triple negative breast cancer (Ito et al., 2016) and *PTEN*-null prostate cancer (Wozniak et al., 2017).

In addition, our analyses also point to specific interaction pairs, further underscoring the importance of adopting a network view that goes beyond “hubs”, individual genes, and known cancer driver, when interpreting the core sets. Table 3 include a number of pairs for colon cancer that can be directly mapped to specific signaling pathways. The *APC* ⇒ *AXIN* 2 pair participates into Wnt signaling, while the *MAML*1 ⇒ *PBX*1 pair is part of the Notch3 signaling network. Both these pathways are known to regulate the homeostasis of the colonic epithelium, and their alterations are well documented in colon cancer (Bertrand et al., 2012).

Finally, interesting differences between cancer subtypes and phenotypes emerged when we analyzed STP coverings and tumor heterogeneity at population level. For instance, in lung cancer, source-level paired aberrations involving *KRAS* were most strongly associated with smoking, whereas those involving *EGRF* showed an opposite trend, consistent with well-established patterns (Herbst et al., 2008). Similarly, *KRAS* aberrations were virtually absent in the CRIS-C colon cancer subtype, which was in turn enriched for aberrations involving *TP53*, as previously described (Isella et al., 2017). *CHEK1* in breast cancer is also a notable regulator of the response to DNA damage, which is over-expressed in triple-negative breast cancer (TNBC) and has therefore been proposed as a potential target for treatment (Marzio et al., 2019). Notably, we were not only able to confirm *CHEK1* aberration (with over-expression) in basal-like tumors (which are enriched for TNBC), but also reveal this aberration in the luminal B subtype, suggesting a possible vulnerability of this more aggressive type of breast cancer.

### 4.2 Heterogeneity

Regarding significant differences in the computed entropy (of the joint distribution of aberrations) between tumor groups, larger entropy estimates were typically associated with more severe disease phenotypes. This suggests that there is more diversity or variation in the DNA and RNA profiles of sub-populations of patients with more aggressive disease phenotypes. Such heterogeneity observed at the population level probably reflects the variability present at the individual level—*i.e.*, the intra-tumoral heterogeneity, stemming from genetic, epigenetic, and cellular variation—which is a well-known factor impacting clinical outcome and therapy response (Jamal-Hanjani et al., 2015; Marusyk et al., 2020). Also notably, aggressive phenotypes are covered more efficiently than are less aggressive ones. This may be reflective of more aggressive phenotypes accumulating many more aberrations over time.

Large inter-patient variations in the genetic aberration profiles have been reported for individuals with the same diagnosis. Due to this heterogeneity combined with small sample sizes, it is difficult to associate specific changes in gene expression with specific aberrations in cancer genomes. In a recent pan-cancer study (PCAWG Transcriptome Core Group et al., 2020), the authors use matched whole-genome DNA and RNA sequencing data for 1,188 patients in order to identify co-occurrent DNA and RNA aberrations focusing on fusions, copy number changes, and mutation-driven aberrant splicing, followed by putative causal or mechanistic explanations. In contrast we leverage mechanistic constraints via prior, independent, biological information.

One finding with some commonality is the significant correlation between DNA and RNA alterations, observed in PCAWG Transcriptome Core Group et al. (2020). Results on our end can be seen from the values of *P* (upstream DNA|RNA) (see Tables 5, S15–S19). In our framework, STPs can and usually do involve different partners and therefore our probabilities are not strictly comparable to those of PCAWG. Thus for us, “RNA”in *P* (upstream DNA|RNA) refers to RNA-aberration in a fixed target gene and “upstream DNA”means that the target gene is linked (forms an STP) with *some* DNA-aberrant gene. Thefore a larger set of explanations is avaibale for a given RNA aberration.

A multivariate statistical approach is seen in Osmanbeyoglu et al. (2017), where the authors first predict gene expression from (phospho)protein expression and gene-specific transcription factor (TF) binding sites using affinity regression, then predict TF and protein activities from somatic changes. Biological analysis centers on specific genes and pathways, notably the dysregulating effect on TFs of activating mutations in the *PIK3CA* pathway. This pathway also emerges as pivotal in our results: indeed, the most common STPs in breast cancer are *PIK3CA* ⇒ *S100B* and *PIK3CA* ⇒ *MMP13* (see Table 1); *PIK3CA* is one of only three core source genes appearing in at least three tissues (see Figure 3); and in breast cancer, *PIK3CA* is virtually certain to have a downstream RNA aberrant target (see Table S10). Whereas the methods here and in Osmanbeyoglu et al. (2017) are largely non-overlapping, the spotlight falls on many of the same DNA-RNA associations.

(Cai et al., 2019) applies sample-specific Bayesian inferences to somatic genomic alteration and differential gene expression data to identify driver genes. Characterizations of cancer types are then obtained by summarizing the discovered relationships at the sample and the population levels. Whereas this program bears similarities with ours the objectives and methodology are quite different: our analysis is top-down, based on applying a known network to directly characterize a tumor population with a relatively concise set of paired genomic-transcriptomic relationships, and designed for quantification of inter-tumor heterogeneity. In contrast, the approach in (Cai et al., 2019) is model-driven and the networks are learned.

### 4.3 Limitations and Extensions

At the DNA level, we have only considered non-synonymous somatic mutations and extreme copy number variations. Specifically, we did not consider any annotation for mutations (*e.g.*, specific base changes) beyond population frequencies. We have also limited our annotation of downstream effects to the presence or absence of deviation of RNA expression from a baseline (normal) population, not accounting for the direction of the aberration from baseline (*i.e.*, up-regulation or down-regulation). The simplicity of such a binarized representation has enabled new findings and allows for some analysis of mechanism. Examination of source level paired aberrations facilitate the process of annotating mutations with an unknown effect (so-called variants of uncertain significance). Mutations that are recovered at the population level together with copy number losses can be presumed to be inactivating, and vice-versa.

Further consistency constraints may be imposed for a deeper analysis of the biology. For example, an STP may appear to be “inconsistent”if the source gene is duplicated or has an activating mutation and yet the target gene is down-aberrant, assuming the intermediate genes do not further modulate signaling propagation through the network. Such situations should clearly not be excluded. Needless to say there are many other explanations for such observations, *e.g.*, methylation to take but one example. Indeed, there are many cases of aberrant target genes which do not appear in any motif, *i.e.*, for which there is no putative explanation in terms of upstream mutations and copy number variations. Uncovering a mechanistically coherent picture of the upstream-downstream synergy would evidently require incorporating additional types of data (*e.g.*, gene fusions, histone modifications and changes in methylation), other sources of transcriptional dysregulation (*e.g.*, expression of microRNAs) and other downstream effects, such as post-transcriptional changes in regulation and aberrant protein structure and concentration. Without such data, making assumptions about consistency among the catalogued and detected anomalies would result in damaging over-simplifications.

The covering signatures vary considerably from one tissue or subtype to another. For instance, among our six tissues and at any level (source, target, or pair) the core set of features (those shared by all minimal coverings) is the smallest in colon and accounts for all colon samples in TCGA, whereas substantially larger signatures were necessary in other tumors (*e.g.*, in breast), and some populations could not even be largely covered (*e.g.*, in prostate) regardless of the number of features. On one hand, a plausible bias here is that better and more refined network information is available for some cancer types than others. For instance, colon cancer has served for years as a model of tumorigenesis, and a wealth of data is available to derive “realistic”signaling pathways and regulatory networks compared with cancer types studied to a lesser extent. On the other hand other DNA alterations, beside mutations and copy number changes, can drive tumorigenesis and may be necessary to efficiently “cover”a cancer population. A prototypical example is that the exceptionally large coverings in prostate cancer may be due to the absence of data on gene fusions; in fact, over a half of the tumors could be accounted for by a small subset of such alterations (*e.g.*, the fusion between *TMPRSS2* and *ERG*, or other *ETS* family genes (Tomlins et al., 2009)).

Finally, the theoretical framework we have developed is based on the “ *Regulators* → *Targets*”paradigm and would support the incorporation of additional *omics* information. In fact, gene fusion, epigenetic measurements, epigenomic states, enhancer expression, and so forth, could be simply integrated to generate an expanded repertoire of STPs, as could proteomics or metabolomics serve as additional downstream “targets”. To this end, epigenetics events (*e.g.*, methylation status, chromatin modification marks, and so on) could be easily integrated with DNA aberrations at the “source”level, while protein levels could be combined with RNA measurements. In both cases, a set of mechanistic rules would be required to integrate the different data types. For instance, a specific “source”gene in a pair could be defined as “aberrant”if it is mutated, OR it is deleted, OR it is hyper-methylated, and so on. Similarly, a “target”gene could be deemed “aberrant”based on biologically justified rules for combining protein and RNA data. Importantly, from the computational point of view, adding further modalities would not change the number of constraints in the optimization.

## 5 Conclusions

We have described an integrated analysis of DNA and RNA aberrations, which is grounded in cancer biology and enabled by a highly simplified summary representation of the complex and heterogeneous landscape of aberrations in cancer populations. The summary is a collection of STPs, each linking a particular DNA aberration with a downstream RNA expression change, and derived automatically from a “covering”algorithm in combinatorial optimization. Beside recapitulating many known alterations, our collection of STPs flags potentially important aberrations and interactions which might go unrecognized using simple frequency criteria, given the accumulation of low frequency events at the population level. This integrated representation could facilitate discriminating cancer drivers from passenger aberrations, and suggest potential novel therapeutic targets for further functional studies. Furthermore, this representation allows for a rigorous quantitative estimate of heterogeneity in a cancer population and across distinct tumor phenotypes, which would not be otherwise feasible. Indeed, in order to quantify heterogeneity beyond a simple listing of possibilities, it is necessary to assign likelihoods to these possibilities and their co-occurrences, in which case the entropy of the distributions over the possible combinations is the natural measure. The heterogeneity differences observed between distinct cancer phenotypes, along with the interactions among paired aberrations, suggest that our approach can represent an alternative to standard statistical filtering to identify important features for predictive model building and machine learning application in cancer. Finally, our analytical framework provides a highly efficient and innovative computational tool for harnessing the expanding data on tumor samples emerging from large consortia projects.

## Supporting information

paper-supplement

## 6 Supporting Information Legends

### supplementaryMaterial.pdf

The file named “supplementaryMaterial.pdf" contains all supplementary figures and tables referenced from the main paper. Specifically, this supporting file contains two subsections. In the Supplementary Table subsection, Tables S1-S2 show the basic statistics of interactions before and after filters. Tables S4-S19 illustrate the statistics of “Optimal Covering" and core set with associated probabilities for 5 tissues at 3 levels. Tables S20-S25 display the divergence probabilities at different cancer subtypes. Tables S26-S27 show the entropy analysis across distinct tissues. In Supplementary Figure subsection, Figures S1-S5 show pair covering network across different cancers. Figures S6-S8 show annotated network for cancer subtypes. Figures S10-S11 show the complete core set across tissues. Figures S12-S14 display coding trees for cancer subtypes.

### supplementaryData.zip

The compressed archive named “supplementaryData.zip”contains 6 “DNABinary.txt.gz”file which correspond to binary DNA aberration matrices for all tumor types, 6 “RN-ABinary.txt.gz”file which corresponds to binary RNA aberration matrices for all tumor types, and “full_signature.xlsx”which contains one full signature for each tumor type at each level (pair, source, target).

## Acknowledgments

We thank Drs. Giovanni Parmigiani, Nathan Price, Diego Fernando Sanchez and Eddie Luidy-Imada for helpful discussions. This research was supported by NIH National Cancer Institute Grant R01CA200859.

