## Supplementary material for "Efficient Representations of Tumor Diversity with Paired DNA-RNA Aberrations": paper-supplement

### 1 Supplementary Text

## 1.1 $k = 3$ v.s $k = 4$

Recall from Section 2.2.2, we described the process of building source-target pairs (STPs) using the Reactome pathway database, and we use $k = 3$ in our experiments where $k$ is the length of STP $g_s \Rightarrow g_t$.

Preliminary computations show that with $k > 4$, the number of matched STPs is quite large and provokes computational problems, while with $k = 1$ or $k = 2$, STPs of possible significance may be omitted. Therefore we may consider $k = 3$ and $k = 4$ to be candidates for this hyper-parameter for further consideration.

Proceeding from these initial conclusions, we now offer a more detailed analysis between $k = 3$ and $k = 4$. As seen in Table S1, the number of interactions with $k \leq 4$ is about 5 times that with $k \leq 3$. Also, the optimal covering statistics for multiple tissues (e.g., breast, colon, liver, lung) are quite similar at all three levels (i.e., STP, source, target) between Table 1 and Table 2 . Moreover, for target level covering with $k = 4$, Supplementary Figure 1 indicates that given a target gene $g_t$ is RNA aberrant, there is strong likelihood that at least one of its sources $g_s$ is DNA aberrant, with $P(\text{upstream DNA}|\text{RNA})$ between $0.7 - 1$ for indicated 4 tissues. For colon cancer and lung cancer at target level, $P(\text{upstream DNA}|\text{RNA})$ is even higher than 0.9 for all core. These relatively high values of $P(\text{upstream DNA}|\text{RNA})$ suggest that we could very likely be able to discover an upstream DNA aberration for any downstream RNA aberration – which may not be reasonable, since a gene can be RNA-aberrant for many reasons other than an upstream genetic aberration (consider for instance, an epigenetic cause or fusion-related). This would suggest $k = 3$ to be the optimal choice providing the most biologically meaningful and reasonable tableau of paired aberrations as input for the minimal covering optimization.

| Tissue | Samples | Covering Type | Quantity | Size of solution | Total features in solution | Fraction of samples covered | Size of core set |
|---|---|---|---|---|---|---|---|
| Breast | 953 | STP | 12890 | 73 | 563 | 0.977 | 16 |
| | | Source | 1133 | 59 | 95 | 0.983 | 42 |
| | | Target | 285 | 45 | 75 | 0.983 | 32 |
| Colon | 207 | STP | 4350 | 7 | 78 | 1.000 | 3 |
| | | Source | 1138 | 7 | 220 | 1.000 | 2 |
| | | Target | 275 | 8 | 137 | 1.000 | 2 |
| Kidney | 336 | STP | 857 | 41 | 113 | 0.929 | 24 |
| | | Source | 232 | 43 | 68 | 0.973 | 33 |
| | | Target | 276 | 53 | 92 | 0.991 | 28 |
| Liver | 360 | STP | 3438 | 36 | 167 | 0.978 | 11 |
| | | Source | 866 | 34 | 149 | 0.992 | 12 |
| | | Target | 347 | 32 | 125 | 0.983 | 10 |
| Lung | 465 | STP | 10291 | 22 | 87 | 0.991 | 6 |
| | | Source | 1698 | 23 | 23 | 0.996 | 23 |
| | | Target | 373 | 21 | 75 | 0.994 | 5 |
| Prostate | 491 | STP | 1977 | 64 | 119 | 0.827 | 43 |
| | | Source | 571 | 67 | 115 | 0.890 | 49 |
| | | Target | 236 | 66 | 97 | 0.845 | 49 |

Table 1: **Statistics of optimal coverings with $k = 4$.** This table shows the statistics of "Optimal Covering" at 3 levels: "STP", "Source (with target)", and "Target (with source)" for the indicated tissues. The quantity is the number of features which passed the 2% filter at the indicated level (e.g. STP, source and target.). After setting the numbers of solutions limit up to $10,000$ in the optimization program, the number of optimal coverings of each type for each tissue is reported. For instance, for breast cancer at the "STP" level, there are 12890 candidate STPs after 2 step filters, each optimal solution contains 73 STPs, 563 STPs involved in all solutions, and about 97.7% of breast cancer samples can be covered by every optimal covering. Finally, there are 16 core STPs that exist in every covering. Similar statistics are reported for all other tissue types considered in this study.

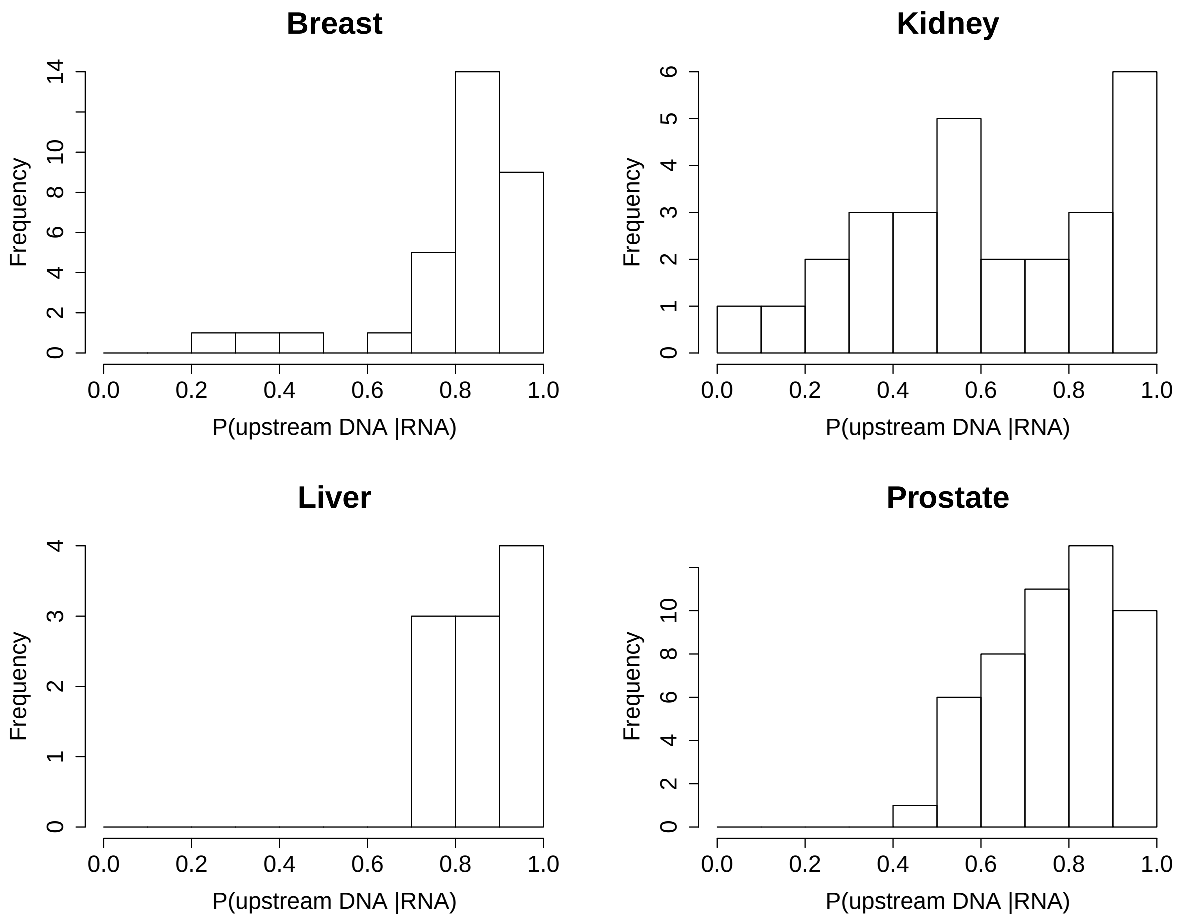

Figure 1: $P$(**upstream DNA|RNA**) **for target core across tissues** This figure shows $P$(upstream DNA|RNA) for "core" at target level of 4 tissues (Breast, Kidney, Liver, and Prostate), where $P$(upstream DNA|RNA) is the fraction of samples with indicated target gene RNA-aberrant for which there exists some DNA-aberrant gene among its source. For all four tissues in the figure, $P$(upstream DNA|RNA) is usually large as between $0.7 - 1$.

#### 1.2 Filters: 1% v.s 2% v.s 5%

Having settled on the choice of $k = 3$, we now proceed to consider the filter which omits rare events.

Recall from Section 3.2, we applied a 2% filter to remove rare events at each of three levels (STP, source, target); that is, the remaining STPs (respectively for sources, targets) after this procedure are aberrant in at least 2% of samples for a given tissue. Generally speaking, the selection of 2% is due to limitations of sample size. We usually have hundreds of samples per tissue, thus each STP (source, target) remaining after such a 2% cutoff is aberrant for at least 2 samples and usually more.

Here we conduct a further analysis and compare results between different percentage values (e.g., $1\%, 2\%, 5\%$) for this filter. Supplementary Figure 2 demonstrates our findings, indicating that a 2% cutoff is a moderate choice based on the resulting covering statistics. For instance, with a 2% filter at source level, the size of optimal coverings is usually between 10-60, and the fraction of samples covered is above 90% for most tissues (Table S4). Further, the core set contains both well-known cancer genes and potential cancer related genes (see Tables S10 -S14).

A thresholding at 1% on the other hand, even though the fraction of samples covered is higher than 2%, the core set contains many STPs (sources, targets) which are relatively more cumbersome to analyze since these STPs (sources, targets) are only aberrant for a small fraction of samples. For instance, in breast cancer with a 1% filter, about 29% of core STPs are aberrant for less than 2% of the samples and 35% of core sources are aberrant with some target for less than 2% of the samples. Further, if we use a 5% filter, while we can usually find those STPs (sources, target) which are well-known to be of interest in cancer biology and pathology, the fraction of samples covered decreases considerably. For instance in prostate cancer, with a 5% filter only 56.8% ( 64%, 63%) of samples are covered by the optimal covering at STP (source, target) level correspondingly. Together these statistics suggest a cutoff of 2% as the most reasonable choice.

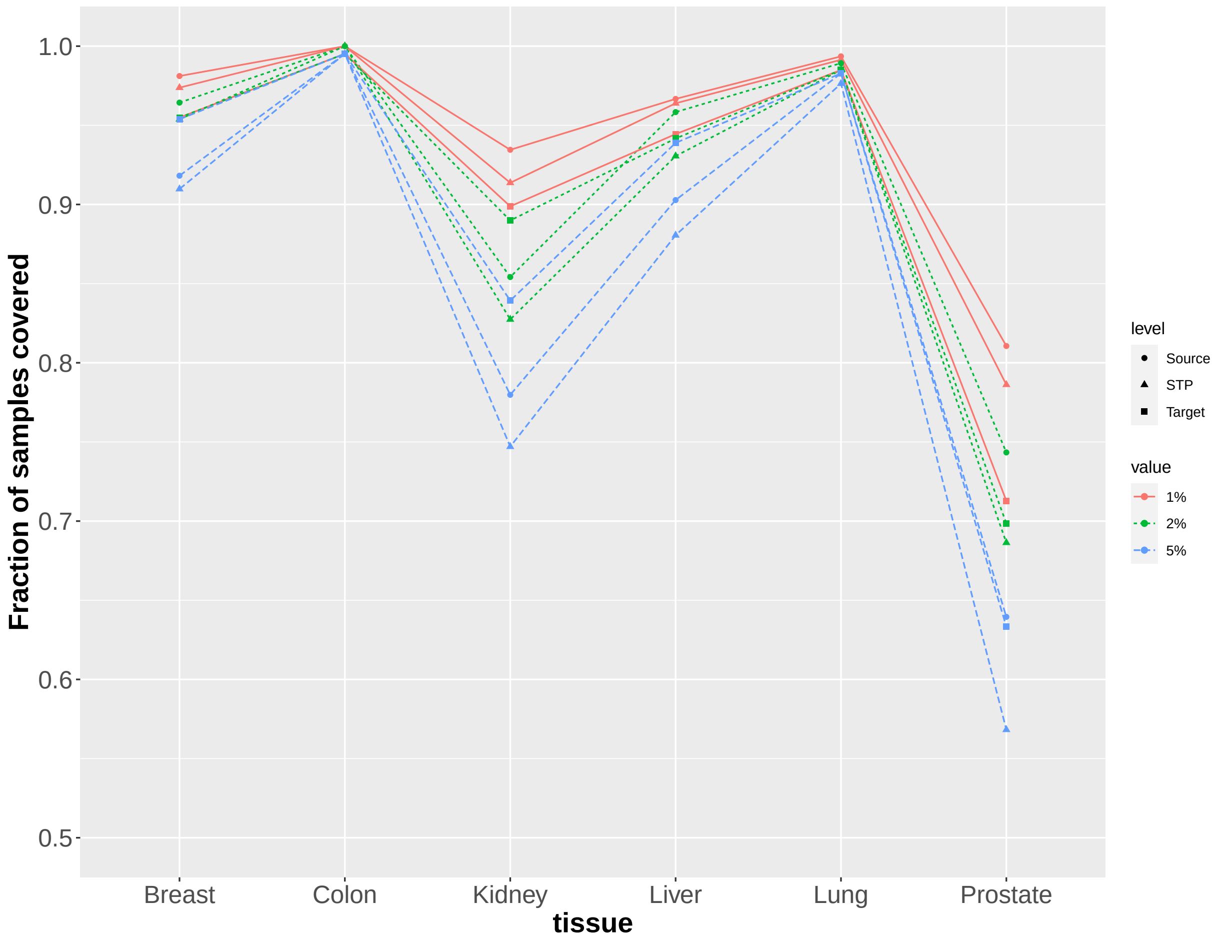

Figure 2: **Fraction of samples covered for different filter values.** This figure shows the fraction of samples covered for different filter values at indicated tissues and levels (STP, Source, Target). There are 3 candidates values: $1\%, 2\%, 5\%$. For fixed tissue and level, as we increase the filter value, the fraction of samples covered decreases.

### 1.3 $L = 4$ v.s $L = 5$

Recall from Section 2.4, we described a greedy aggregation procedure to estimate an upper bound of the entropy of $\bar{Z} = (Z_j, j \in J)$, there is one parameter $L$ which is the maximal size of subsets that partition $J$. And we use $L = 4$ in our experiments. Our criterion here is that we require at least $2^L$ samples for reliable estimations. With $L$ variables, there are $2^L$ parameters to estimate in computing entropy; requiring $2^L$ samples is then a minimum. It's also appropriate to use $L = 5$ if the sample size is reasonable. For instance, in breast cancer, the estimated entropy statistics for source level (source aberration with target) with $L = 4$ and $L = 5$ are very similar as shown in Table 2. We see that the entropy estimated with $L = 4$ is within the range of confidence interval estimated with $L = 5$, and vise versa. However, for some other cancer subtypes like Tumor Status T2 of colon cancer, we only have 28 samples for which it is not reliable to estimate entropy with $L = 5$. For consistency and accuracy, we use $L = 4$ for all cancer subtypes.

**Breast**

| Subtype | Value | N | L=4 | | L=5 | |
|---|---|---|---|---|---|---|
| | | | Entropy | Conf.Interval | Entropy | Conf.Interval |
| All | | 953 | 17.02 | [16.63, 17.40] | 16.84 | [16.43, 17.23] |
| PAM50 | Basal-like | 87 | 22.59 | [21.34, 23.69] | 22.07 | [20.89, 23.13] |
| PAM50 | HER2-enriched | 55 | 18.95 | [17.52, 20.14] | 18.42 | [17.10, 19.48] |
| PAM50 | Luminal A | 219 | 12.72 | [11.96, 13.38] | 12.44 | [11.67, 13.13] |
| PAM50 | Luminal B | 121 | 17.43 | [16.45, 18.34] | 17.11 | [16.15, 18.05] |
| ER Status | Negative | 167 | 22.39 | [21.36, 23.25] | 22.06 | [21.18, 22.95] |
| ER Status | Positive | 569 | 15.51 | [15.04, 15.97] | 15.27 | [14.81, 15.75] |
| Tumor Status | T1 | 249 | 13.56 | [12.88, 14.21] | 13.25 | [12.59, 13.89] |
| Tumor Status | T2 | 551 | 18.25 | [17.72, 18.74] | 18.04 | [17.54, 18.52] |
| Tumor Status | T3-T4 | 152 | 16.42 | [15.53, 17.26] | 16.13 | [15.24, 16.95] |
| Lymph Node Status | Negative | 446 | 17.35 | [16.76, 17.89] | 17.15 | [16.57, 17.67] |
| Lymph Node Status | Positive | 318 | 16.11 | [15.45, 16.75] | 15.89 | [15.22, 16.52] |

Table 2: **Entropy for source aberration with target with $L = 4$ and $L = 5$.** This table provides detailed entropy statistics estimated with $L = 4$ and $L = 5$ for Breast cancer at source level. The estimated entropy statistics with $L = 4$ and $L = 5$ are very similar.

#### 1.4 STP formation with Reactome

Here we present some further information regarding our use of Reactome for building STPs.

Reactome is a curated, open-source relational database of biological pathways and processes, encompassing proteins, metabolites, and their relationships. Overall, the Reactome network accounts for 336,214 interactions of 12 different types, occurring between 12,085 nodes (genes, proteins, complexes, and metabolites), which participate in 6,352 biological pathways. For our work, we restricted the analysis to gene regulatory motifs, excluding all metabolites, their modification, and their interactions. This resulted in a network encompassing a total of 110,686 interactions of 2 types ("controls state change of" and "controls expression of"), occurring between 6,725 genes participating in 1,400 biological pathways and processes. Source-target pairs were built from this network as explained in Section 2.2.2

#### 1.5 Aberrations belonging to the cores

Our observations in the core sets re-iterate some know and expected facets of cancer biology. For example, *PTEN* (inactivation or deletion) Worby and Dixon (2014) appears in the core signatures for breast, kidney, and prostate cancers, *PIK3CA* (mostly by mutation) Arafeh and Samuels (2019) was found in breast and colon cancer, while *BRCA1* Turner et al. (2004) was inactivated in breast and liver cancer. In addition, a number of aberrations affecting important cancer genes proved to be restricted to specific tissue of origin according to cell lineage and identity, and to canonical pathways of pathogenesis. For instance, *GATA3*, a transcription factor required for normal mammary gland development, and known to be altered in breast tumors Eeckhoute et al. (2007), was only found in the core signature for breast cancer. Similarly, *APC*, the tumor suppressor gene mutated in familial colon cancer and most sporadic cases of the disease Goss and Groden (2000), was present only in the colon cancer core set. Finally, among other known cancer drivers, *VHL*, for instance, was part of the core set only in kidney cancer Dizman et al. (2020), *EGFR* and *BRAF* were recovered in lung cancer Oberndorfer and Müllauer (2018), and *CDKNB1* Dong (2006) and *FOXA1* Arora and Barbieri (2018); Cancer Genome Atlas Research Network (2015) in prostate cancer, according to expectations in line with known cancer biology and recent extensive analyses ICGC/TCGA Pan-Cancer Analysis of Whole Genomes Consortium (2020); Martincorena et al. (2017).

Among the core target set, *FABP4* is a fatty acid binding protein playing an important role in glucose and lipid homeostasis Cao et al. (2008) and has aberrant expression (with an upstream DNA alteration) in all cancer types except colon and lung, suggesting that the expression of this gene could serve as cancer biomarker. Similarly, and unexpectedly, several well-known cancer drivers, including *BRCA1, MSH2, TOP2A* and *EZH2*, were recovered among the core target genes. For instance, *EZH2* is a core target in prostate cancer Varambally et al. (2002) and *S100B* is one in breast tumors Charmsaz et al. (2017).

#### 1.6 Supplementary Tables

| | Interactions | Source Genes | Target Genes | Distinct Genes |
|---|---|---|---|---|
| $k = 1$ | 3730 | 510 | 598 | 1016 |
| $k \leq 2$ | 34955 | 1915 | 598 | 2320 |
| $k \leq 3$ | 272237 | 3124 | 598 | 3479 |
| $k \leq 4$ | 1322385 | 3616 | 598 | 3952 |

**Table S1: Basic statistics of interactions within $k$ steps.** This table shows the number of interactions, source genes, target genes, and distinct genes within $k$ steps, as retrieved from the Reactome network. For instance, in our experiment, when we set $k \leq 3$, there are 272237 interactions in total, and there are 3124 source genes, 598 target genes, and 3479 distinct genes in total.

| Tissue | Filters | Interactions | Source Genes | Target Genes | Distinct Genes |
|---|---|---|---|---|---|
| Breast | After $\chi^2$ test | 17261 | 2130 | 421 | 2396 |
| Colon | After $\chi^2$ test | 6297 | 1646 | 354 | 1892 |
| Kidney | After $\chi^2$ test | 4870 | 1206 | 359 | 1485 |
| Liver | After $\chi^2$ test | 7232 | 1688 | 421 | 1989 |
| Lung | After $\chi^2$ test | 10908 | 1912 | 453 | 2218 |
| Prostate | After $\chi^2$ test | 9301 | 1732 | 372 | 1985 |

**Table S2: Basic statistics of interactions after $\chi^2$ test.**This table shows the number of interactions, source genes, target genes, and distinct genes after $\chi^2$ Test. The interactions remained are those which have P-value (uncorrected for multiple comparisons) less or equal than 0.05 between the binary DNA expression of source gene and ternary RNA expression of target gene.

| Tissue | Pair | $P$(DNA&RNA) | $P$(DNA mut &RNA up) | $P$(DNA mut &RNA down) | $P$(DNA cnv del &RNA up) | $P$(DNA cnv del &RNA down) | $P$(DNA cnv dup &RNA up) | $P$(DNA cnv dup &RNA down) |
|---|---|---|---|---|---|---|---|---|
| Breast | *PIK3CA* ⇒ *S100B* | 0.316 | 0.000 | 0.295 | 0.000 | 0.001 | 0.000 | 0.036 |
| Colon | *APC*⇒ *AXIN2* | 0.585 | 0.580 | 0.000 | 0.010 | 0.000 | 0.000 | 0.000 |
| Kidney | *VHL*⇒ *CA9* | 0.482 | 0.429 | 0.000 | 0.107 | 0.000 | 0.000 | 0.000 |
| Liver | *TP53*⇒ *MYBL2* | 0.308 | 0.294 | 0.000 | 0.022 | 0.000 | 0.000 | 0.000 |
| Lung | *TP53*⇒ *TOP2A* | 0.529 | 0.527 | 0.000 | 0.013 | 0.000 | 0.000 | 0.000 |
| Prostate | *PTEN*⇒*TWIST1* | 0.161 | 0.024 | 0.000 | 0.147 | 0.000 | 0.000 | 0.000 |

**Table S3: Examples of STPs.** For each of six tissues, one example of a common STP $\lambda = (g \Rightarrow g')$ is shown. $P$(DNA&RNA) is the fraction of samples for which the indicated source gene is DNA-aberrant and indicated target gene is RNA-aberrant. $P$(DNA mut&RNA up) is the fraction of samples for which the indicated source gene is mutated and indicated target gene is over-expressed. $P$(DNA mut&RNA down) is the fraction of samples for which the indicated source gene is mutated and indicated target gene is under-expressed. $P$(DNA cnv det&RNA up) is the fraction of samples for which the indicated source gene has copy number deleted and target gene over-expressed. $P$(DNA cnv det&RNA down) is the fraction of samples for which the indicated source gene has copy number deleted and target gene under-expressed. $P$(DNA cnv dup&RNA up) is the fraction of samples for which the indicated source gene has copy number duplicated or more and target gene over-expressed. $P$(DNA cnv dup&RNA down) is the fraction of samples for which the indicated source gene has copy number duplicated or more and target gene under-expressed. To be noted that the sum of last 6 columns does not necessarily equal to $P$(DNA&RNA), since there could be two different type of aberrations occurring at same source genes for a fixed sample.

| Tissue | Samples | Covering Type | Quantity | No. of solutions (up to 100000) | Size of solution | Total features in solution | Fraction of samples covered | Size of core set | Fraction of samples covered with core set |
|---|---|---|---|---|---|---|---|---|---|
| Breast | 953 | STP | 4026 | 100000 | 67 | 281 | 0.954 | 21 | 0.709 |
| | | Source | 690 | 100000 | 60 | 127 | 0.964 | 34 | 0.908 |
| | | Target | 256 | 83712 | 53 | 87 | 0.955 | 35 | 0.917 |
| Colon | 207 | STP | 1195 | 2353 | 11 | 101 | 1.000 | 4 | 0.807 |
| | | Source | 525 | 170 | 10 | 75 | 1.000 | 5 | 0.937 |
| | | Target | 226 | 368 | 15 | 65 | 0.995 | 6 | 0.807 |
| Kidney | 336 | STP | 347 | 100000 | 26 | 67 | 0.827 | 12 | 0.732 |
| | | Source | 133 | 256 | 28 | 37 | 0.854 | 21 | 0.830 |
| | | Target | 176 | 18180 | 60 | 88 | 0.890 | 45 | 0.812 |
| Liver | 360 | STP | 1198 | 100000 | 32 | 303 | 0.931 | 11 | 0.533 |
| | | Source | 460 | 9840 | 34 | 77 | 0.958 | 20 | 0.822 |
| | | Target | 287 | 702 | 41 | 75 | 0.942 | 26 | 0.858 |
| Lung | 465 | STP | 3154 | 100000 | 27 | 110 | 0.985 | 10 | 0.712 |
| | | Source | 908 | 140 | 25 | 42 | 0.989 | 19 | 0.974 |
| | | Target | 350 | 15 | 29 | 34 | 0.985 | 26 | 0.981 |
| Prostate | 491 | STP | 430 | 100000 | 53 | 110 | 0.686 | 32 | 0.578 |
| | | Source | 211 | 2620 | 53 | 69 | 0.743 | 42 | 0.707 |
| | | Target | 160 | 192 | 72 | 81 | 0.699 | 66 | 0.686 |

**Table S4: Statistics of optimal coverings.** This table shows the statistics of "Optimal Covering" at 3 levels: "STP", "Source (with target)", and "Target (with source)" for the indicated tissues. The quantity is the number of features which passed 2% filter at the indicated level (e.g. STP, source and target.). After setting the numbers of solutions limit up to $100,000$ in the optimization program, the number of optimal coverings of each type for each tissue is reported. For instance, for breast cancer at the "STP" level, there are 4026 candidate STPs after 2 step filters, there are $100,000$ solutions found from the optimization model, each solution contains 67 STPs, 281 STPs involved in all $100,000$ solutions, and about 95.4% of breast cancer samples can be covered by every optimal covering. Finally, there are 21 core STPs that exist in every covering, and 70.9% of breast cancer samples can be covered by such core set. Similar statistics are reported for all other tissue types considered in this study.

| Pair | $P(\text{DNA}\,\&\,\text{RNA})$ | $P(\text{DNA})$ | $P(\text{RNA})$ | $P(\text{RNA}\mid\text{DNA})$ | $P(\text{DNA}\mid\text{RNA})$ |
|---|---|---|---|---|---|
| *PIK3CA* ⇒ *MMP13* | 0.222 | 0.356 | 0.574 | 0.625 | 0.388 |
| *BAG4* ⇒ *LIFR* | 0.133 | 0.148 | 0.831 | 0.901 | 0.160 |
| *GATA3* ⇒ *CDC25C* | 0.116 | 0.137 | 0.745 | 0.847 | 0.156 |
| *CLTC* ⇒ *S100B* | 0.102 | 0.111 | 0.838 | 0.915 | 0.121 |
| *CDH1* ⇒ *LIFR* | 0.100 | 0.132 | 0.831 | 0.754 | 0.120 |
| *GATA3* ⇒ *LGALS3* | 0.081 | 0.137 | 0.434 | 0.588 | 0.186 |
| *CDH1* ⇒ *KCTD6* | 0.069 | 0.132 | 0.359 | 0.524 | 0.193 |
| *PTEN* ⇒ *S100B* | 0.067 | 0.093 | 0.838 | 0.719 | 0.080 |
| *PTEN* ⇒ *FOXP3* | 0.063 | 0.093 | 0.534 | 0.674 | 0.118 |
| *GAB2* ⇒ *SOD2* | 0.048 | 0.077 | 0.508 | 0.630 | 0.095 |
| *IFT140* ⇒ *PTCH1* | 0.039 | 0.055 | 0.526 | 0.712 | 0.074 |
| *CNOT3* ⇒ *CDC25C* | 0.035 | 0.037 | 0.745 | 0.943 | 0.046 |
| *DNM3* ⇒ *MMP3* | 0.031 | 0.108 | 0.207 | 0.291 | 0.152 |
| *PLXNA4* ⇒ *CDC25C* | 0.028 | 0.029 | 0.745 | 0.964 | 0.038 |
| *MYB* ⇒ *CDC25A* | 0.028 | 0.039 | 0.509 | 0.730 | 0.056 |
| *AARS* ⇒ *ACACB* | 0.027 | 0.029 | 0.704 | 0.929 | 0.039 |
| *LAMA1* ⇒ *MMP1* | 0.026 | 0.033 | 0.540 | 0.806 | 0.049 |
| *MMP9* ⇒ *MMP1* | 0.026 | 0.036 | 0.540 | 0.735 | 0.049 |
| *PRKDC* ⇒ *OPRD1* | 0.025 | 0.088 | 0.193 | 0.286 | 0.130 |
| *MBTPS1* ⇒ *LPL* | 0.025 | 0.030 | 0.614 | 0.828 | 0.041 |
| *MUC20* ⇒ *SAA1* | 0.023 | 0.057 | 0.566 | 0.407 | 0.041 |

**Table S5: Breast core STPs.** There are twenty-one "core" STPs which appear in every minimal covering of the breast samples. $P(\text{DNA}\,\&\,\text{RNA})$ is the fraction of samples for which the source gene $g$ is DNA-aberrant and target gene $g'$ is RNA-aberrant; $P(\text{DNA})$ is the fraction of samples satisfying source gene $g$ is DNA-aberrant; $P(\text{RNA})$ is the fraction of samples with $g'$ RNA-aberrant; $P(\text{RNA}\mid\text{DNA})$ is the fraction of DNA-aberrant samples for which $g'$ is RNA-aberrant.

| Pair | $P(\text{DNA}\,\&\,\text{RNA})$ | $P(\text{DNA})$ | $P(\text{RNA})$ | $P(\text{RNA}\mid\text{DNA})$ | $P(\text{DNA}\mid\text{RNA})$ |
|---|---|---|---|---|---|
| *VHL* ⇒ *CA9* | 0.482 | 0.485 | 0.967 | 0.994 | 0.498 |
| *PBRM1* ⇒ *BCL2* | 0.223 | 0.429 | 0.446 | 0.521 | 0.500 |
| *PBRM1* ⇒ *ATM* | 0.134 | 0.429 | 0.250 | 0.312 | 0.536 |
| *SLC12A2* ⇒ *EBI3* | 0.113 | 0.164 | 0.557 | 0.691 | 0.203 |
| *BAP1* ⇒ *TP53AIP1* | 0.083 | 0.202 | 0.289 | 0.412 | 0.289 |
| *RAF1* ⇒ *HEY1* | 0.068 | 0.113 | 0.438 | 0.605 | 0.156 |
| *PBRM1* ⇒ *RAG1* | 0.057 | 0.429 | 0.220 | 0.132 | 0.257 |
| *TTC37* ⇒ *CASP1* | 0.045 | 0.051 | 0.536 | 0.882 | 0.083 |
| *ATM* ⇒ *TNFRSF10D* | 0.039 | 0.080 | 0.271 | 0.481 | 0.143 |
| *MTOR* ⇒ *ITCH* | 0.030 | 0.071 | 0.146 | 0.417 | 0.204 |
| *BAP1* ⇒ *CASP6* | 0.027 | 0.202 | 0.062 | 0.132 | 0.429 |
| *RYR2* ⇒ *GZMB* | 0.027 | 0.042 | 0.360 | 0.643 | 0.074 |

**Table S6: Kidney core STPs.** There are twelve "core" STPs which appear in every minimal covering of the kidney samples. $P(\text{DNA}\,\&\,\text{RNA})$ is the fraction of samples for which the source gene $g$ is DNA-aberrant and target gene $g'$ is RNA-aberrant; $P(\text{DNA})$ is the fraction of samples satisfying source gene $g$ is DNA-aberrant; $P(\text{RNA})$ is the fraction of samples with $g'$ RNA-aberrant; $P(\text{RNA}\mid\text{DNA})$ is the fraction of DNA-aberrant samples for which $g'$ is RNA-aberrant.

| Pair | $P$(DNA & RNA) | $P$(DNA) | $P$(RNA) | $P$(RNA\|DNA) | $P$(DNA\|RNA) |
|---|---|---|---|---|---|
| *CTNNB1* ⇒ *CXCL12* | 0.261 | 0.269 | 0.897 | 0.969 | 0.291 |
| *CTNNB1* ⇒ *LCN2* | 0.136 | 0.269 | 0.383 | 0.505 | 0.355 |
| *RB1* ⇒ *CHEK1* | 0.100 | 0.111 | 0.506 | 0.900 | 0.198 |
| *BAP1* ⇒ *STEAP3* | 0.064 | 0.069 | 0.725 | 0.920 | 0.088 |
| *HERC2* ⇒ *IGFBP3* | 0.053 | 0.053 | 0.833 | 1.000 | 0.063 |
| *ALMS1* ⇒ *MSH2* | 0.042 | 0.058 | 0.567 | 0.714 | 0.074 |
| *ANK1* ⇒ *HEY2* | 0.036 | 0.078 | 0.322 | 0.464 | 0.112 |
| *BRCA1* ⇒ *TNFRSF10B* | 0.033 | 0.047 | 0.367 | 0.706 | 0.091 |
| *CDC27* ⇒ *FGF2* | 0.028 | 0.058 | 0.253 | 0.476 | 0.110 |
| *NCOR1* ⇒ *F7* | 0.028 | 0.053 | 0.411 | 0.526 | 0.068 |
| *CLTC* ⇒ *S1PR1* | 0.025 | 0.069 | 0.256 | 0.360 | 0.098 |

**Table S7: Liver core STPs.** There are eleven "core" STPs which appear in every minimal covering of the liver samples. $P$(DNA & RNA) is the fraction of samples for which the source gene $g$ is DNA-aberrant and target gene $g'$ is RNA-aberrant; $P$(DNA) is the fraction of samples satisfying source gene $g$ is DNA-aberrant; $P$(RNA) is the fraction of samples with $g'$ RNA-aberrant; $P$(RNA|DNA) is the fraction of DNA-aberrant samples for which $g'$ is RNA-aberrant.

| Pair | $P$(DNA & RNA) | $P$(DNA) | $P$(RNA) | $P$(RNA\|DNA) | $P$(DNA\|RNA) |
|---|---|---|---|---|---|
| *KRAS* ⇒ *CD36* | 0.335 | 0.351 | 0.914 | 0.957 | 0.367 |
| *ANK2* ⇒ *DLGAP5* | 0.189 | 0.196 | 0.826 | 0.967 | 0.229 |
| *EGFR* ⇒ *LPL* | 0.159 | 0.168 | 0.832 | 0.949 | 0.191 |
| *SMARCA4* ⇒ *CHEK1* | 0.092 | 0.095 | 0.766 | 0.977 | 0.121 |
| *KIF4B* ⇒ *RUNX2* | 0.058 | 0.069 | 0.613 | 0.844 | 0.095 |
| *LRRK2* ⇒ *TP53* | 0.052 | 0.069 | 0.503 | 0.750 | 0.103 |
| *ALMS1* ⇒ *PTK6* | 0.047 | 0.071 | 0.744 | 0.667 | 0.064 |
| *FRS2* ⇒ *IL23A* | 0.047 | 0.065 | 0.527 | 0.733 | 0.090 |
| *BRAF* ⇒ *CDKN1A* | 0.026 | 0.105 | 0.108 | 0.245 | 0.240 |
| *FYN* ⇒ *VIM* | 0.024 | 0.028 | 0.798 | 0.846 | 0.030 |

**Table S8: Lung core STPs.** There are ten "core" STPs which appear in every minimal covering of the lung samples. $P$(DNA & RNA) is the fraction of samples for which the source gene $g$ is DNA-aberrant and target gene $g'$ is RNA-aberrant; $P$(DNA) is the fraction of samples satisfying source gene $g$ is DNA-aberrant; $P$(RNA) is the fraction of samples with $g'$ RNA-aberrant; $P$(RNA|DNA) is the fraction of DNA-aberrant samples for which $g'$ is RNA-aberrant.

| Pair | $P$(DNA & RNA) | $P$(DNA) | $P$(RNA) | $P$(RNA\|DNA) | $P$(DNA\|RNA) |
|---|---|---|---|---|---|
| *PTEN* ⇒ *TWIST1* | 0.161 | 0.216 | 0.654 | 0.745 | 0.246 |
| *PTEN* ⇒ *LGALS3* | 0.151 | 0.216 | 0.550 | 0.698 | 0.274 |
| *FGF17* ⇒ *BNIP3L* | 0.120 | 0.169 | 0.462 | 0.711 | 0.260 |
| *HDAC2* ⇒ *HEY2* | 0.073 | 0.145 | 0.257 | 0.507 | 0.286 |
| *PTEN* ⇒ *ITGBL1* | 0.073 | 0.216 | 0.189 | 0.340 | 0.387 |
| *NRG1* ⇒ *PBX1* | 0.071 | 0.116 | 0.409 | 0.614 | 0.174 |
| *FYN* ⇒ *BOLA2* | 0.067 | 0.134 | 0.365 | 0.500 | 0.184 |
| *DVL2* ⇒ *TWIST1* | 0.057 | 0.067 | 0.654 | 0.848 | 0.087 |
| *FOXA1* ⇒ *MYL9* | 0.047 | 0.081 | 0.381 | 0.575 | 0.123 |
| *SPTA1* ⇒ *MYL9* | 0.045 | 0.063 | 0.381 | 0.710 | 0.118 |
| *DSCAM* ⇒ *BNIP3L* | 0.043 | 0.130 | 0.462 | 0.328 | 0.093 |
| *ZFHX3* ⇒ *DHFR* | 0.039 | 0.126 | 0.163 | 0.306 | 0.237 |
| *STAT3* ⇒ *TWIST1* | 0.037 | 0.039 | 0.654 | 0.947 | 0.056 |
| *PIK3CA* ⇒ *MYL9* | 0.035 | 0.061 | 0.381 | 0.567 | 0.091 |
| *SNAP91* ⇒ *PTGS2* | 0.033 | 0.126 | 0.159 | 0.258 | 0.205 |
| *TP53* ⇒ *TP53* | 0.031 | 0.179 | 0.055 | 0.170 | 0.556 |
| *DIS3* ⇒ *TNFRSF10C* | 0.029 | 0.110 | 0.106 | 0.259 | 0.269 |
| *NRG1* ⇒ *CPT1A* | 0.029 | 0.116 | 0.100 | 0.246 | 0.286 |
| *PTEN* ⇒ *MMP9* | 0.024 | 0.216 | 0.059 | 0.113 | 0.414 |
| *RAD17* ⇒ *TP53I3* | 0.024 | 0.063 | 0.141 | 0.387 | 0.174 |
| *RFC3* ⇒ *BCL2* | 0.024 | 0.071 | 0.175 | 0.343 | 0.140 |
| *MYC* ⇒ *DHFR* | 0.024 | 0.086 | 0.163 | 0.286 | 0.150 |
| *IL6ST* ⇒ *ANXA2* | 0.022 | 0.077 | 0.464 | 0.289 | 0.048 |
| *JAK1* ⇒ *BOLA2* | 0.022 | 0.037 | 0.365 | 0.611 | 0.061 |
| *PIK3CA* ⇒ *ITGBL1* | 0.022 | 0.061 | 0.189 | 0.367 | 0.118 |
| *NUP205* ⇒ *PBX1* | 0.022 | 0.022 | 0.409 | 1.000 | 0.055 |
| *ZFPM1* ⇒ *MAPKAPK5* | 0.022 | 0.094 | 0.104 | 0.239 | 0.216 |
| *CAMK2G* ⇒ *LGALS3* | 0.020 | 0.022 | 0.550 | 0.909 | 0.037 |
| *CAST* ⇒ *SCO2* | 0.020 | 0.051 | 0.189 | 0.400 | 0.108 |
| *IRS2* ⇒ *LGALS3* | 0.020 | 0.022 | 0.550 | 0.909 | 0.037 |
| *FANCD2* ⇒ *HIGD1A* | 0.020 | 0.035 | 0.210 | 0.588 | 0.097 |
| *MYC* ⇒ *BTG2* | 0.020 | 0.086 | 0.116 | 0.238 | 0.175 |

**Table S9: Prostate core STPs.** There are thirty-two "core" STPs which appear in every minimal covering of the prostate samples. $P(\text{DNA}\,\&\,\text{RNA})$ is the fraction of samples for which the source gene $g$ is DNA-aberrant and target gene $g'$ is RNA-aberrant; $P(\text{DNA})$ is the fraction of samples satisfying source gene $g$ is DNA-aberrant; $P(\text{RNA})$ is the fraction of samples with $g'$ RNA-aberrant; $P(\text{RNA}|\text{DNA})$ is the fraction of DNA-aberrant samples for which $g'$ is RNA-aberrant.

| Source | $P$(DNA & downstream RNA) | $P$(DNA) | $P$(downstream RNA\|DNA) |
|---|---|---|---|
| *PIK3CA* | 0.353 | 0.356 | 0.991 |
| *TP53* | 0.313 | 0.313 | 1.000 |
| *PTK2* | 0.164 | 0.165 | 0.994 |
| *BAG4* | 0.145 | 0.148 | 0.979 |
| *GATA3* | 0.134 | 0.137 | 0.977 |
| *CDH1* | 0.129 | 0.132 | 0.976 |
| *NCSTN* | 0.114 | 0.123 | 0.932 |
| *CLTC* | 0.111 | 0.111 | 1.000 |
| *NCOA2* | 0.103 | 0.103 | 1.000 |
| *MED1* | 0.100 | 0.101 | 0.990 |
| *PTEN* | 0.093 | 0.093 | 1.000 |
| *MUC20* | 0.056 | 0.057 | 0.981 |
| *DNM3* | 0.046 | 0.108 | 0.427 |
| *IFT140* | 0.042 | 0.055 | 0.769 |
| *BRCA1* | 0.037 | 0.038 | 0.972 |
| *CNOT3* | 0.037 | 0.037 | 1.000 |
| *ATM* | 0.036 | 0.042 | 0.850 |
| *GRIN2B* | 0.036 | 0.036 | 1.000 |
| *MMP9* | 0.035 | 0.036 | 0.971 |
| *TFDP1* | 0.035 | 0.035 | 1.000 |
| *FURIN* | 0.031 | 0.035 | 0.909 |
| *DSCAM* | 0.030 | 0.030 | 1.000 |
| *DUSP16* | 0.029 | 0.031 | 0.933 |
| *PIK3R1* | 0.029 | 0.029 | 1.000 |
| *AARS* | 0.028 | 0.029 | 0.964 |
| *PHLPP2* | 0.028 | 0.028 | 1.000 |
| *POM121* | 0.028 | 0.029 | 0.964 |
| *HIST1H3B* | 0.027 | 0.027 | 1.000 |
| *PIK3C2G* | 0.025 | 0.030 | 0.828 |
| *MBTPS1* | 0.025 | 0.030 | 0.828 |
| *PPARG* | 0.024 | 0.025 | 0.958 |
| *ITSN2* | 0.023 | 0.023 | 1.000 |
| *SLC27A5* | 0.023 | 0.030 | 0.759 |
| *FAM20C* | 0.021 | 0.021 | 1.000 |

**Table S10: Breast core source genes.** There are thirty-four "core" source genes which appear in every minimal source covering of the breast samples. $P$(DNA) is the fraction of samples for which the indicated source gene is DNA-aberrant; $P$(DNA & downstream RNA) is the fraction of samples for which the indicated source gene is DNA-aberrant and there exists an RNA-aberrant gene among its targets. $P$(downstream RNA|DNA) is the fraction of the samples with the indicated source gene DNA-aberrant for which there exists some RNA-aberrant gene among its targets.

| Source | $P(\text{DNA}\,\&\,\text{downstream RNA})$ | $P(\text{DNA})$ | $P(\text{downstream RNA}\vert\text{DNA})$ |
|---|---|---|---|
| *VHL* | 0.482 | 0.485 | 0.994 |
| *PBRM1* | 0.307 | 0.429 | 0.715 |
| *MAML1* | 0.179 | 0.179 | 1.000 |
| *NRG2* | 0.173 | 0.173 | 1.000 |
| *SLC12A2* | 0.113 | 0.164 | 0.691 |
| *CDC25C* | 0.095 | 0.167 | 0.571 |
| *RAF1* | 0.095 | 0.113 | 0.842 |
| *BAP1* | 0.092 | 0.202 | 0.456 |
| *ATM* | 0.065 | 0.080 | 0.815 |
| *MTOR* | 0.060 | 0.071 | 0.833 |
| *PTEN* | 0.048 | 0.048 | 1.000 |
| *TTC37* | 0.045 | 0.051 | 0.882 |
| *CDC27* | 0.030 | 0.057 | 0.526 |
| *RYR2* | 0.027 | 0.042 | 0.643 |
| *TP53* | 0.027 | 0.027 | 1.000 |
| *FGFR1* | 0.024 | 0.024 | 1.000 |
| *ROCK1* | 0.024 | 0.024 | 1.000 |
| *GRIN2B* | 0.021 | 0.021 | 1.000 |
| *JAK2* | 0.021 | 0.021 | 1.000 |
| *TEX15* | 0.021 | 0.021 | 1.000 |
| *TAF1* | 0.021 | 0.021 | 1.000 |

**Table S11: Kidney core source genes.** There are twenty-one "core" source genes which appear in every minimal source covering of the kidney samples. $P(\text{DNA})$ is the fraction of samples for which the indicated source gene is DNA-aberrant; $P(\text{DNA}\,\&\,\text{downstream RNA})$ is the fraction of samples for which the indicated source gene is DNA-aberrant and there exists an RNA-aberrant gene among its targets. $P(\text{downstream RNA}|\text{DNA})$ is the fraction of the samples with the indicated source gene DNA-aberrant for which there exists some RNA-aberrant gene among its targets.

| Source | $P$(DNA & downstream RNA) | $P$(DNA) | $P$(downstream RNA\|DNA) |
|---|---|---|---|
| *TP53* | 0.319 | 0.319 | 1.000 |
| *CTNNB1* | 0.269 | 0.269 | 1.000 |
| *RB1* | 0.108 | 0.111 | 0.975 |
| *NRG1* | 0.081 | 0.081 | 1.000 |
| *CCND1* | 0.075 | 0.075 | 1.000 |
| *BAP1* | 0.069 | 0.069 | 1.000 |
| *EHMT2* | 0.058 | 0.058 | 1.000 |
| *HERC2* | 0.053 | 0.053 | 1.000 |
| *PIK3CA* | 0.053 | 0.053 | 1.000 |
| *BRCA1* | 0.047 | 0.047 | 1.000 |
| *CDC27* | 0.047 | 0.058 | 0.810 |
| *ALMS1* | 0.044 | 0.058 | 0.762 |
| *IL6ST* | 0.033 | 0.036 | 0.923 |
| *CLTC* | 0.028 | 0.069 | 0.400 |
| *FANCB* | 0.028 | 0.028 | 1.000 |
| *PAN2* | 0.028 | 0.028 | 1.000 |
| *ITGB3* | 0.025 | 0.031 | 0.818 |
| *CAMK2A* | 0.022 | 0.028 | 0.800 |
| *CEP290* | 0.022 | 0.028 | 0.800 |
| *MTM1* | 0.022 | 0.028 | 0.800 |

**Table S12: Liver core source genes.** There are twenty "core" source genes which appear in every minimal source covering of the liver samples. $P$(DNA) is the fraction of samples for which the indicated source gene is DNA-aberrant; $P$(DNA & downstream RNA) is the fraction of samples for which the indicated source gene is DNA-aberrant and there exists an RNA-aberrant gene among its targets. $P$(downstream RNA|DNA) is the fraction of the samples with the indicated source gene DNA-aberrant for which there exists some RNA-aberrant gene among its targets.

| Source | $P$(DNA & downstream RNA) | $P$(DNA) | $P$(downstream RNA\|DNA) |
|---|---|---|---|
| *TP53* | 0.535 | 0.535 | 1.000 |
| *KRAS* | 0.351 | 0.351 | 1.000 |
| *SPTA1* | 0.303 | 0.303 | 1.000 |
| *ANK2* | 0.189 | 0.196 | 0.967 |
| *STK11* | 0.178 | 0.178 | 1.000 |
| *EGFR* | 0.168 | 0.168 | 1.000 |
| *RYR1* | 0.142 | 0.189 | 0.750 |
| *NUP155* | 0.127 | 0.127 | 1.000 |
| *PIK3C2B* | 0.105 | 0.110 | 0.961 |
| *SMARCA4* | 0.095 | 0.095 | 1.000 |
| *BRAF* | 0.092 | 0.105 | 0.878 |
| *MET* | 0.084 | 0.084 | 1.000 |
| *VWF* | 0.082 | 0.084 | 0.974 |
| *ALMS1* | 0.069 | 0.071 | 0.970 |
| *LRRK2* | 0.067 | 0.069 | 0.969 |
| *MTMR9* | 0.065 | 0.069 | 0.938 |
| *LAMB2* | 0.030 | 0.034 | 0.875 |
| *FYN* | 0.028 | 0.028 | 1.000 |
| *RAD50* | 0.024 | 0.026 | 0.917 |

**Table S13: Lung core source genes.** There are nineteen "core" source genes which appear in every minimal source covering of the lung samples. $P$(DNA) is the fraction of samples for which the indicated source gene is DNA-aberrant; $P$(DNA & downstream RNA) is the fraction of samples for which the indicated source gene is DNA-aberrant and there exists an RNA-aberrant gene among its targets. $P$(downstream RNA|DNA) is the fraction of the samples with the indicated source gene DNA-aberrant for which there exists some RNA-aberrant gene among its targets.

| Source | $P$(DNA & downstream RNA) | $P$(DNA) | $P$(downstream RNA\|DNA) |
|---|---|---|---|
| *PTEN* | 0.210 | 0.216 | 0.972 |
| *FGF17* | 0.128 | 0.169 | 0.759 |
| *TP53* | 0.128 | 0.179 | 0.716 |
| *FYN* | 0.118 | 0.134 | 0.879 |
| *NRG1* | 0.112 | 0.116 | 0.965 |
| *SNAP91* | 0.102 | 0.126 | 0.806 |
| *ZFHX3* | 0.092 | 0.126 | 0.726 |
| *MYC* | 0.086 | 0.086 | 1.000 |
| *DIS3* | 0.069 | 0.110 | 0.630 |
| *FOXA1* | 0.067 | 0.081 | 0.825 |
| *IL6ST* | 0.065 | 0.077 | 0.842 |
| *NCOA2* | 0.059 | 0.063 | 0.935 |
| *ITGA2B* | 0.055 | 0.067 | 0.818 |
| *LYN* | 0.049 | 0.051 | 0.960 |
| *PDE7A* | 0.049 | 0.055 | 0.889 |
| *PIK3CA* | 0.049 | 0.061 | 0.800 |
| *RAD17* | 0.043 | 0.063 | 0.677 |
| *TSLP* | 0.041 | 0.049 | 0.833 |
| *POLK* | 0.039 | 0.051 | 0.760 |
| *ZFPM1* | 0.037 | 0.094 | 0.391 |
| *STAT3* | 0.037 | 0.039 | 0.947 |
| *JAK1* | 0.035 | 0.037 | 0.944 |
| *CDH1* | 0.033 | 0.059 | 0.552 |
| *KL* | 0.033 | 0.069 | 0.471 |
| *RFC3* | 0.033 | 0.071 | 0.457 |
| *CTNNB1* | 0.031 | 0.031 | 1.000 |
| *DYNC1H1* | 0.031 | 0.031 | 1.000 |
| *E2F4* | 0.031 | 0.035 | 0.882 |
| *ERBB4* | 0.029 | 0.033 | 0.875 |
| *CCNH* | 0.026 | 0.043 | 0.619 |
| *SGIP1* | 0.024 | 0.026 | 0.923 |
| *ACTN2* | 0.022 | 0.035 | 0.647 |
| *HSP90AA1* | 0.022 | 0.022 | 1.000 |
| *CAMK2G* | 0.020 | 0.022 | 0.909 |
| *CDK5RAP2* | 0.020 | 0.022 | 0.909 |
| *CDKN1B* | 0.020 | 0.069 | 0.294 |
| *GNG12* | 0.020 | 0.024 | 0.833 |
| *IL33* | 0.020 | 0.024 | 0.833 |
| *IL6R* | 0.020 | 0.020 | 1.000 |
| *IRS2* | 0.020 | 0.022 | 0.909 |
| *LRP5* | 0.020 | 0.022 | 0.909 |
| *NUP160* | 0.020 | 0.026 | 0.769 |

**Table S14: Prostate core source genes.** There are forty-two "core" source genes which appear in every minimal source covering of the prostate samples. $P$(DNA) is the fraction of samples for which the indicated source gene is DNA-aberrant; $P$(DNA & downstream RNA) is the fraction of samples for which the indicated source gene is DNA-aberrant and there exists an RNA-aberrant gene among its targets. $P$(downstream RNA|DNA) is the fraction of the samples with the indicated source gene DNA-aberrant for which there exists some RNA-aberrant gene among its targets.

| Target | $P$(RNA & upstream DNA) | $P$(RNA) | $P$(upstream DNA\|RNA) |
|---|---|---|---|
| *CDC25C* | 0.706 | 0.745 | 0.948 |
| *S100B* | 0.659 | 0.838 | 0.786 |
| *CDC25A* | 0.489 | 0.509 | 0.961 |
| *CXCL12* | 0.483 | 0.627 | 0.769 |
| *MMP1* | 0.467 | 0.540 | 0.864 |
| *PIK3R1* | 0.463 | 0.524 | 0.884 |
| *FOS* | 0.445 | 0.507 | 0.878 |
| *MMP13* | 0.441 | 0.574 | 0.768 |
| *FABP4* | 0.424 | 0.663 | 0.639 |
| *CD36* | 0.421 | 0.793 | 0.530 |
| *FOXO1* | 0.416 | 0.687 | 0.605 |
| *BRCA1* | 0.410 | 0.460 | 0.893 |
| *RHOU* | 0.410 | 0.663 | 0.619 |
| *BCL6* | 0.393 | 0.576 | 0.683 |
| *PLAGL1* | 0.376 | 0.535 | 0.702 |
| *LIFR* | 0.367 | 0.831 | 0.442 |
| *S1PR1* | 0.347 | 0.409 | 0.849 |
| *E2F7* | 0.347 | 0.362 | 0.959 |
| *SOCS2* | 0.341 | 0.437 | 0.781 |
| *MYL9* | 0.292 | 0.523 | 0.558 |
| *PTCH1* | 0.285 | 0.526 | 0.543 |
| *CASP1* | 0.282 | 0.366 | 0.771 |
| *CASP6* | 0.247 | 0.375 | 0.658 |
| *VIM* | 0.240 | 0.346 | 0.694 |
| *KCTD6* | 0.232 | 0.359 | 0.646 |
| *IFNB1* | 0.230 | 0.346 | 0.664 |
| *JAG1* | 0.220 | 0.319 | 0.691 |
| *MLH1* | 0.195 | 0.242 | 0.805 |
| *EPO* | 0.195 | 0.267 | 0.732 |
| *PTK6* | 0.172 | 0.370 | 0.465 |
| *RUNX2* | 0.159 | 0.255 | 0.626 |
| *RAG1* | 0.095 | 0.191 | 0.500 |
| *ITGAL* | 0.095 | 0.183 | 0.523 |
| *IFNA10* | 0.050 | 0.057 | 0.889 |
| *HIGD1A* | 0.045 | 0.084 | 0.537 |

**Table S15: Breast core target genes.** There are thirty-five "core" target genes which appear in every minimal target covering of the breast samples. $P$(RNA) is the fraction of samples for which the indicated target gene is RNA-aberrant; $P$(RNA & upstream DNA) is the fraction of samples for which the indicated target gene is RNA-aberrant and there exists an DNA-aberrant gene among its sources. $P$(upstream DNA|RNA) is the fraction of the samples with the indicated gene RNA-aberrant for which at least one of its sources is DNA-aberrant.

| Target | $P$(RNA & upstream DNA) | $P$(RNA) | $P$(upstream DNA\|RNA) |
|---|---|---|---|
| *CA9* | 0.503 | 0.967 | 0.520 |
| *HIGD1A* | 0.473 | 0.929 | 0.510 |
| *BCL2* | 0.390 | 0.446 | 0.873 |
| *FABP7* | 0.223 | 0.881 | 0.253 |
| *EBI3* | 0.199 | 0.557 | 0.358 |
| *BAX* | 0.170 | 0.429 | 0.396 |
| *ATM* | 0.170 | 0.250 | 0.679 |
| *NDN* | 0.146 | 0.280 | 0.521 |
| *STEAP3* | 0.146 | 0.250 | 0.583 |
| *TP53AIP1* | 0.131 | 0.289 | 0.454 |
| *BBC3* | 0.122 | 0.277 | 0.441 |
| *HEY1* | 0.113 | 0.438 | 0.259 |
| *LIFR* | 0.092 | 0.402 | 0.230 |
| *TGFA* | 0.080 | 0.646 | 0.124 |
| *TNFRSF10D* | 0.080 | 0.271 | 0.297 |
| *AXIN1* | 0.080 | 0.268 | 0.300 |
| *BRCA1* | 0.080 | 0.280 | 0.287 |
| *FANCI* | 0.074 | 0.339 | 0.219 |
| *AIFM2* | 0.071 | 0.098 | 0.727 |
| *POU2F1* | 0.068 | 0.098 | 0.697 |
| *ZEB1* | 0.065 | 0.152 | 0.431 |
| *DDB2* | 0.065 | 0.964 | 0.068 |
| *MMP9* | 0.065 | 0.384 | 0.171 |
| *APAF1* | 0.060 | 0.089 | 0.667 |
| *FABP4* | 0.060 | 0.074 | 0.800 |
| *BRWD1* | 0.054 | 0.077 | 0.692 |
| *FOXO3* | 0.054 | 0.107 | 0.500 |
| *FGF2* | 0.051 | 0.095 | 0.531 |
| *NFE2* | 0.051 | 0.092 | 0.548 |
| *HEY2* | 0.051 | 0.485 | 0.104 |
| *GZMB* | 0.045 | 0.360 | 0.124 |
| *BCL2L1* | 0.042 | 0.068 | 0.609 |
| *NPAS2* | 0.039 | 0.065 | 0.591 |
| *TNFRSF10C* | 0.036 | 0.051 | 0.706 |
| *CDC25A* | 0.033 | 0.033 | 1.000 |
| *GADD45A* | 0.033 | 0.048 | 0.688 |
| *HGF* | 0.033 | 0.045 | 0.733 |
| *LGALS3* | 0.033 | 0.042 | 0.786 |
| *GCG* | 0.030 | 0.042 | 0.714 |
| *EPO* | 0.030 | 0.253 | 0.118 |
| *PIM1* | 0.027 | 0.027 | 1.000 |
| *EBAG9* | 0.027 | 0.062 | 0.429 |
| *SPP1* | 0.027 | 0.101 | 0.265 |
| *MMP1* | 0.024 | 0.140 | 0.170 |
| *HES1* | 0.021 | 0.065 | 0.318 |

**Table S16: Kidney core target genes.** There are forty-five "core" target genes which appear in every minimal target covering of the kidney samples. $P$(RNA) is the fraction of samples for which the indicated target gene is RNA-aberrant; $P$(RNA & upstream DNA) is the fraction of samples for which the indicated target gene is RNA-aberrant and there exists an DNA-aberrant gene among its sources. $P$(upstream DNA|RNA) is the fraction of the samples with the indicated gene RNA-aberrant for which at least one of its sources is DNA-aberrant.

| Target | $P$(RNA & upstream DNA) | $P$(RNA) | $P$(upstream DNA\|RNA) |
|---|---|---|---|
| *MSH2* | 0.492 | 0.567 | 0.868 |
| *STEAP3* | 0.469 | 0.725 | 0.648 |
| *IGFBP3* | 0.436 | 0.833 | 0.523 |
| *FANCD2* | 0.422 | 0.481 | 0.879 |
| *CXCL12* | 0.367 | 0.897 | 0.409 |
| *MDC1* | 0.342 | 0.417 | 0.820 |
| *PPARGC1A* | 0.314 | 0.414 | 0.758 |
| *BNIP3L* | 0.261 | 0.406 | 0.644 |
| *LCN2* | 0.258 | 0.383 | 0.674 |
| *KLK2* | 0.250 | 0.361 | 0.692 |
| *CA9* | 0.244 | 0.511 | 0.478 |
| *TGFA* | 0.206 | 0.481 | 0.428 |
| *MYC* | 0.178 | 0.206 | 0.865 |
| *BGLAP* | 0.178 | 0.239 | 0.744 |
| *HEY2* | 0.164 | 0.322 | 0.509 |
| *TP53I3* | 0.164 | 0.281 | 0.584 |
| *KLK3* | 0.131 | 0.194 | 0.671 |
| *PCBP4* | 0.128 | 0.236 | 0.541 |
| *HEYL* | 0.111 | 0.231 | 0.482 |
| *CASP6* | 0.106 | 0.164 | 0.644 |
| *SMAD6* | 0.106 | 0.181 | 0.585 |
| *TRIAP1* | 0.103 | 0.111 | 0.925 |
| *FABP4* | 0.094 | 0.200 | 0.472 |
| *SOD1* | 0.078 | 0.347 | 0.224 |
| *NOS2* | 0.067 | 0.153 | 0.436 |
| *CLDN5* | 0.042 | 0.058 | 0.714 |

**Table S17: Liver core target genes.** There are twenty-six “core” target genes which appear in every minimal target covering of the liver samples. $P$(RNA) is the fraction of samples for which the indicated target gene is RNA-aberrant; $P$(RNA & upstream DNA) is the fraction of samples for which the indicated target gene is RNA-aberrant and there exists an DNA-aberrant gene among its sources. $P$(upstream DNA|RNA) is the fraction of the samples with the indicated gene RNA-aberrant for which at least one of its sources is DNA-aberrant.

| Target | $P$(RNA & upstream DNA) | $P$(RNA) | $P$(upstream DNA\|RNA) |
|---|---|---|---|
| *CDC25C* | 0.744 | 0.858 | 0.867 |
| *CHEK1* | 0.682 | 0.766 | 0.890 |
| *IHH* | 0.660 | 0.933 | 0.707 |
| *MYBL2* | 0.641 | 0.787 | 0.814 |
| *TOP2A* | 0.641 | 0.923 | 0.695 |
| *VIM* | 0.626 | 0.798 | 0.784 |
| *SALL4* | 0.563 | 0.701 | 0.804 |
| *PTK6* | 0.501 | 0.744 | 0.673 |
| *PPARGC1A* | 0.467 | 0.514 | 0.908 |
| *ITCH* | 0.402 | 0.484 | 0.831 |
| *TNFRSF18* | 0.372 | 0.518 | 0.718 |
| *FGF2* | 0.368 | 0.458 | 0.803 |
| *AIFM2* | 0.359 | 0.376 | 0.954 |
| *BCL2L14* | 0.346 | 0.378 | 0.915 |
| *E2F7* | 0.344 | 0.351 | 0.982 |
| *TFF1* | 0.329 | 0.424 | 0.777 |
| *CEBPA* | 0.301 | 0.383 | 0.787 |
| *SOCS2* | 0.301 | 0.353 | 0.854 |
| *PPARG* | 0.228 | 0.383 | 0.596 |
| *RORC* | 0.222 | 0.232 | 0.954 |
| *PBX1* | 0.211 | 0.385 | 0.547 |
| *BCL2L11* | 0.153 | 0.163 | 0.934 |
| *ITGA5* | 0.144 | 0.342 | 0.421 |
| *NOS2* | 0.047 | 0.058 | 0.815 |
| *SERPINB13* | 0.026 | 0.028 | 0.923 |
| *ABCB4* | 0.024 | 0.325 | 0.073 |

**Table S18: Lung core target genes.** There are twenty-six "core" target genes which appear in every minimal target covering of the lung samples. $P(\text{RNA})$ is the fraction of samples for which the indicated target gene is RNA-aberrant; $P(\text{RNA}\,\&\,\text{upstream DNA})$ is the fraction of samples for which the indicated target gene is RNA-aberrant and there exists an DNA-aberrant gene among its sources. $P(\text{upstream DNA}|\text{RNA})$ is the fraction of the samples with the indicated gene RNA-aberrant for which at least one of its sources is DNA-aberrant.

| Target | $P$(RNA & upstream DNA) | $P$(RNA) | $P$(upstream DNA\|RNA) |
|---|---|---|---|
| *LGALS3* | 0.316 | 0.550 | 0.574 |
| *MYL9* | 0.293 | 0.381 | 0.770 |
| *EZH2* | 0.271 | 0.369 | 0.735 |
| *BNIP3L* | 0.246 | 0.462 | 0.533 |
| *TWIST1* | 0.224 | 0.654 | 0.343 |
| *PBX1* | 0.189 | 0.409 | 0.463 |
| *HIGD1A* | 0.181 | 0.210 | 0.864 |
| *CCNA2* | 0.171 | 0.253 | 0.677 |
| *CDC25C* | 0.155 | 0.171 | 0.905 |
| *HEY2* | 0.149 | 0.257 | 0.579 |
| *ITGBL1* | 0.134 | 0.189 | 0.710 |
| *PTGS2* | 0.132 | 0.159 | 0.833 |
| *PMS2* | 0.128 | 0.196 | 0.656 |
| *BAX* | 0.124 | 0.171 | 0.726 |
| *TP53I3* | 0.122 | 0.141 | 0.870 |
| *TOP2A* | 0.120 | 0.177 | 0.678 |
| *CDC25A* | 0.118 | 0.191 | 0.617 |
| *TIMP1* | 0.118 | 0.320 | 0.369 |
| *DHFR* | 0.112 | 0.163 | 0.688 |
| *BOLA2* | 0.110 | 0.365 | 0.302 |
| *SCO2* | 0.110 | 0.189 | 0.581 |
| *APP* | 0.108 | 0.269 | 0.402 |
| *MSH2* | 0.106 | 0.116 | 0.912 |
| *BCL2* | 0.102 | 0.175 | 0.581 |
| *ARID3A* | 0.100 | 0.106 | 0.942 |
| *TNFRSF10C* | 0.096 | 0.106 | 0.904 |
| *FOS* | 0.094 | 0.124 | 0.754 |
| *FANCC* | 0.092 | 0.110 | 0.833 |
| *RAG1* | 0.086 | 0.141 | 0.609 |
| *ZIC3* | 0.086 | 0.189 | 0.452 |
| *CDKN1A* | 0.084 | 0.086 | 0.976 |
| *RRM2B* | 0.075 | 0.100 | 0.755 |
| *FABP4* | 0.073 | 0.122 | 0.600 |
| *MAPKAPK5* | 0.073 | 0.104 | 0.706 |
| *CCNE1* | 0.071 | 0.098 | 0.729 |
| *BRCA1* | 0.071 | 0.077 | 0.921 |
| *HIF1A* | 0.065 | 0.132 | 0.492 |
| *PLK1* | 0.065 | 0.206 | 0.317 |
| *BCL6* | 0.063 | 0.081 | 0.775 |
| *TNFRSF18* | 0.057 | 0.112 | 0.509 |
| *ATR* | 0.057 | 0.096 | 0.596 |
| *HSP90AA1* | 0.057 | 0.130 | 0.438 |
| *CPT1A* | 0.055 | 0.100 | 0.551 |
| *RBL1* | 0.047 | 0.084 | 0.561 |
| *PKLR* | 0.039 | 0.092 | 0.422 |
| *VIM* | 0.039 | 0.077 | 0.500 |
| *IFNB1* | 0.039 | 0.071 | 0.543 |
| *MDM2* | 0.037 | 0.061 | 0.600 |
| *NDN* | 0.037 | 0.043 | 0.857 |
| *PTPN9* | 0.037 | 0.090 | 0.409 |

| | | | |
|---|---|---|---|
| *IL2RA* | 0.035 | 0.049 | 0.708 |
| *EBI3* | 0.035 | 0.045 | 0.773 |
| *NANOG* | 0.035 | 0.043 | 0.810 |
| *HMGCS1* | 0.035 | 0.059 | 0.586 |
| *PTK6* | 0.033 | 0.051 | 0.640 |
| *PLAGL1* | 0.031 | 0.033 | 0.938 |
| *GLI1* | 0.031 | 0.061 | 0.500 |
| *ITGAL* | 0.026 | 0.033 | 0.813 |
| *NEUROG3* | 0.026 | 0.035 | 0.765 |
| *CSN2* | 0.024 | 0.043 | 0.571 |
| *MLH1* | 0.022 | 0.024 | 0.917 |
| *CCNB2* | 0.022 | 0.051 | 0.440 |
| *CCND1* | 0.020 | 0.022 | 0.909 |
| *SOCS4* | 0.020 | 0.020 | 1.000 |
| *IGFBP3* | 0.020 | 0.057 | 0.357 |
| *UGT1A9* | 0.020 | 0.077 | 0.263 |

**Table S19: Prostate core target genes.** There are sixty-six "core" target genes which appear in every minimal target covering of the prostate samples. $P(\text{RNA})$ is the fraction of samples for which the indicated target gene is RNA-aberrant; $P(\text{RNA} \,\&\, \text{upstream DNA})$ is the fraction of samples for which the indicated target gene is RNA-aberrant and there exists an DNA-aberrant gene among its sources. $P(\text{upstream DNA}|\text{RNA})$ is the fraction of the samples with the indicated gene RNA-aberrant for which at least one of its sources is DNA-aberrant.

| | Luminal A | Luminal B | HER2-enriched | Basal-like |
|---|---|---|---|---|
| *CDC25A* | 0.228 | 0.694 | 0.800 | 0.943 |
| *CD36* | 0.228 | 0.512 | 0.800 | 0.770 |
| *S100B* | 0.685 | 0.760 | 0.982 | 0.402 |
| *MMP1* | 0.329 | 0.545 | 0.855 | 0.747 |
| *PLAGL1* | 0.370 | 0.669 | 0.582 | 0.172 |
| *CDC25C* | 0.516 | 0.901 | 0.964 | 0.977 |
| *E2F7* | 0.160 | 0.595 | 0.436 | 0.632 |
| *FABP4* | 0.260 | 0.570 | 0.745 | 0.667 |
| *EPO* | 0.132 | 0.289 | 0.564 | 0.115 |
| *FOS* | 0.352 | 0.645 | 0.818 | 0.713 |
| *BCL6* | 0.292 | 0.463 | 0.745 | 0.586 |
| *PTK6* | 0.142 | 0.223 | 0.527 | 0.126 |
| *BRCA1* | 0.269 | 0.686 | 0.418 | 0.540 |
| *FOXO1* | 0.269 | 0.570 | 0.673 | 0.494 |
| *IFNB1* | 0.119 | 0.223 | 0.309 | 0.517 |
| *MMP13* | 0.475 | 0.463 | 0.709 | 0.299 |
| *SOCS2* | 0.237 | 0.413 | 0.545 | 0.609 |
| *PIK3R1* | 0.311 | 0.603 | 0.527 | 0.690 |
| *CXCL12* | 0.406 | 0.570 | 0.764 | 0.701 |
| *S1PR1* | 0.242 | 0.504 | 0.545 | 0.540 |
| *LIFR* | 0.279 | 0.430 | 0.582 | 0.310 |
| *RHOU* | 0.406 | 0.529 | 0.527 | 0.253 |

**Table S20: Aberration probabilities of selected targets with sources in PAM50 sub-types.** Aberration probabilities for PAM50 sub-types for targets with source selected from the union of the target coverings; targets with source were required to have at least probability 0.4 in at least one sub-type.

| | CRIS-A | CRIS-B | CRIS-C | CRIS-D | CRIS-E |
|---|---|---|---|---|---|
| *TNFRSF10B* | 0.739 | 0.783 | 0.200 | 0.500 | 0.645 |
| *AXIN2* | 0.391 | 0.435 | 0.714 | 0.893 | 0.742 |
| *MYBL2* | 0.022 | 0.087 | 0.457 | 0.357 | 0.419 |
| *PDX1* | 0.848 | 0.826 | 0.600 | 0.464 | 0.677 |
| *SALL4* | 0.739 | 0.826 | 0.457 | 0.643 | 0.645 |
| *PERP* | 0.717 | 0.826 | 0.571 | 0.643 | 0.806 |

**Table S21: Aberration probabilities of selected targets with sources for colon tumor groups.** Targets with sources were selected by requiring a probability of 0.4 or more in at least one group.

| | G1 | G2 | G3 |
|---|---|---|---|
| *STEAP3* | 0.240 | 0.402 | 0.630 |
| *FANCD2* | 0.220 | 0.385 | 0.555 |
| *MSH2* | 0.340 | 0.425 | 0.622 |
| *MDC1* | 0.260 | 0.293 | 0.445 |
| *IGFBP3* | 0.300 | 0.483 | 0.437 |
| *PPARGC1A* | 0.220 | 0.282 | 0.395 |
| *LCN2* | 0.240 | 0.236 | 0.303 |
| *CXCL12* | 0.380 | 0.356 | 0.387 |

**Table S22: Aberration probabilities of selected targets with sources in liver tumor groups.** Targets with source were selected requiring a probability of 0.2 or more in at least one group.

| | Smoker | Recently Reformed | Reformed | Non Smoker |
|---|---|---|---|---|
| *AIFM2* | 0.465 | 0.371 | 0.367 | 0.074 |
| *CHEK1* | 0.814 | 0.774 | 0.483 | 0.556 |
| *BCL2L14* | 0.372 | 0.355 | 0.483 | 0.630 |
| *PPARGC1A* | 0.558 | 0.323 | 0.350 | 0.481 |
| *PTK6* | 0.535 | 0.274 | 0.417 | 0.444 |
| *E2F7* | 0.465 | 0.306 | 0.217 | 0.370 |
| *TFF1* | 0.395 | 0.403 | 0.317 | 0.185 |
| *CDC25C* | 0.814 | 0.774 | 0.667 | 0.593 |
| *MYBL2* | 0.698 | 0.677 | 0.600 | 0.481 |
| *FGF2* | 0.488 | 0.323 | 0.317 | 0.296 |
| *TOP2A* | 0.674 | 0.613 | 0.617 | 0.481 |
| *VIM* | 0.721 | 0.629 | 0.683 | 0.556 |
| *IHH* | 0.767 | 0.613 | 0.633 | 0.630 |
| *ITCH* | 0.488 | 0.435 | 0.367 | 0.333 |
| *TNFRSF18* | 0.465 | 0.339 | 0.333 | 0.333 |
| *SALL4* | 0.535 | 0.500 | 0.550 | 0.444 |

**Table S23: Aberration probabilities of selected targets with sources in lung tumor groups.** Targets with source were selected requiring a probability of 0.4 or more in at least one group.

| | 3+3 | 3+4 | 4+3 | 4+4 | >(4+4) |
|---|---|---|---|---|---|
| *EZH2* | 0.045 | 0.172 | 0.202 | 0.347 | 0.460 |
| *MYL9* | 0.068 | 0.221 | 0.263 | 0.449 | 0.410 |
| *CCNA2* | 0.045 | 0.041 | 0.111 | 0.184 | 0.367 |
| *CDC25C* | 0.000 | 0.055 | 0.141 | 0.143 | 0.324 |
| *BNIP3L* | 0.091 | 0.193 | 0.242 | 0.204 | 0.374 |
| *LGALS3* | 0.273 | 0.262 | 0.343 | 0.245 | 0.396 |
| *TWIST1* | 0.159 | 0.193 | 0.232 | 0.163 | 0.302 |

**Table S24: Aberration probabilities of selected targets with sources in prostate Gleason groups.** Targets with source were selected requiring a probability of 0.2 or more in at least one group.

| | 3 | 4 | 5 |
|---|---|---|---|
| *CCNA2* | 0.051 | 0.200 | 0.490 |
| *CDC25C* | 0.046 | 0.180 | 0.469 |
| *TOP2A* | 0.026 | 0.131 | 0.429 |
| *EZH2* | 0.148 | 0.310 | 0.551 |
| *BNIP3L* | 0.168 | 0.253 | 0.510 |
| *MYL9* | 0.189 | 0.343 | 0.449 |
| *CDC25A* | 0.092 | 0.098 | 0.306 |
| *PBX1* | 0.153 | 0.196 | 0.306 |
| *LGALS3* | 0.265 | 0.335 | 0.408 |

**Table S25: Aberration probabilities of selected targets with sources in prostate primary Gleason groups.** Targets with source were selected requiring a probability of 0.2 or more in at least one group.

| Subtype | Value | N | Entropy | Conf. Interval |
|---|---|---|---|---|
| | | **Breast** | | |
| All | | 953 | 17.02 | [16.63, 17.40] |
| PAM50 | Basal-like | 87 | 22.59 | [21.34, 23.69] |
| PAM50 | HER2-enriched | 55 | 18.95 | [17.52, 20.14] |
| PAM50 | Luminal A | 219 | 12.72 | [11.96, 13.38] |
| PAM50 | Luminal B | 121 | 17.43 | [16.45, 18.34] |
| ER Status | Negative | 167 | 22.39 | [21.36, 23.25] |
| ER Status | Positive | 569 | 15.51 | [15.04, 15.97] |
| Tumor Status | T1 | 249 | 13.56 | [12.88, 14.21] |
| Tumor Status | T2 | 551 | 18.25 | [17.72, 18.74] |
| Tumor Status | T3-T4 | 152 | 16.42 | [15.53, 17.26] |
| Lymph Node Status | Negative | 446 | 17.35 | [16.76, 17.89] |
| Lymph Node Status | Positive | 318 | 16.11 | [15.45, 16.75] |
| | | **Colon** | | |
| All | | 207 | 6.20 | [5.92, 6.46] |
| Stage | I-II | 113 | 6.52 | [6.13, 6.87] |
| Stage | III-IV | 85 | 5.63 | [5.22, 5.99] |
| Tumor Status | T2 | 28 | 5.72 | [5.01, 6.37] |
| Tumor Status | T3-T4 | 173 | 6.22 | [5.91, 6.51] |
| Lymph Node Status | Negative | 122 | 6.48 | [6.14, 6.83] |
| Lymph Node Status | Positive | 85 | 5.72 | [5.28, 6.13] |
| | | **Kidney** | | |
| All | | 336 | 8.25 | [7.86, 8.62] |
| Stage | I | 178 | 7.61 | [7.10, 8.09] |
| Stage | II-IV | 157 | 8.66 | [8.08, 9.22] |
| Tumor Status | T1 | 184 | 7.65 | [7.15, 8.14] |
| Tumor Status | T2-T4 | 152 | 8.60 | [7.98, 9.18] |
| | | **Liver** | | |
| All | | 360 | 10.93 | [10.45, 11.38] |
| Stage | I | 168 | 10.35 | [9.67, 10.90] |
| Stage | II | 85 | 11.01 | [10.10, 11.83] |
| Stage | III-IV | 81 | 10.12 | [9.26, 10.87] |
| Histology | G1 | 50 | 9.28 | [8.10, 10.30] |
| Histology | G2 | 174 | 10.85 | [10.23, 11.45] |
| Histology | G3-G4 | 131 | 10.68 | [9.94, 11.33] |

| Subtype | Value | N | Entropy | Conf. Interval |
|---|---|---|---|---|
| Tumor Status | T1 | 178 | 10.33 | [9.70, 10.90] |
| Tumor Status | T2 | 93 | 10.91 | [10.06, 11.69] |
| Tumor Status | T3-T4 | 86 | 10.77 | [9.87, 11.64] |
| | | **Lung** | | |
| All | | 465 | 11.65 | [11.30, 11.97] |
| Stage | I | 241 | 11.67 | [11.21, 12.09] |
| Stage | II | 110 | 11.57 | [10.96, 12.18] |
| Stage | III-IV | 91 | 10.67 | [9.94, 11.36] |
| Smoking history | Ancien | 87 | 9.71 | [8.95, 10.31] |
| Smoking history | Recent | 105 | 12.33 | [11.64, 12.97] |
| Tumor Status | T1 | 151 | 10.51 | [9.95, 10.98] |
| Tumor Status | T2 | 252 | 11.83 | [11.40, 12.25] |
| Tumor Status | T3-T4 | 60 | 12.50 | [11.70, 13.28] |
| Lymph Node Status | Negative | 296 | 11.91 | [11.46, 12.30] |
| Lymph Node Status | Positive | 158 | 11.03 | [10.51, 11.56] |

**Table S26: Entropy for source aberration with target across distinct tissues.** Entropy estimation (upper bound) on source aberration with target for several tissue types and tumor subtypes. $N$ is the total number of samples available in the given subtype.

| Subtype | Value | N | Entropy | Conf. Interval |
|---|---|---|---|---|
| | | **Breast** | | |
| All | | 953 | 39.03 | [38.72, 39.35] |
| PAM50 | Basal-like | 87 | 33.22 | [32.31, 33.97] |
| PAM50 | HER2-enriched | 55 | 35.05 | [33.97, 35.87] |
| PAM50 | Luminal A | 219 | 34.22 | [33.53, 34.88] |
| PAM50 | Luminal B | 121 | 38.46 | [37.72, 39.13] |
| ER Status | Negative | 167 | 37.11 | [36.39, 37.78] |
| ER Status | Positive | 569 | 38.41 | [38.01, 38.76] |
| Tumor Status | T1 | 249 | 36.31 | [35.69, 36.90] |
| Tumor Status | T2 | 551 | 39.88 | [39.49, 40.27] |
| Tumor Status | T3-T4 | 152 | 37.77 | [36.99, 38.50] |
| Lymph Node Status | Negative | 446 | 38.96 | [38.53, 39.34] |
| Lymph Node Status | Positive | 318 | 38.18 | [37.63, 38.69] |
| | | **Colon** | | |
| All | | 207 | 12.40 | [12.11, 12.68] |
| Stage | I-II | 113 | 12.07 | [11.65, 12.44] |
| Stage | III-IV | 85 | 12.28 | [11.80, 12.77] |
| Tumor Status | T2 | 28 | 11.26 | [10.47, 11.93] |
| Tumor Status | T3-T4 | 173 | 12.43 | [12.11, 12.74] |
| Lymph Node Status | Negative | 122 | 12.11 | [11.70, 12.53] |
| Lymph Node Status | Positive | 85 | 12.35 | [11.91, 12.76] |
| | | **Kidney** | | |
| All | | 336 | 20.88 | [20.19, 21.48] |
| Stage | I | 178 | 18.68 | [17.81, 19.44] |
| Stage | II-IV | 157 | 22.47 | [21.50, 23.32] |
| Tumor Status | T1 | 184 | 18.80 | [17.90, 19.62] |
| Tumor Status | T2-T4 | 152 | 22.33 | [21.36, 23.30] |
| | | **Liver** | | |
| All | | 360 | 26.05 | [25.58, 26.50] |
| Stage | I | 168 | 24.97 | [24.21, 25.58] |
| Stage | II | 85 | 25.59 | [24.62, 26.36] |
| Stage | III-IV | 81 | 25.65 | [24.65, 26.47] |
| Histology | G1 | 50 | 20.82 | [19.61, 21.86] |
| Histology | G2 | 174 | 25.27 | [24.60, 25.86] |
| Histology | G3-G4 | 131 | 26.76 | [26.01, 27.43] |

| Subtype | Value | N | Entropy | Conf. Interval |
|---|---|---|---|---|
| Tumor Status | T1 | 178 | 24.92 | [24.26, 25.56] |
| Tumor Status | T2 | 93 | 25.70 | [24.74, 26.48] |
| Tumor Status | T3-T4 | 86 | 26.16 | [25.22, 26.97] |

**Lung**

| | | | | |
|---|---|---|---|---|
| All | | 465 | 21.12 | [20.83, 21.42] |
| Stage | I | 241 | 21.17 | [20.73, 21.53] |
| Stage | II | 110 | 20.43 | [19.81, 20.94] |
| Stage | III-IV | 91 | 20.69 | [20.00, 21.31] |
| Smoking history | Ancien | 87 | 19.92 | [19.22, 20.53] |
| Smoking history | Recent | 105 | 20.94 | [20.32, 21.47] |
| Tumor Status | T1 | 151 | 20.39 | [19.86, 20.91] |
| Tumor Status | T2 | 252 | 21.25 | [20.83, 21.60] |
| Tumor Status | T3-T4 | 60 | 20.47 | [19.74, 21.07] |
| Lymph Node Status | Negative | 296 | 21.20 | [20.82, 21.55] |
| Lymph Node Status | Positive | 158 | 20.52 | [20.01, 21.02] |

**Prostate**

| | | | | |
|---|---|---|---|---|
| All | | 491 | 25.88 | [25.23, 26.47] |
| Gleason | 6 | 45 | 14.44 | [12.77, 15.68] |
| Gleason | 7 | 244 | 20.80 | [19.90, 21.61] |
| Gleason | 8 | 63 | 25.29 | [23.61, 26.69] |
| Gleason | 9 | 135 | 31.57 | [30.41, 32.53] |
| Primary | 3 | 196 | 17.53 | [16.63, 18.37] |
| Primary | 4 | 245 | 28.12 | [27.21, 29.02] |
| Primary | 5 | 49 | 32.13 | [30.25, 33.54] |
| Tumor Status | T2 | 186 | 19.72 | [18.86, 20.63] |
| Tumor Status | T3-T4 | 298 | 28.38 | [27.57, 29.12] |
| Lymph Node Status | Negative | 342 | 24.57 | [23.78, 25.27] |
| Lymph Node Status | Positive | 77 | 30.87 | [29.43, 32.18] |

**Table S27: Entropy for target aberration with source across distinct tissues.** Entropy estimation (upper bound) on target aberration with source for several tissue types and subtypes.

#### 1.7 Supplementary Figures

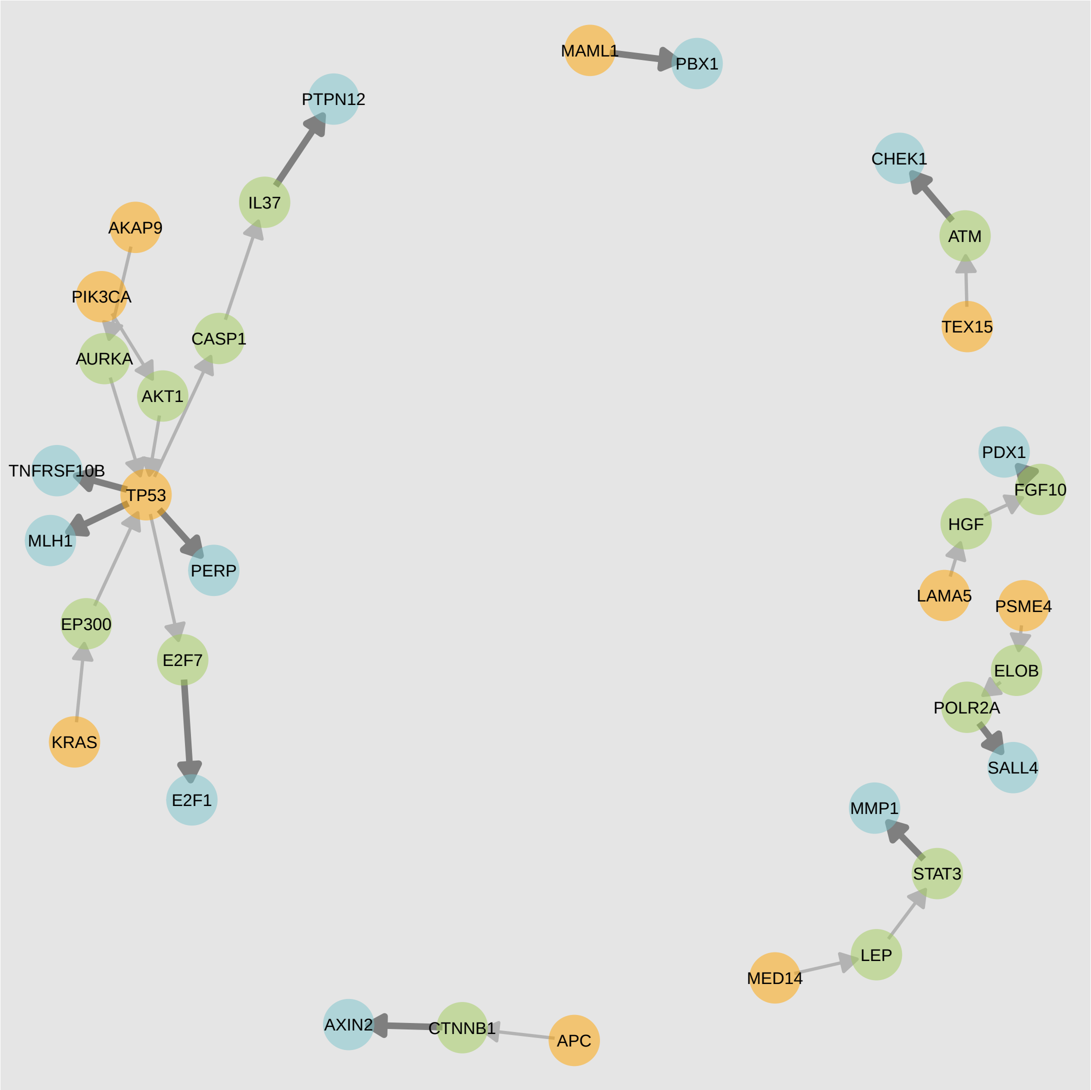

**Figure S1: Pair covering network for colon cancer.** The graphical network here shows a selected pair covering obtained for colon tumors; sources are shown in orange, targets in blue, and the intermediary genes in green. Strong connections are shown in bold edges. Many genes known to play an important role in colon tumorigenesis such as *TP53* and *KRAS* can be seen in the network.

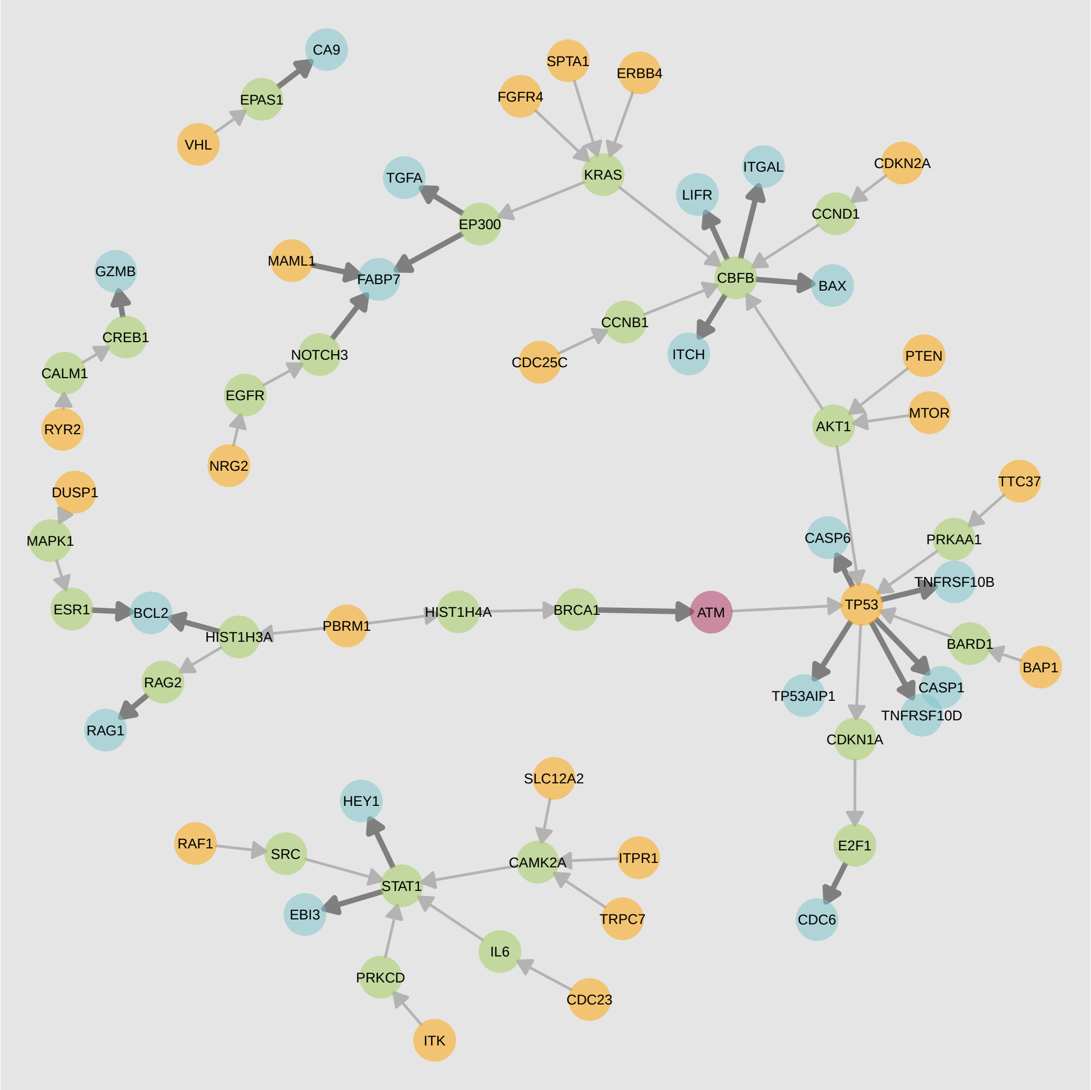

**Figure S2: Pair covering network for kidney cancer.** A pair covering obtained for kidney renal clear cell carcinoma (KIRC) is shown as a network with sources are shown in orange, targets in blue, and the intermediary genes in green. Genes that are both source and a targets, are shown in red. Strong connections are shown in bold edges.

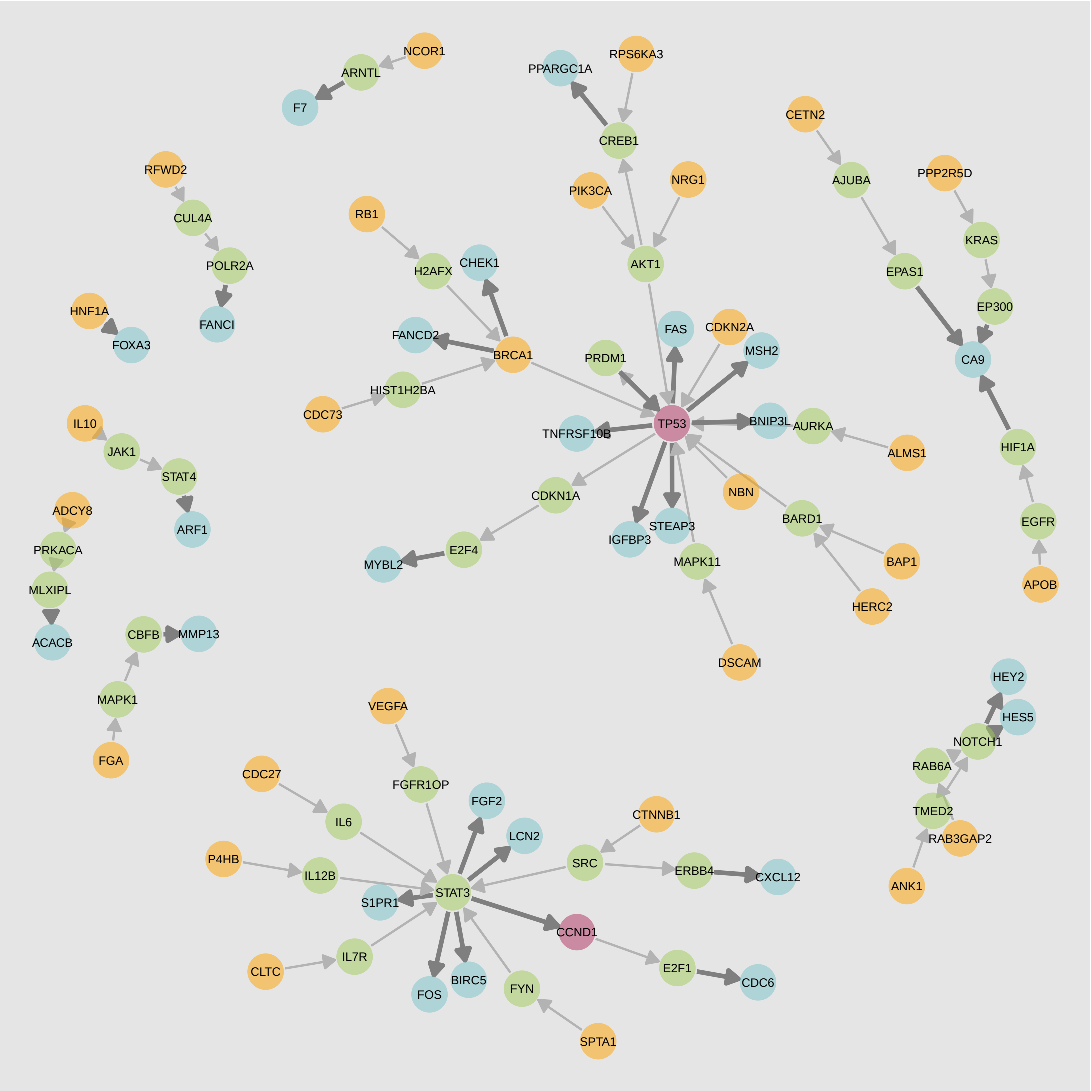

**Figure S3: Pair covering network for liver cancer.** A selected pair covering obtained for liver tumors are presented here as a graphical network; sources are shown in orange, targets in blue, and the intermediary genes in green. Genes that are both source and a targets, are shown in red. Strong connections leading to a target are presented as bolder edges.

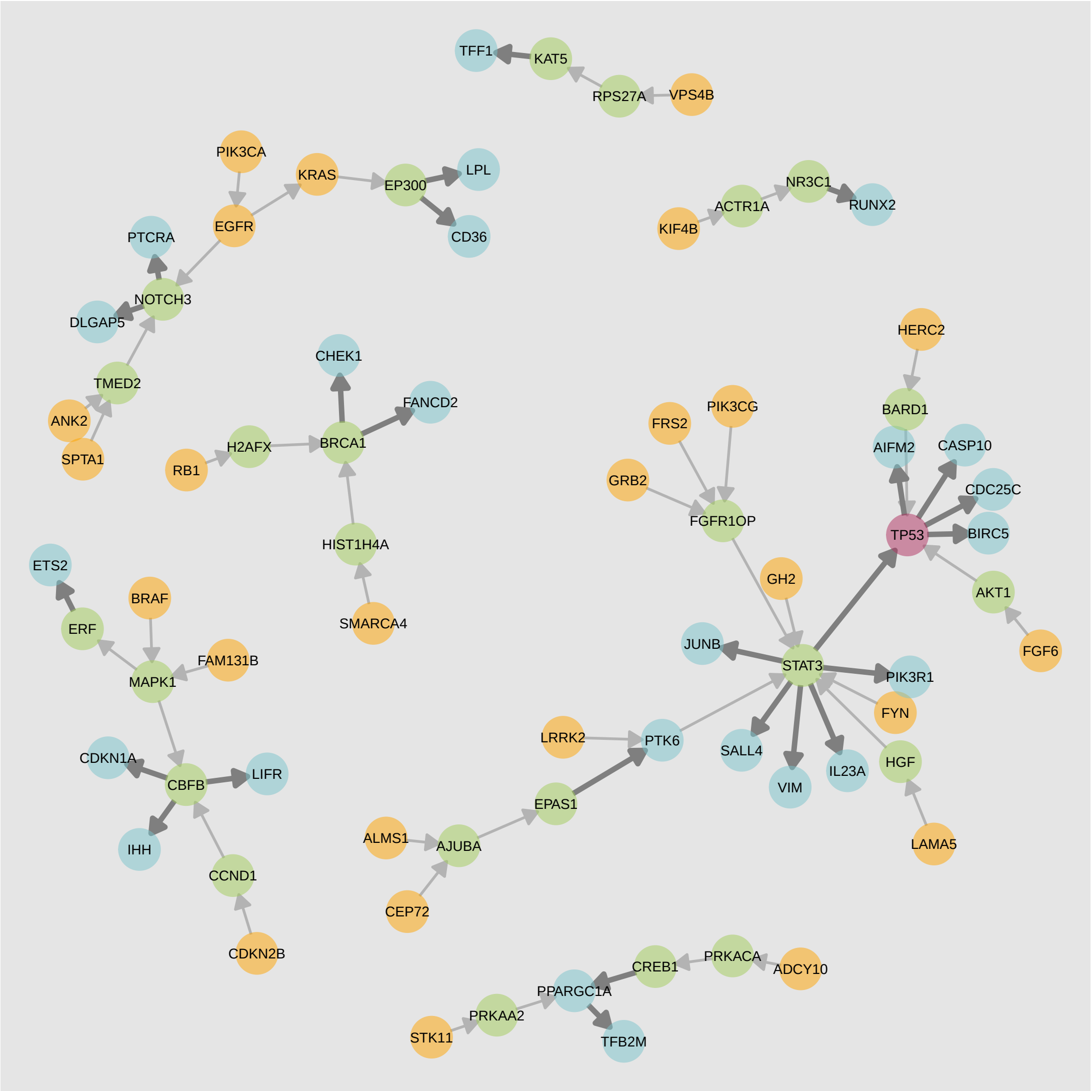

**Figure S4: Pair covering network for lung cancer.** A pair covering obtained for lung tumors is presented as a graphical network. Sources are shown in orange, targets in blue, and the intermediary genes in green. *TP53* which is both source and a target, is shown in red. *TP53, STAT3* and *NOTCH3* form important hubs. Many other known genes of importance such as *KRAS, EGFR* and *MAPK1* can be observed as well. Strong connections are shown in bold edges.

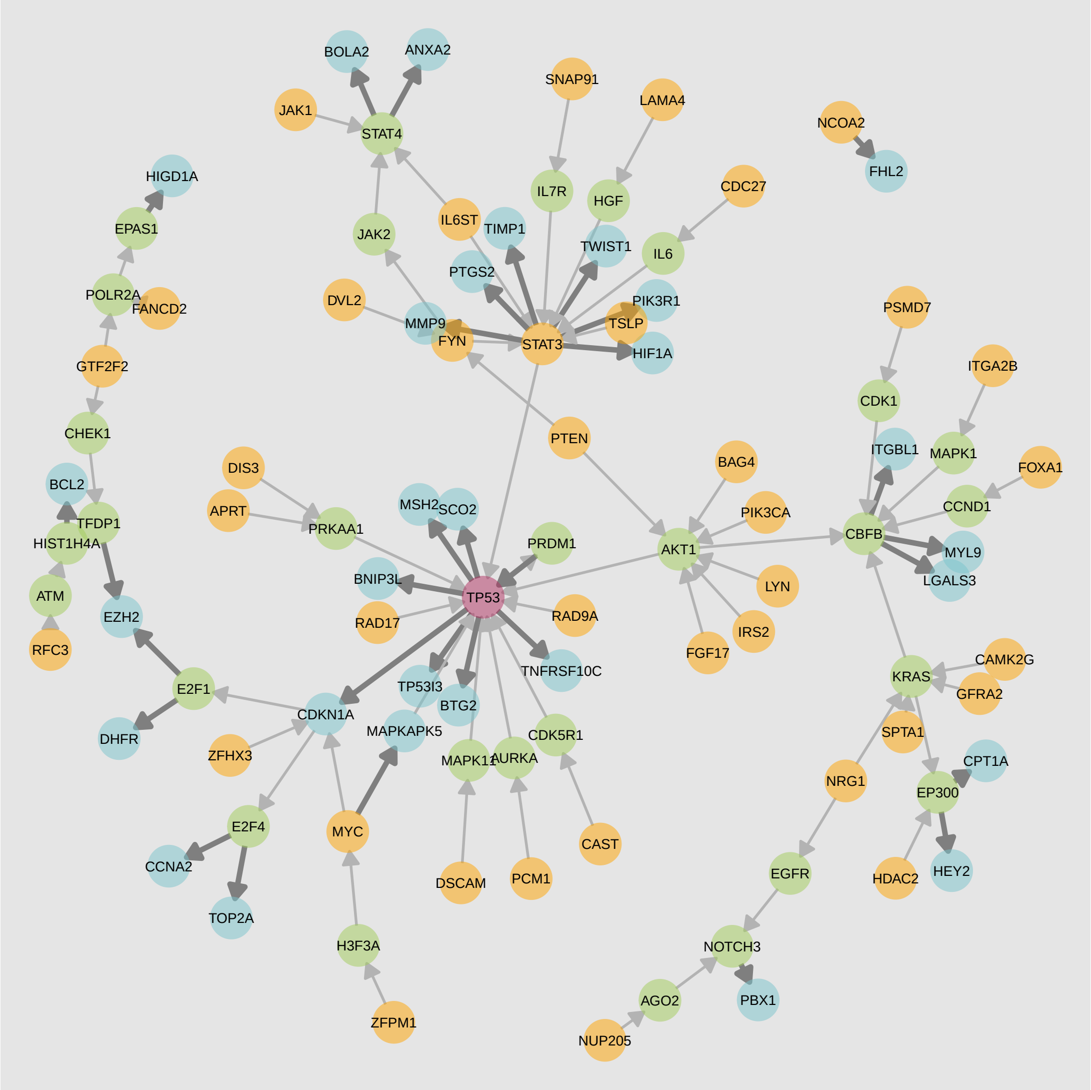

**Figure S5: Pair covering network for prostate cancer.** A selected pair covering obtained for prostate tumors are presented here as a graphical network; sources are shown in orange, targets in blue, and the intermediary genes in green. *TP53* which is both source and a target, is shown in red. Strong connections leading to a target are presented as bolder edges.

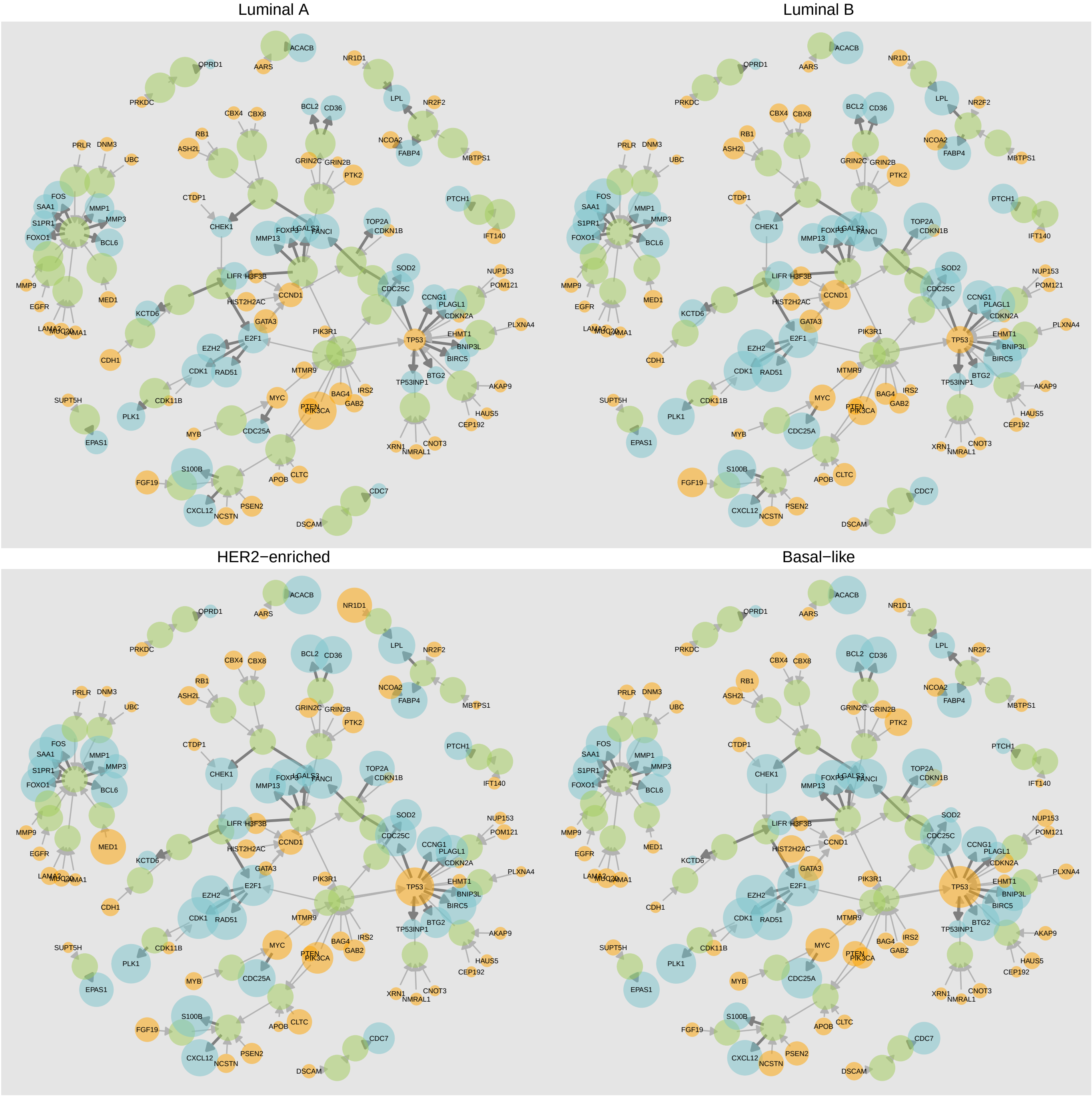

**Figure S6: Annotated networks for PAM50 breast cancer sub-types.** For four PAM50 sub-types considered (with the exception of the Normal-like sub-type) a breast covering presented as a graphical network with the size of nodes scaled to indicate source (with target) aberration probabilities and target (with source) aberration probabilities for samples in the given sub-type.

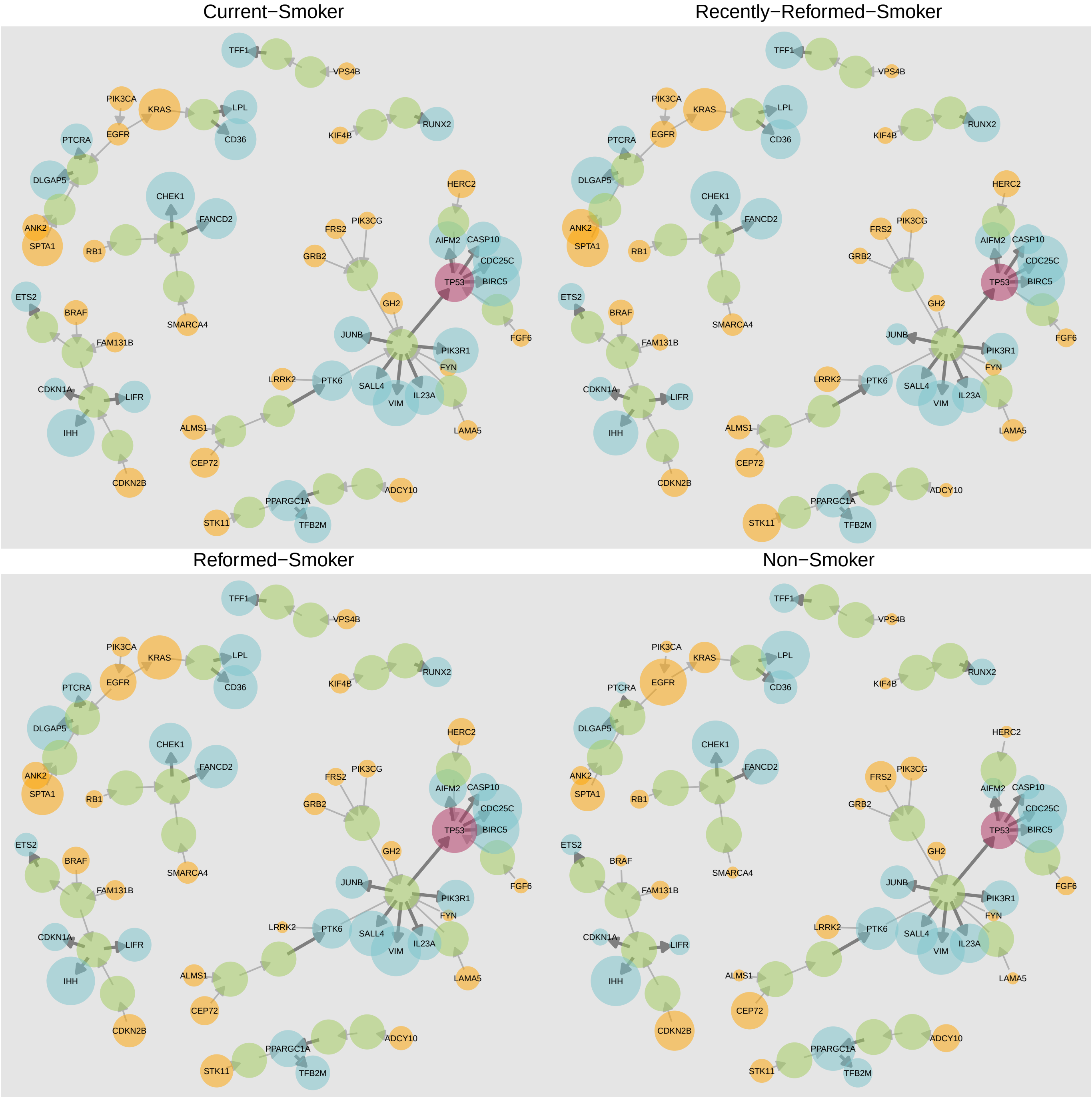

**Figure S7: Annotated networks for lung cancer based groups based on smoking history.** A lung covering is presented here with the size scaled to indicate source (with target) aberration probabilities and target (with source) aberration probabilities for samples in the given sub-type.

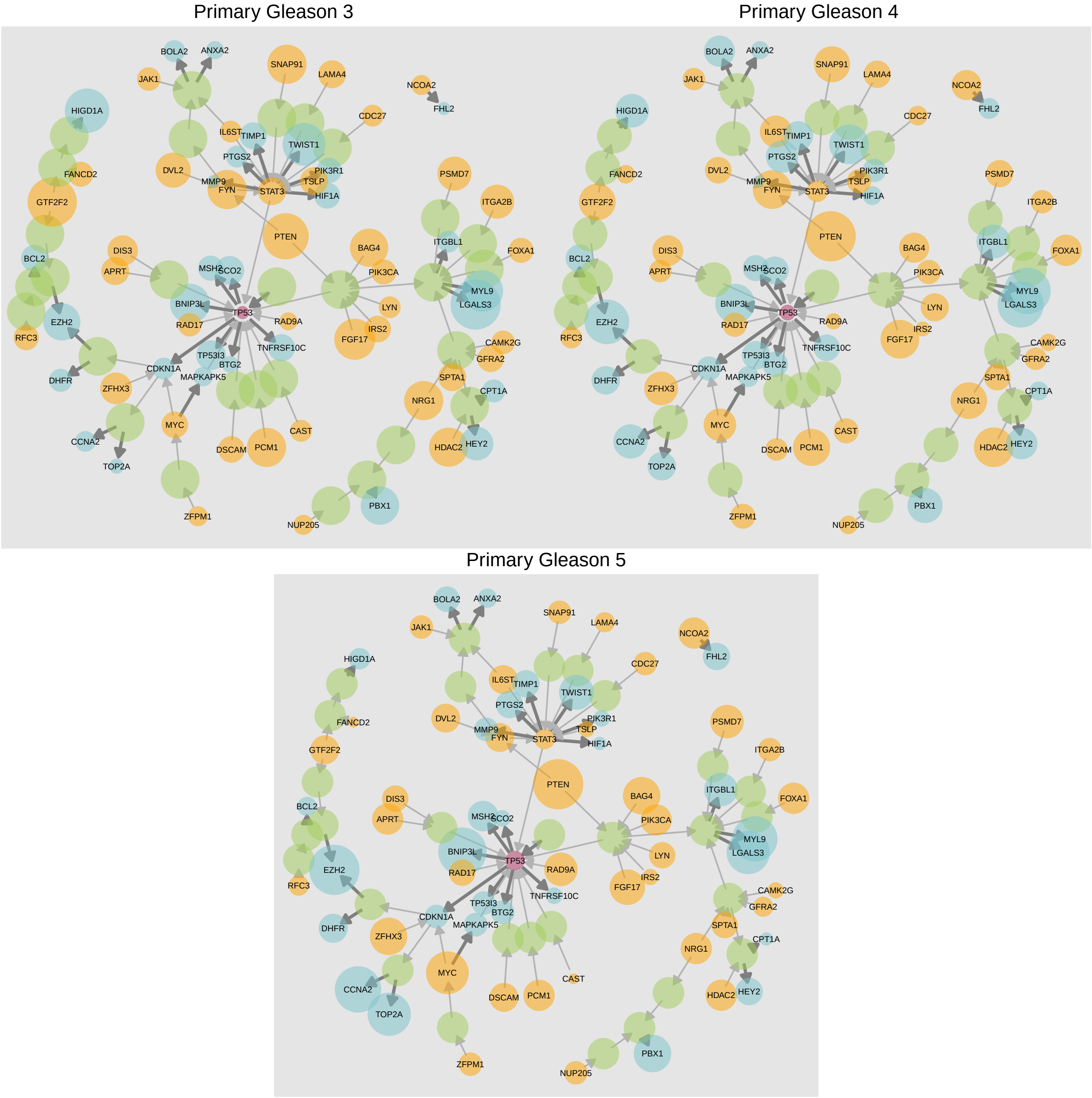

**Figure S8: Annotated networks for primary Gleason grade in prostate cancer.** A prostate covering is presented here with the size scaled to indicate source (with target) aberration probabilities and target (with source) aberration probabilities for samples in the given sub-type.

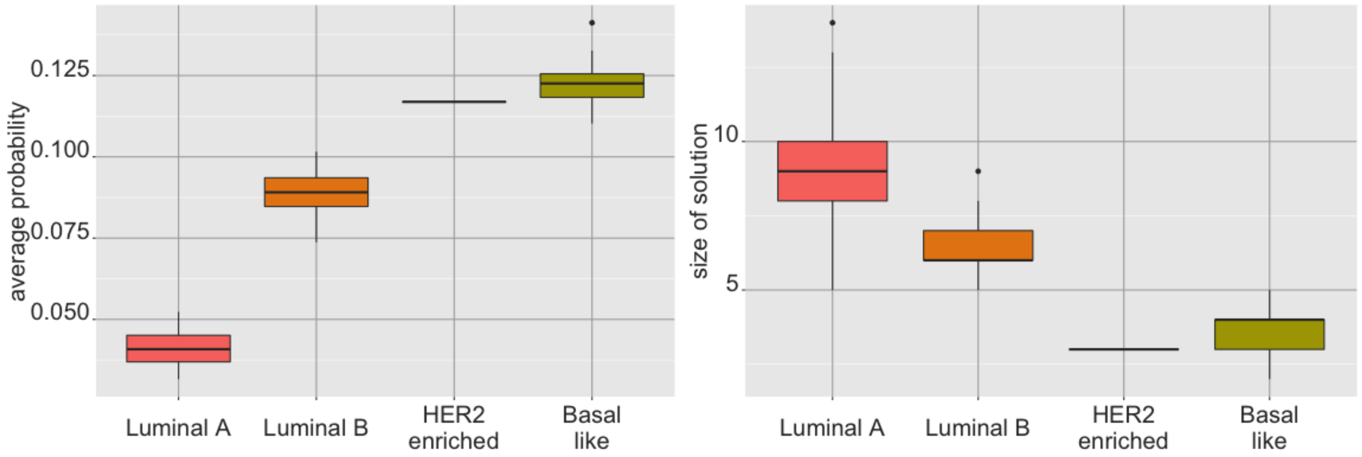

**Figure S9: Comparison of source sub-type coverings and pair probabilities across breast PAM50 classes.** For four PAM50 sub-types considered (with the exception of the Normal-like sub-type), an equal number of samples from each sub-type was randomly sampled. (Left) The average pair aberration probability over the breast pair covering union for 100 iterations of sampling; (Right) The average size of the source covering (computed over the breast source covering union with 90% or more coverage requested for the sub-type) for 100 iterations of such sampling. Between the two Luminal sub-types, the results indicate the more benign Luminal A group exhibiting lower aberration probabilities on average and larger covering sizes than the Luminal B group. A similar pattern is present between the Luminal sub-types and the more malignant Basal-like and HER2-enriched sub-types.

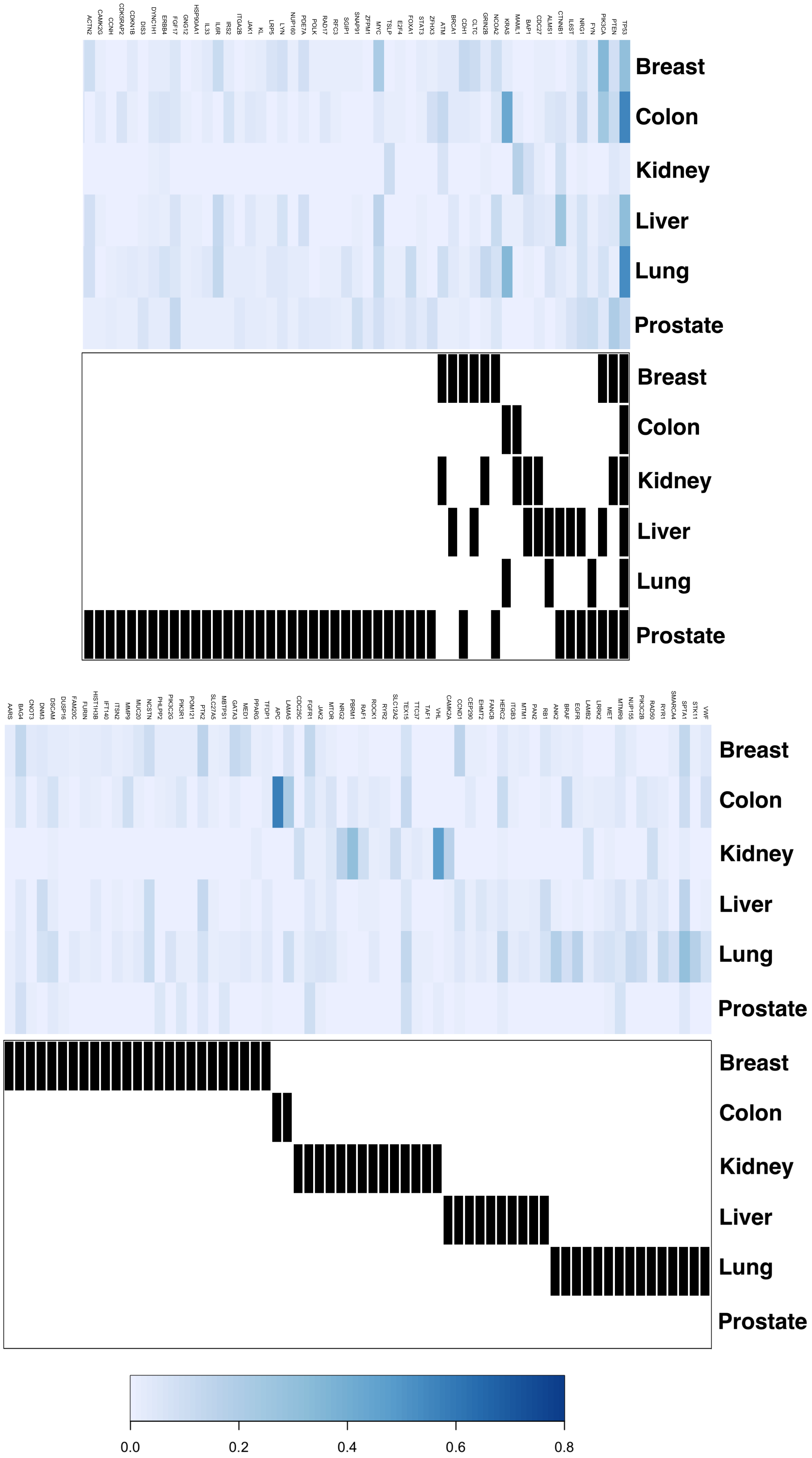

**Figure S10: Core set across tissues at the source level.** Complete core set across tissues at the source level (heatmap split for readability).

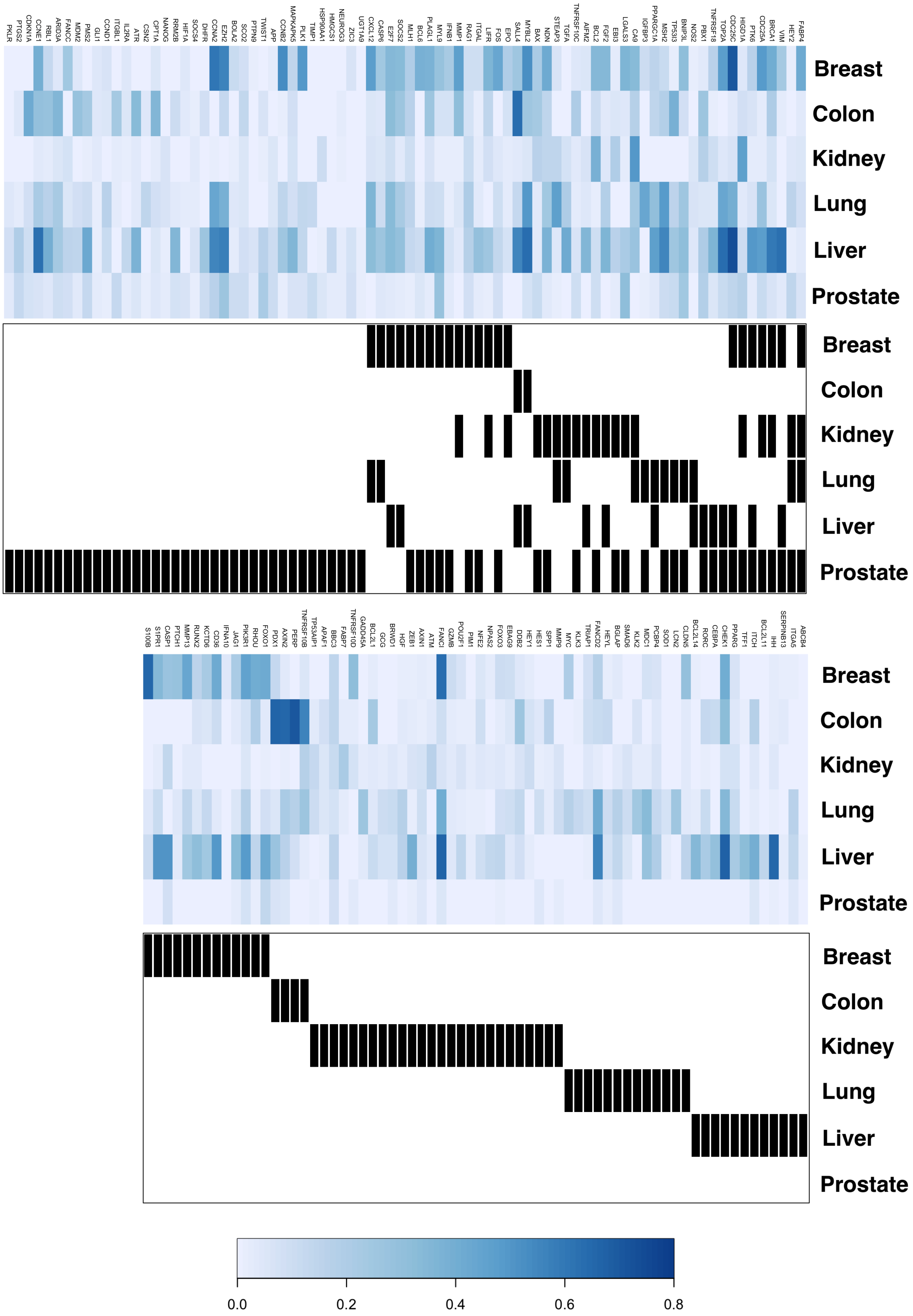

**Figure S11: Core set across tissues at the target level.** Complete core set across tissues at the target level (heatmap split for readability).

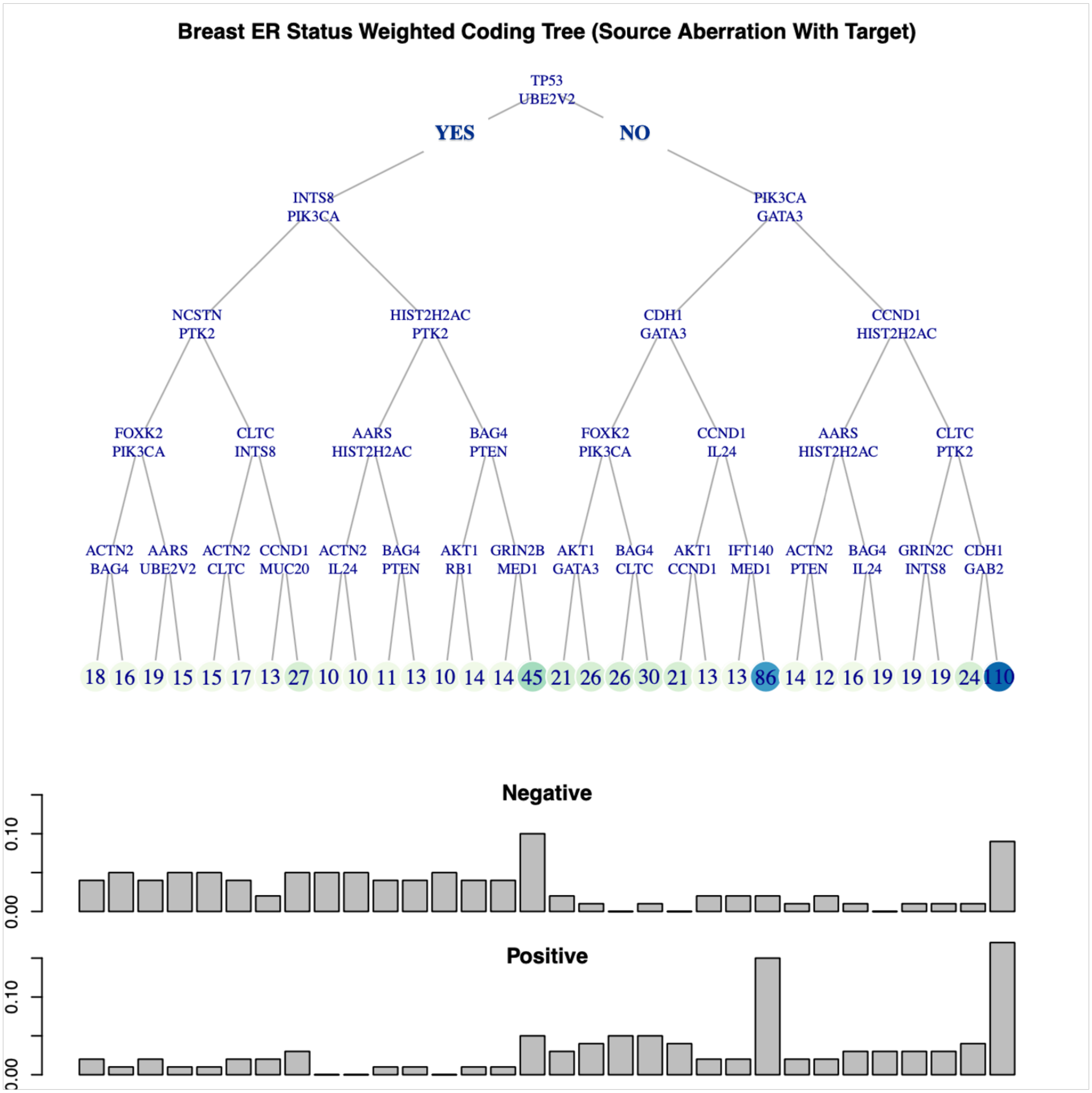

**Figure S12: Coding tree for breast ER status using source DNA aberration with target.** Coding tree for breast ER status using source DNA aberration with target.

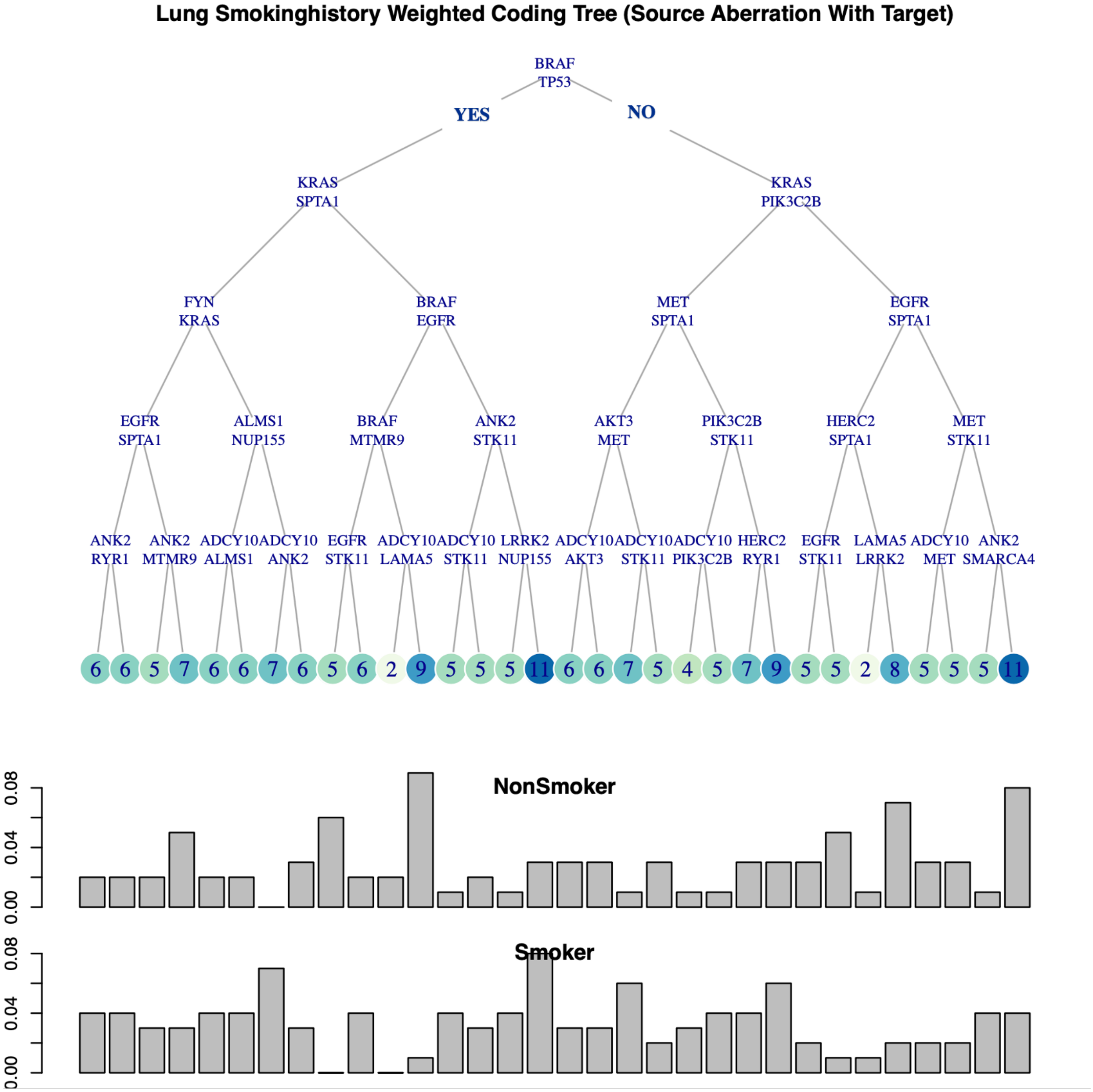

**Figure S13: Coding tree for lung smoking history using source DNA aberration with target.** Coding tree for lung smoking history using source DNA aberration with target.

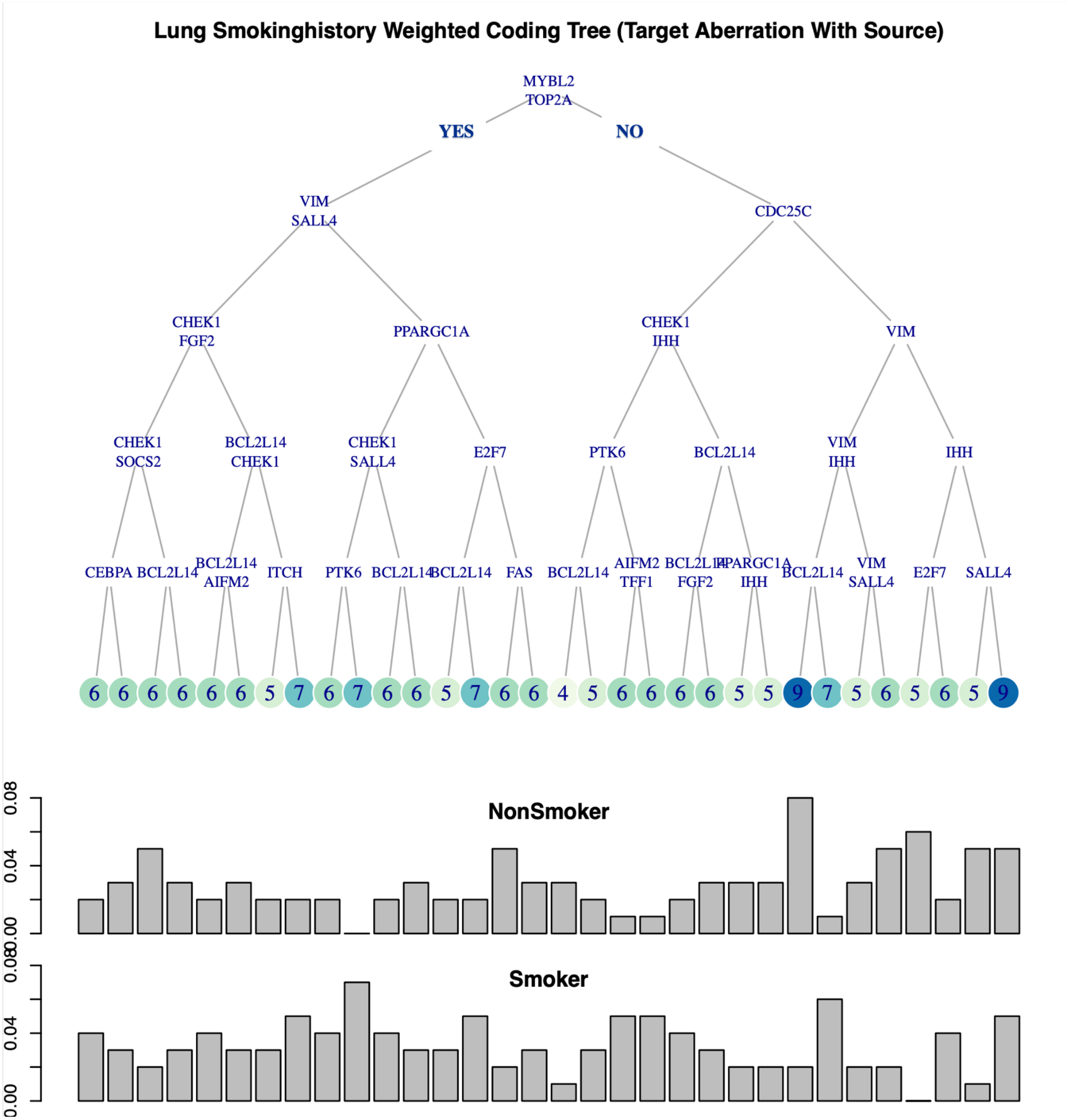

**Figure S14: Coding tree for lung smoking history using target RNA aberration with source.** Coding tree for lung smoking history using target RNA aberration with source.
